# The information architecture of the naturalistic human brain

**DOI:** 10.64898/2026.11.18.767548

**Authors:** George Blackburne, Alberto Liardi, Andreea Bacalum, Egor Levchenko, Greg Cooper, Fred Dick, Andrea I. Luppi, Jeremy I. Skipper, Pedro A. M. Mediano

## Abstract

How does the brain orchestrate its central currency, information, for complex adaptive cognition in the real world? Here we provide a comprehensive investigation into the spatiotemporal structure of qualitatively distinct modes of information dynamics in the human brain under ecological conditions. Applying a novel framework for information decomposition to fMRI data from individuals watching full-length feature films, we reveal a multidimensional large-scale information architecture, wherein different schemes for storing, transferring, and transforming representations occupy disjoint topo-graphic and network profiles. Further, this architecture reveals a computational division of labour, with certain regions acting as generalists that support multiple information-processing schemes and others specialising in a few. We show how these interactions reconfigure over time, coalescing into a structured cycle synchronised by sensory input, whose elements are tuned to semantic kinematics. Finally, we replicate these findings in an independent, large-scale dataset. Together, we provide a formal and empirical foundation for understanding the principles that govern distributed computation in the human brain during everyday life.

## I. INTRODUCTION

Fundamental to contemporary neuroscience is our conception of the brain as an information-processing system^1–3^. To make sense of human brain function, we must understand how it natively manipulates its central currency, information. However, information in a complex interacting system like the brain is a multi-faceted construct, consisting of several qualitatively different forms which may operate and interchange autonomously from each other^4–7^. Therefore, understanding how the brain uses information relies on our ability to go beyond simply asking how much information is present, towards interrogating what types of information dynamics are engaged^8^. Formal frameworks for information decomposition have revealed how information in multi-element systems occurs in three principal formats: redundant, synergistic, and unique information^5^. Redundant information is repeated over multiple elements, accessible from anywhere, and hence robust to the dysfunction of individual units. Synergistic information is contained within the whole but not in the parts, it is truly collective information which exceeds the sum of the constituting elements. Unique information, on the other hand, is information provided by one and only one particular element. Recent work has illustrated how the brain may exploit these formats of representation, for example, ensuring the fidelity of information for readout via redundancy^9,10^, or ensuring the parsimonious encoding of stimuli or distributed signals of surprise via synergy^11–14^.

For an adaptive system like the human brain, however, faced with a wealth of incessantly changing allostatic demands, it will not suffice to merely possess static representations, no matter how complex they are. Cognition, as commonly construed, is computation over representations^15^, entailing webs of input-output relations that reflect the storage, transfer, and transformation of information over time. In other words, it is essential to understand not just how an arbitrary bit of information is encoded, but how that specific bit changes its encoding from one moment to the next. For example, a bit of redundant information may stay redundant, but may also get erased from one element, such that it is unique in the future of the other. Alternatively, that bit of information may get “encrypted”, such that it can only be decoded by reading all parts simultaneously, and thus is encoded synergistically in the future of both. Fortunately, recent developments in information theory provide such a lens onto computation over representations, allowing the decomposition of any temporally symmetric or asymmetric relation of information between inputs and outputs in empirical data, yielding an encompassing map of the structure of computation^4^. Accordingly, it is now possible to ask: what is the large-scale organisation of information dynamics in the human brain?

The organisation of the human brain is often studied in the so-called “resting state”^16–20^, in which participants lie in the scanner without structured stimuli. Resting state has been valuable for neuroscience: it reduced reliance on subtractive task designs and simple activation contrasts, opened spontaneous, unconstrained cognition to study, and remains essential for populations or species for whom engaging with complex sensory demands is infeasible^21–23^. Yet a brain at rest is, by construction, a brain removed from the structured, ecological contexts in which it usually has to perceive, decide, act, and adapt. As such, there have been growing calls in human neuroimaging to employ integrated designs that situate the brain in those contexts directly^24^.

In this vein, naturalistic paradigms such as movie-watching have emerged. Feature-length films are long, continuous, contextualised, multisensory stimuli that engage individuals and mimic the sensory statistics of the real world^25^. They contain features that evoke responses at every stage of cognitive abstraction, with rapid perceptual content woven into more gradual narrative arcs, and thus demand in the scanner the similar information-processing to that everyday waking life demands of us. Importantly, unlike in resting state, where the cognitive operations that individuals are performing are unspecified, the contents of a film are known moment by moment, providing a reference for mapping how cortical computation entails cognition with constrained reverse inference^24^. Empirically, movie-watching have also been shown to out-perform rest for test-retest reliability^26^, revealing individual differences^27^, and brain-behaviour relationships^28^. In sum, movie-watching is a crucial complement to resting-state for the present aim of charting the large-scale organisation of the human brain in day-to-day existence^22^.

Here, we combine these threads to address the information architecture of the human brain under naturalistic conditions, applying a novel framework for information decomposition to functional magnetic resonance imaging (fMRI) data from 86 individuals watching full-length feature films. We first ask how prevalent the different modes of information dynamics are, where in the brain each is concentrated, and whether these topographies follow established divisions of the brain or reveal new axes of organisation. We then ask whether some regions specialise in a few modes while others engage the full repertoire, a distinction we capture with a new measure of computational diversity. Since computation is bounded by the connections that carry it, we ask how far each mode is supported by the axon bundles of the anatomical connectome, which strategy of signalling best explains it, and what kind of topologies they organise into. To shed light on the origin of individual differences in information-processing, we perform brain fingerprinting, and locate the idiosyncratic edges that drive individuality. Going beyond the time-averaged snapshot, we ask whether the brain’s information architecture is static or reconfigures from moment to moment, what kind of trajectory it expresses, and whether, and to what, in the world information dynamics are tuned to. Finally, we replicate all analyses in an independent cohort. Together, we comprehensively map the operation human brain in the real world, and open the door for a naturalistic neuroscience resolved in space, time, and information^31^.

## II. RESULTS

### Large-scale systems for information dynamics

We apply integrated information decomposition via information geometry (ΦID^IG^)^32^ (see Methods, Supplementary Methods) to fMRI data of individuals watching ten full-length feature films (*N* = 86: 500 Days of Summer, *n* = 20; Citizenfour, *n* = 18; and *n* = 6 each for The Usual Suspects, Pulp Fiction, The Shawshank Redemption, The Prestige, Back to the Future, Split, Little Miss Sunshine, and 12 Years a Slave), allowing us to precisely quantify how information remains or changes between redundant (*R*), synergistic (*S*), and unique (*U*) formats of representation across the human brain under ecological conditions.

To build intuition, we briefly describe each mode in turn. The simplest of these are modes in which information keeps its format over time. Redundant storage (*R*→*R*) denotes information that is redundant in the past and remains redundant in the future, information that the brain ensures is identically represented across both spatial and temporal coordinates. In synergistic storage (*S*→*S*), however, information is persistently encoded by both brain regions together, but neither one alone, residing in their joint state transcending individual contributions. Unique storage (*U*_*X*_ →*U*_*X*_), meanwhile, confines information to a single region, preserving it exclusively there. Information can also change location without changing format: information unique to one region in the past can migrate to become unique to the other in the future (*U*_*X*_→*U*_*Y*_). The remaining six modes convert information between formats, and come in time-reversed pairs: information unique to one region may be copied (*U*→ *R*), becoming redundant as it is replicated in the other, while redundant information may conversely be erased (*R* →*U*) from one region, leaving it unique to the other. Likewise, unique information may be unified by the brain over time into a synergistic state (*U* →*S*), or synergistic information may dissolve (*S*→ *U*), disintegrating from the whole to be held independently by a single region. Finally, information may be encrypted (*R* →*S*), transformed from the multilocalised format accessible format of redundancy either region into the delocalised format of synergy, or decrypted (*S* →*R*) along the reverse path ^4,32–34^.

We perform ΦID^IG^ for all pairs of 1000 cortical and 54 subcortical regions, and produce ten symmetric (1054 × 1054) matrices for each distinct mode of information dynamics (*R*→ *R, S* →*S, U*_*X*_ →*U*_*X*_ , *U*_*X*_ →*U*_*Y*_ , *U* →*R, R* →*U* , *U* →*S, S* →*U* , *R* →*S, S* →*R*) for each individual. Pooling across all interactions, it is clear that the brain hosts sharply different gross levels of each form of information dynamics (rm-ANOVA: 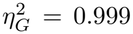, *p* = 2.40 × 10^−159^; pairwise comparisons: *p*_FDR_ = 1.30 10^−166^ to *p*_FDR_ = 1.51 ×10^−17^) (Fig. 1a). Specifically, we find that the brain is dominated by the unique storage of information (*U*_*X*_ → *U*_*X*_), while most impoverished in the unique transfer of information (*U*_*X*_ → *U*_*Y*_). Further, we find that the copy (*U* → *R*) and erasure (*R*→*U*) of information is richer than the unification (*U* →*S*) and dissolution (*S* →*U*) of information, which is in turn richer than the encryption (*R* →*S*) and decryption (*S* →*R*) of information. Importantly, we also find that the synergistic storage of information (*S S*) is higher than the redundant storage of information (*R* →*R*). Averaging across all interactions per region per mode of information dynamics, we observe a suite of novel functional topographies (Fig. 1b,c): *R*→ *R* shows the strongest effects in medial frontoparietal regions – such as the paracentral lobule and presupplementary motor area – as well as the dorsal thalamus. *S*→ *S* exhibits a similar pattern, but with pronounced effects in the parietal operculum and medial posterior prefrontal cortex (PFC). *U*_*X*_ →*U*_*X*_ is most prominent in the orbitofrontal and medial prefrontal cortices, as well as across the superior temporal gyrus and extrastriate regions. *U*_*X*_ →*U*_*Y*_ is strongly expressed across the subcortex, particularly within basal ganglia structures such as the putamen and globus pallidus. *U* →*R, R*→ *U* , *U* →*S*, and *S* →*U* all demonstrate dominance across the lateral dorsal prefrontal cortex, precuneus, and superior parietal lobule. *R* →*S* and *S* →*R*, by contrast, show the largest effects in the striate, extrastriate, and anterior insular regions. We note here that each mode of information dynamics has no mathematical mandate for distinction, and it is trivial to construct models that express exactly any or all^32^, and further, that these information dynamics do not simply reflect traditional functional connectivity (Supplementary Fig. S1).

**FIG. 1.**
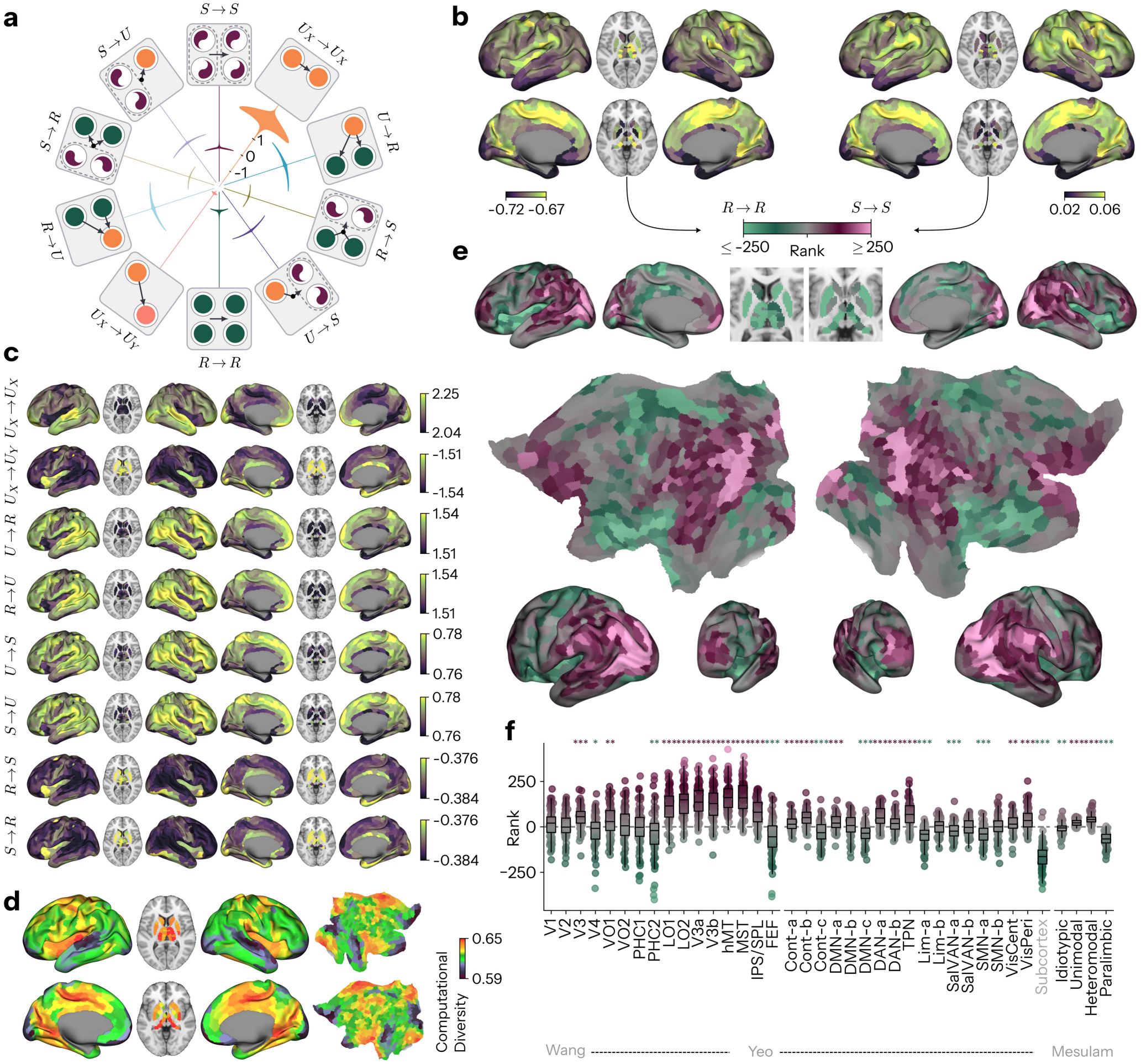
The information architecture of the naturalistic human brain. **a** Gross levels of different information dynamics in naturalistic human brain activity (bits) collapsed across individuals. **b** Group-average topographic patterns of regional (mean across interactions for each region) information storage via redundancy (*R→R*) and synergy (*S →S*) in bits. **c** Group-average topographic patterns of regional information dynamics for: unique storage (*U*_*X*_ *→U*_*X*_), unique transfer (*U*_*X*_*→ U*_*Y*_), copy (*U →R*), erasure (*R →U*), unification (*U→ S*), dissolution (*S→U*), encryption (*R→ S*), and decryption (*S→ R*) in bits (from top to bottom). **d** Group-average topographic patterns of regional (mean across interactions for each region) “computational diversity” (*D*) indexing the amount of distinct modes information dynamics engaged in an interaction (see Methods). **e** Gradient of ranked group-average *S → S* − *R →R* maps, the redundancy-to-synergy gradient of Luppi *et al*.^16^. **f** Redundancy-to-synergy gradient values for each subject subset by visual processing areas^29^, canonical resting-state networks (Yeo-Krienen-17)^30^, and canonical cytoarchitectonic classes (Mesulam)^3^ (points are participants). (*,**,*** indicates p<0.05,0.01,0.001 FDR-corrected Benjamini-Hochberg procedure. VO: ventral occipital areas; LO: lateral occipital; V3A, V3B: dorsal extrastriate visual areas; hMT: human middle temporal area (V5); MST: medial superior temporal area; HC: parahippocampal; IPS/SPL: pooled intraparietal and superior parietal fields; FEF: frontal eye fields; Cont: control; DMN: default mode network; DAN: dorsal attention network; TPN: temporoparietal network; Lim: limbic; Sal: salience; VAN: ventral attention network; SMN: somatomotor; Vis: visual.)

These maps expose both convergence and divergence in regional computation: some regions recur highly across many modes, whereas others are restricted to one or two. To capture this explicitly, we introduce computational diversity (*D*), quantifying the breadth of information dynamics engaged in a particular interaction, as the mean normalised participation ratio of each atom (see Methods, Supplementary Note 1, Supplementary Figs. S2 and S3) Averaged across interactions per region (Fig. 1d), we find that *D* is relatively strong across the brain (minimum mean 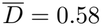). *D* is highest in the lateral dorsoanterior, dorsoposterior and medial ventroposterior thalamus, the insula, and medial parietal and occipital cortex, specifically the marginal extension of the central sulcus and retrosplenial cortex. These regions appear to not discriminate, agnostically engaging in the largest repertoire. On the other hand, we find *D* to be low in primary auditory and visual cortices, and thus they are the most skewed toward or focused on executing specific forms of information processing.

To understand the organisation of relative preference for the two antipodes of dynamical information sharing, pure synergistic storage (*S* →*S*) versus pure redundant storage (*R*→ *R*), we follow Luppi *et al*.^16^ by subtracting regional ranks of *R* →*R* from *S* →*S* to yield a “redundancy-to-synergy gradient” (Fig. 1e, see Methods). Fascinatingly, we find that during movie-watching, synergy rank dominance increases as we ascend the occipital lobe, most strongly in the dorsal and lateral directions, with further synergy-dominated patches in the precuneus, the supramarginal gyrus and the anterior end of the middle frontal gyrus. Conversely, the whole subcortex and most of the insular cortex are redundancy-dominated. To quantify how this organisation relates to existing stratifications of the cortex, for each individual’s redundancy-to-synergy gradient we average over regions assigned to the subdivisions prescribed by retinotopic visual processing areas^29^, resting-state functional connectivity networks^30^, and classical cytoarchitectonic areas^3^ (Fig. 1f). Among the restingstate networks, the visual, dorsal-attention, and temporoparietal networks show synergy-dominance during movie-watching, while the first limbic and somatomotor networks, and the entire subcortex were redundancy-dominated (all *p*_FDR_ *<* 0.01). Interestingly, synergy was most dominant in heteromodal but lowest in paralimbic cortex (both *p*_FDR_ *<* 10^−23^). Across the retinotopic hierarchy, as just mentioned, synergy dominates the later dorsal, lateral-occipital and motion-selective areas (V3A, V3B, LO1/2, hMT, MST; all *p*_FDR_ *<* 10^−18^), peaking in hMT and MST. This ascent is proper to the dorsal and lateral stream alone: equivalently downstream from V1 the ventral areas hV4 and PHC2 are instead redundancy- dominated (*p*_FDR_ = 0.043 and 0.005). Nor does the gradient climb past that peak: the pooled intraparietal and superior parietal fields remain synergy-dominated but fall short of hMT and MST (*p* = 6.1×10^−19^), while the frontal eye fields at the stream’s terminus revert to redundancy dominance outright (*p*_FDR_ = 1.1× 10^−6^; against hMT and MST, *p* = 8.1 ×10^−33^). In sum, during movie-watching the redundancy-to-synergy gradient resists symmetry across sensory systems, monotonic progression from “unimodal” to “heteromodal” cortex, and reduction to levels of laminar differentiation. Instead, we observe a novel limbic-to-extrastriate axis, running from redundancy-dominated subcortical and paralimbic territory to a synergistic peak in motion-selective cortex, that divides the dorsal from the ventral visual stream at every level and reverses at the dorsal stream’s frontal terminus.

Taken together, we discover that information dynamics in the human brain under ecological conditions depart from existing notions of large-scale brain organisation. Instead, they form a multidimensional organisational space of distributed computation occurring within and across brain systems.

### Networks of information dynamics

Each ensemble of region-by-region interactions can be cast as the adjacency matrix of a network^35,36^, allowing us to individually chart the structure of functional pathways created by specific distributed computations (Fig. 2).

**FIG. 2.**
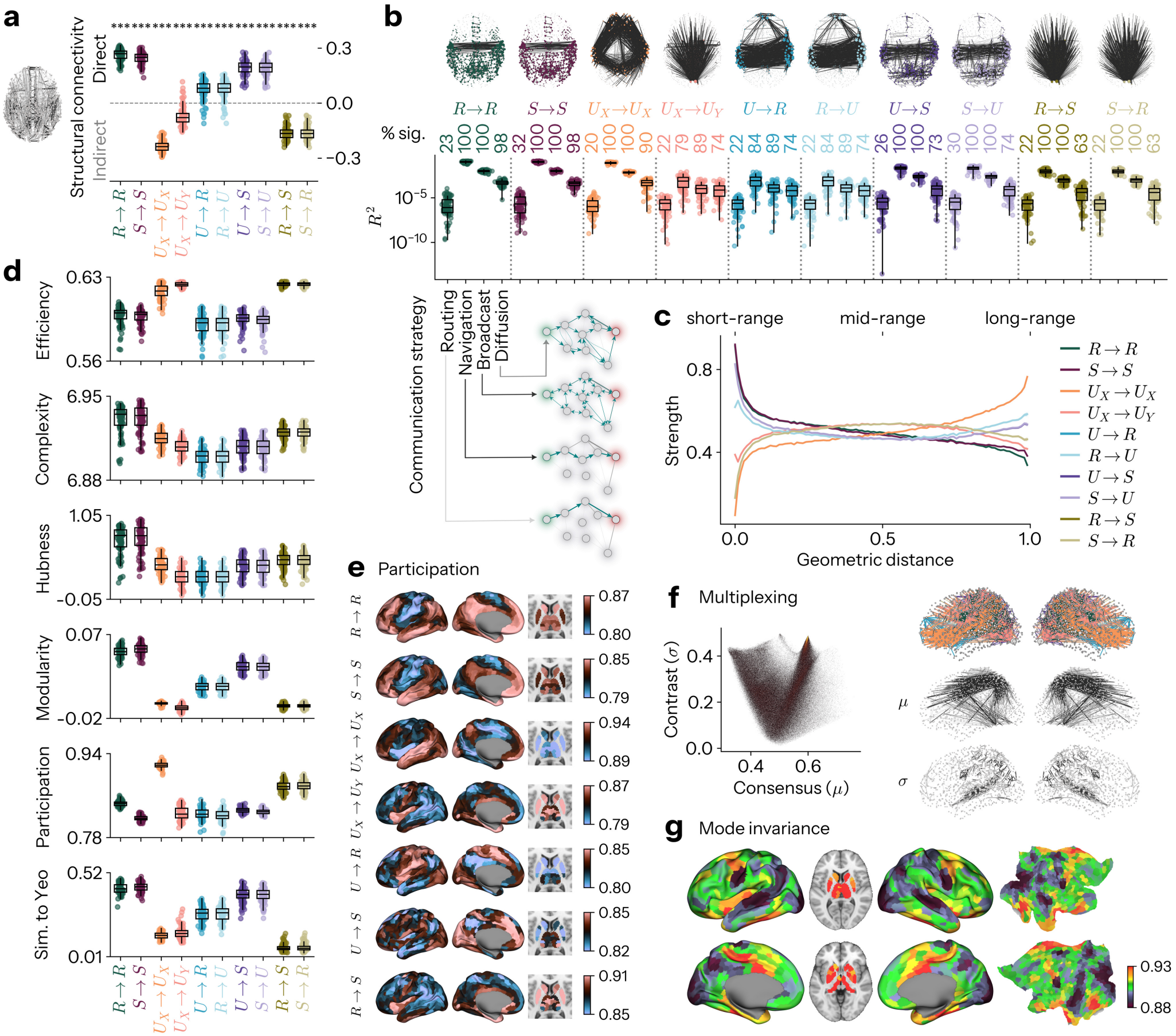
Network properties of information dynamics. **a** Relationship between information dynamics and types of structural connections. Rank-biserial correlation contrasts the atom’s edge weights on directly connected region pairs against those on structurally unconnected (polysynaptic) pairs. Points are participants and boxes their distribution; asterisks denote a one-sample *t*-test of the effect against zero. **b** Top: the 1000 strongest edges of each atom’s network. Prediction of networks of information dynamics from communication models applied to the structural connectome, implementing distinct strategies for polysynaptic signal transmission, ranging from centralised shortest pathway finding (Routing), to geometrically greedy iterative travel via spatially proximal nodes (Navigation), to simultaneous propagation among all available paths (Broadcast), to totally unbiased diffusion via random walks (Diffusion) (see Methods). Box plots show the variance in each atom’s edge weights explained (*R*^2^) by a per-participant OLS regression on the corresponding communication matrix, points denoting participants. *% sig*. is the percentage of participants whose *R*^2^ exceeds that of strength-preserving randomised surrogates of the communication matrix (*p*_perm_ *<* 0.05) (points are participants). **c** Relationship between information dynamics and scaled geometric (Euclidean) distance (mean *±* SEM). **d** Global topological properties of networks of information dynamics, from top to bottom: global efficiency (Efficiency), entropy of eigenmode contributions (Complexity), hierarchical heavy-tailed connectivity via the *s*-metric of scale-freeness (Hubness), separability of communities *Q* (Modularity), prevalence of mixed intermodular connectivity from the participation coefficient (Participation), and similarity to the Yeo-Krienen-17 networks via adjusted mutual information (Sim. to Yeo) (see Methods) (points are participants). **e** Topographic plots of nodal participation coefficient for modes of information dynamics. **f** Conceiving each network of information dynamics as a layer in a multilayer graph, we summarise each edge’s profile across layers by its mean *µ* (consensus) and standard deviation *σ* (contrast). Left: joint scatter density plot. Right, top to bottom: the top 500 edges per atom, the top 1000 consensus edges (*µ*), and the top 1000 contrast edges (*σ*). **g** Topographic plot of mode invariance: the global similarity of each region’s connectivity profile across all modes of information dynamics (see Methods). (*** indicates p<0.001 FDR-corrected Benjamini-Hochberg procedure.)

To understand how networks of information dynamics arise from signal propagation along networks of white matter bundles, first, we ask whether each mode of information dynamics is stronger between regions that are directly (perhaps monosynaptically) connected without passing through an intermediate grey matter node, or between those linked only indirectly, via multi-node polysynaptic pathways, using group-level structural connectomes from a separate cohort (see Methods, Fig. 2a). We find that *R*→ *R, S*→ *S, U* →*R, R* →*U* , *U*→ *S, S* →*U* all significantly colocalise with direct connections, with *R*→ *R* and *S* →*S* showing very strong evidence (*BF*_10_ =3.07 ×10^77^, 1.05 ×10^74^ respectively), whereas *U*_*X*_ →*U*_*X*_ , *U*_*X*_ →*U*_*Y*_ , *R*→ *S, S* →*R* are expressed most strongly between regions lacking a direct structural connection, with *U*_*X*_ →*U*_*X*_ being the strongest (*BF*_10_ =3.18 ×10^74^) (Overall: max *p*_FDR_=5.94×10^−21^, min *BF*_10_=8.07 ×10^17^). In other words, persistent redundancy and synergy are maximised, and unique information minimised, in the presence of immediate axonal support. Moreover, an intriguing dissociation appears whereby the conversion of information between delocalised and multilocalised formats occurs circuitously, but delocalised and singularly-localised formats occur directly.

To determine which principles of polysynaptic signalling might explain these networks of information dynamics, we tested how well a spectrum of communication models computed on the structural connectome could predict each of them (Fig. 2b, Supplementary Fig. S4, see Methods). These models range from centralised strategies, which rely on global knowledge of the network to coordinate signalling, to decentralised strategies, in which signals propagate using only local, node-level information. Ordered from centralised to decentralised, these spanned shortest-path finding (*Routing*), travel toward spatially proximal nodes (*Navigation*), simultaneous propagation along all walks (*Broadcast*), and unbiased random-walk diffusion (*Diffusion*)^37^. For each participant and model, we regressed the model’s communication matrix onto each atom’s edge weights to obtain a variance explained (*R*^2^), and assessed its significance against a null of strength-preserving surrogate networks^38,39^ (see Methods). The two extremes of the spectrum performed poorly. Averaged across atoms, routing was at chance 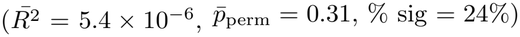, as expected given its need for global knowledge of network topology and correspond- ingly high informational cost, which makes it biologically implausible at scale. Unbiased diffusion was likewise only weakly predictive 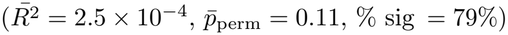, as expected from the long, gratuitous walks and delays it entails. On the other hand, the decentralised-yet- efficient strategies dominated. Across atoms, Navigation explained the most variance 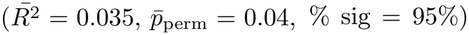, rising to 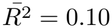 with 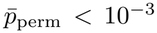 (the permutation floor of 1,000 surrogates) for *R*→ *R* and *S*→ *S* individually; Broadcast was significant in nearly every participant 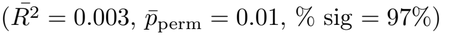, reaching % sig = 100% 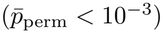 for *R* →*R, S*→ *S, U*_*X*_→ *U*_*X*_ , *U*→ *S, S* →*U* , *R* →*S* and *S*→ *R*. This ordering (Navigation *>* Broadcast *>* Diffusion *>* Routing) held for every atom, though the migration (*U*_*X*_ →*U*_*Y*_), copy (*U*→ *R*) and erasure (*R* →*U*) of information were poorly captured by any model 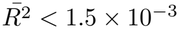 (Fig. 2b). We therefore provide preliminary evidence that large-scale neural computation may reflect both the geometrically greedy unfolding of signals following local spatial structure and the parallel poly-pathway broadcasting of signals across the connectome.

Consistent with these connectome- and communication- based results, the strength of each mode varied systematically with inter-regional Euclidean distance (Fig. 2c). Fitting each participant’s binned strength-by-distance profile with a quadratic function (strength ∼*β*_1_*x*+*β*_2_*x*^2^), the linear term recapitulated the direct/indirect streamline dissociation: persistent redundancy and synergy predominantly occur over short ranges (*R* →*R, β*_1_ = − 0.25; *S* →*S*, 0.23), whereas persistent unique information increases at long ranges (*U*_*X*_ →*U*_*X*_ , +0.29; all *p*_FDR_ *<* 10^−70^). The transformation of information is instead geometrically non-monotonic (curvature *β*_2_ significant throughout, *p*_FDR_ *<* 10^−7^): *U*_*X*_ →*U*_*Y*_ , *R*→ *S* and *S*→ *R* peak at intermediate distances (inverted-U), whereas *U* →*R, R*→ *U* , *U*→ *S*, and *S* →*U* are weakest there (U-shaped). Thus, each mode of computation occupies a characteristic spatial scale.

To gain insight into the overall organisation of each network of information dynamics, we systematically characterised their global topology (Fig. 2d,e, Supplementary Fig. S5). Contrasting each mode against the average of all others, *R*→ *R* and *S* →*S* formed the most complex (*p*_FDR_ = 3.0 ×10^−28^), hierarchical (*s*-metric; *p*_FDR_ = 2.8× 10^−32^) and modular (*p*_FDR_ = 7.2 ×10^−84^) architectures, most closely resembling the Yeo-Krienen functional systems (*p*_FDR_ = 7.2 ×10^−84^). Strikingly, the networks that transform between redundancy and synergy (*R*→ *S, S*→ *R*) revealed an entirely novel architecture, forming dramatically integrated (*p*_FDR_ = 6.1 ×10^−57^) and non-modular (*p*_FDR_ = 5.4× 10^−80^) networks that bear no resemblance to the Yeo-Krienen systems (*p*_FDR_ = 5.7 × 10^−67^). The preservation and migration of unique information (*U*_*X*_ → *U*_*X*_ , *U*_*X*_ → *U*_*Y*_) shared this distributed profile, with *U*_*X*_ → *U*_*X*_ forming the most participatory networks (*p*_FDR_ = 9.7 × 10^−89^). In between, the inter- conversion of unique and synergistic information (*U* → *S, S* → *U*) scored relatively moderately on every measure, while the copying and erasure of information (*U* →*R, R* → *U*) formed the least complex (*p*_FDR_ = 9.2 × 10 ^51^) and least efficient (*p*_FDR_ = 2.2 ×10^−43^) networks of all. Combined, these findings provide a topological lens on the division of computational labour in the brain under naturalistic conditions: persistent redundancy and synergy are maintained within canonical compartmentalised hierarchies, whereas the transformation between them is carried out in more efficient and egalitarian architectures, unlike known functional systems.

Given their concurrency, we can collect these different structures into a single multiplex network with multiple layers, where each layer represents a mode of information dynamics. Summarising this network by taking the mean (consensus *µ*) and standard deviation (contrast *σ*) across layers for each edge reveals that no connections are universally the strongest or weakest across all modes, with consensus being tightly constrained. Contrast, however, was near-maximal. Together, they formed a distinct V-shaped distribution where contrast dropped at moderate consensus levels but spiked at the extremes, meaning that the connections dominating the organisation of each layer were precisely those distinguishing the layer from other layers (Fig. 2f left). Examining the localisation of these quantities revealed a striking anatomical dissociation: a consensus network backbone linking distant frontotemporoparietal regions, and a contrasting periphery concentrated within sensory, subcortical and limbic systems (Fig. 2f right). Despite these edge-wise divergences, regional projection patterns were fairly conserved across modes (mode invariance, *v*; Fig. 2g). Invariance was high throughout 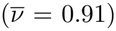, but strongest for subcortical regions such as the thalamus and the tail of the hippocampus, whereas primary visual, auditory and motor cortices displayed the lowest invariance (Fig. 2g).

In summary, we demonstrate how distinct modes of distributed computation in the naturalistic brain exhibit novel and distinct topological traits, but are nonetheless statistically explained by a shared set of polysynaptic communication policies.

### Subject-specific information dynamics

Having characterised the structural properties of networks of information dynamics in the human brain under naturalistic conditions, we next ask whether, and where, they come to constitute stable individual traits: reliable within a person, yet distinct between people (Fig. 3). Every group-average network reported above is representative of the individuals composing it (Supplementary Fig. S6).

**FIG. 3.**
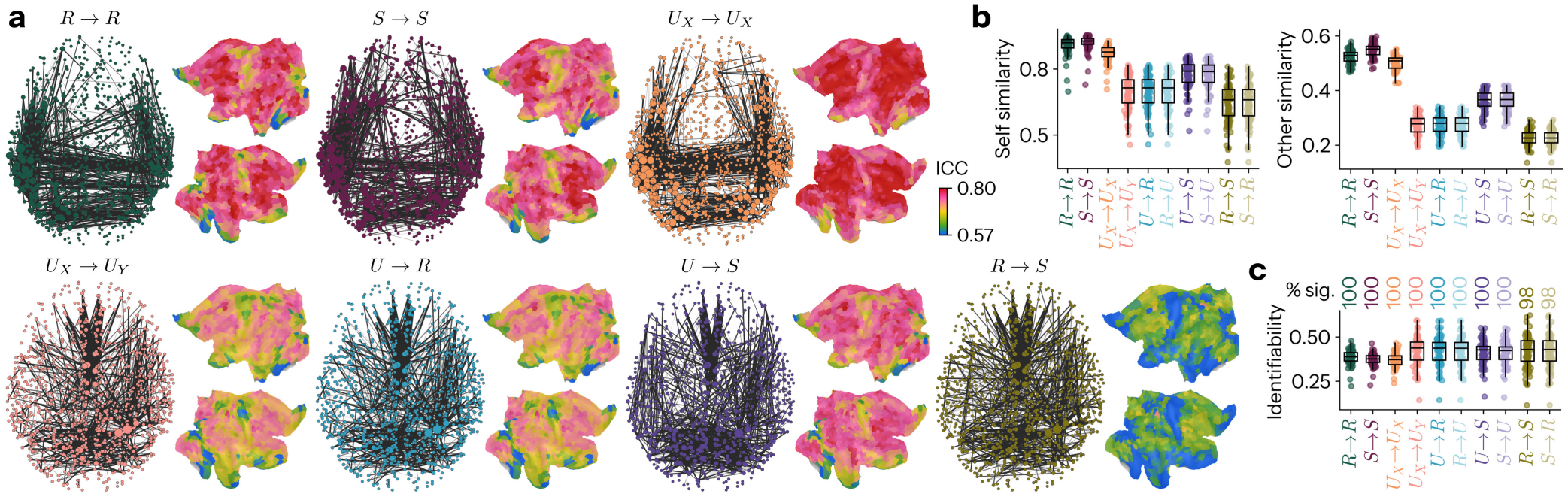
Individual differences in information dynamics. **a** Split-half reliability of each connection, via intra-class correlation (ICC) of its weight between the two halves of each individual’s scan; higher values denote connections that are reproducible within, and discriminative between, individuals. Brain networks show the 1000 most reliable edges; cortical flat maps show node-averaged ICC. **b** Fingerprinting of individuals from each mode of information dynamics. Top left: correlation between a mode’s connectivity in the first and second half of the same individual’s scan (Self similarity). Top right: correlation between one individual’s scan half and the matching half of every other individual (Other similarity). Bottom right: Self minus Other similarity (Identifiability), tested against a permutation null of shuffled subject labels (see Methods). (Points are participants)

We first quantified the reliability of each edge, for each mode of information dynamics, as the intra-class correlation (ICC) of its weight between the two halves of each participant’s scan (see Methods; Fig. 3a). Reliability varied markedly across edges (range −0.11 to 0.99) and modes (ANOVA: *F* =3.19×10^5^, p*<* 10^−300^; *U*_*X*_ → *U*_*X*_ > *R* → *R* > *S* → *S* > *U* → *S* > *S* → *U* > *R* → *U* > *U* → *R* > *U*_*X*_ → *U*_*Y*_ > *R* → *S* > *S* → *R*), and the most reliable edges of each mode occupied distinct territories. For the storage of information (*R* → *R, S* → *S, U*_*X*_ → *U*_*X*_), reliability concentrated across the lateral association cortex, linking the temporoparietal junction and the inferior and superior parietal lobules to lateral and dorsal prefrontal cortex. For the transfer and modification of information (*U*_*X*_ → *U*_*Y*_ , *U* → *R, R* → *U* , *R* → *S, S* → *R*), it instead concentrated along the cortical midline, among medial prefrontal cortex, the anterior and mid-cingulate, and the precuneus and posterior cingulate.

We then asked whether networks of information dynamics during movie-watching can be used to identify individuals via “brain fingerprinting” (see Methods; Fig. 3b). Every mode was strongly identifying, exceeding a permutation null of shuffled identities in at least 98% of participants (*p*_perm_ *<* 10^−3^, at the permutation floor of 1,000 permutations). The modes nonetheless fingerprinted in opposing ways. Modes of information storage (*R* → *R, S* → *S, U*_*X*_ → *U*_*X*_) were the most self-similar and reliable, but also the most similar between individuals. Modes of information transformation were instead the least selfconsistent yet the most idiosyncratic: for example, the interconversions of redundancy and synergy (*R* → *S, S* → *R*) were the least reproducible of all, but among the least shared, with the migration, copying and erasure of information behaving alike. Modes thus differed strongly in absolute similarity (self: *F* =942.019, 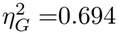 other: *F* = 4.37 × 10^3^, 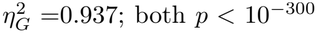) but only modestly in net identifiability (*F* =35.809, 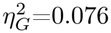, *p* = 5.54 × 10^−53^). In sum, information dynamics are not just an identifiable function of an individual, but show marked variability within an individual over the course of a scan.

### Time-resolved information dynamics

Recent work has increasingly emphasised that functional interactions in the brain are organised not just in space, but also in time^40,41^. Regions generate and dismantle statistical dependencies, with whole-brain patterns of cofluctuation tracing distinct sets of affiliations over the course of their observation^42,43^. However, as we have extolled here, brain dynamics do not simply evolve along an axis of engagement and disengagement, but in a multi-dimensional space of qualitatively distinct informational dependencies. Thus, to fully understand how computation unfolds, we develop a novel procedure to resolve each mode of information dynamics for each sample in time (see Methods).

We find that information dynamics exhibit complex and continuous reconfigurations over time (Fig. 4a). For example, generating the “redundancy-to-synergy gradient” of Fig. 1e but for each timepoint, we find that the static, time-averaged picture bears little resemblance to its time-resolved counterparts (r = 0.054 ± 0.096; Supplementary Fig. S7a). To quantify the intrinsic heterogeneity of each region’s fluctuations in computation, we estimate the state-space entropy rate^44^ of regional time courses of information dynamics (see Methods). Entropy rate here quantifies how much a region explores possible trajectories in a mode of information dynamics, and thus the inherent error associated with predicting its evolution. We reveal a spectrum of state-space exploration, in which the globus pallidus, parahippocampus, temporal pole, mid-cingulate, and insula possess the highest entropy rates, whereas the angular gyrus and precuneus possess the lowest. Thus, the human brain under naturalistic conditions possesses a hierarchy of prediction error strengths that progresses from parietal to paralimbic cortices (Fig. 4b).

**FIG. 4.**
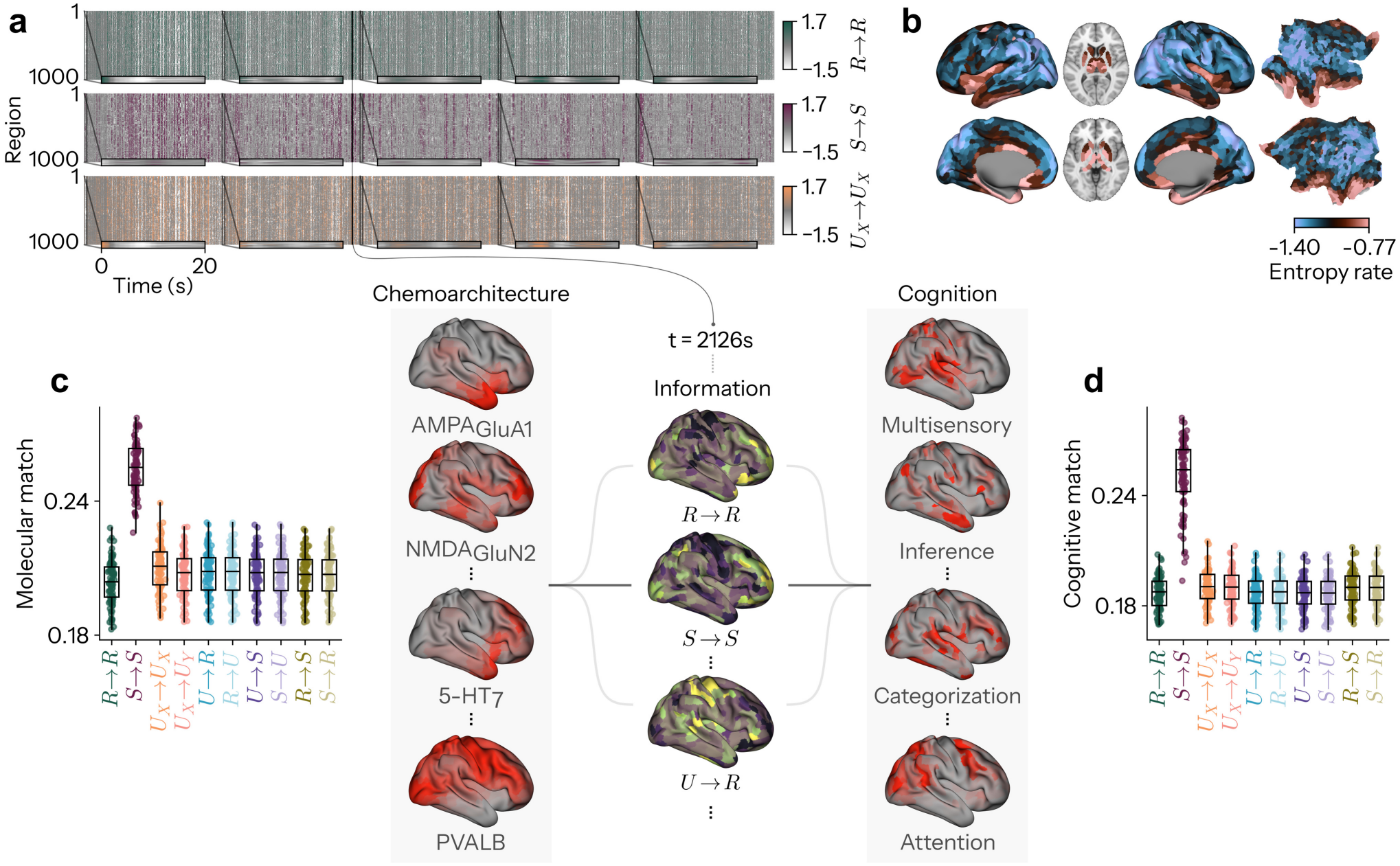
Chemoarchitectonic and cognitive matching of time-resolved information dynamics. **a** Time-resolved fluctuations in information storage (Top: *R → R*, Middle: *S → S*, Bottom: *U*_*X*_ *→ U*_*X*_), averaged over interactions for each region for an example participant (Sub-1 5469 total seconds). **b** Topography of temporal prediction error, estimated via mean state-space entropy rate of regional information-dynamics time courses (nats). Lower panel: At each timepoint, the whole-brain map of each mode of information dynamics is compared against reference maps of brain chemoarchitecture (left; AHBA mRNA expression maps for >1,900 brain-specific genes) and meta-analytic cognitive function (right; NeuroSynth meta-analytic term maps from over >14,000 neuroimaging experiments) and averaged across time, measuring the extent to which each mode of information dynamics mirrors molecular or cognitive patterns, respectively (see Methods). **c** Molecular match: correspondence between each mode’s time-resolved maps and the chemoarchitectural maps (points are participants). **d** Cognitive match: correspondence between each mode’s time-resolved maps and the cognitive maps (points are participants).

To assess the extent to which these unfolding whole-brain patterns of information dynamics can be cashed out in terms of lower-level biophysical detail and higher-level cognitive function, we assayed their spatial correspondence to two corpora from different cohorts, reflecting the brain’s chemoarchitecture and its cognitive architecture (Fig. 4c,d; see Methods). Across modes, *S* → *S* stood out as by far the most strongly matched to both corpora (molecular match: max *p*_FDR_ = 5.40 × 10^−56^; cognitive match: max *p*_FDR_ = 4.96 × 10^−44^; Fig. 4c,d; Supplementary Figs. S9 and S10), an effect that persists for two alternative neuron- and synapse-specific gene sets (Supplementary Fig. S8). To discern the relative allegiance to structural (chemoarchitectonic) versus functional (cognitive) maps, we took the normalised ratio of the two, dividing the cognitive match by the sum of the molecular and cognitive matches, and found *S* → *S* to be the most functionally biased of all modes (max *p*_FDR_ = 1.62 × 10^−16^). Thus, distributed information dynamics play out over time in non-trivial topographies differentially anchored by molecular and cognitive templates.

### Cycles of global information dynamics

Inspection of the time courses of information dynamics highlighted repeated moments where particular modes featured globally prominent, showing widespread peaks across the brain (Fig. 4a). To capture this quantitatively, we collapsed the multidimensional information dynamics at each timepoint onto the single mode with the relative strongest global expression, yielding for each participant a discrete winner-take-all sequence of *global information states* (see Methods; Fig. 5a). Every mode uniquely peaked above all others at some point, but they did so with grossly different temporal statistics (rm-ANOVA: dwell time: *F* (9, 765) = 615.55, 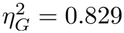 appearance rate (Freq): *F* (9, 765) = 1.32 ×10^3^, 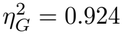 fractional occupancy (FO): *F* (9, 765) = 934.12, 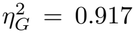 all *p <* 10^−300^) (Fig. 5b). Specifically, we find that the preservation of unique information (*U*_*X*_ → *U*_*X*_) has the highest FO (19.1±3.2%; max *p*_FDR_ = 3.20×10^−5^), driven by its long mean dwell time (3.35 ± 0.35 s; max *p*_FDR_ = 5.64 × 10^−35^) rather than its appearance rate, reflecting long spans of global functional differentiation. *U* → *R* and *R* → *U* also possess large FO’s, but this is driven by having the highest appearance rates (max *p*_FDR_ = 2.19×10^−28^), meaning frequent bursts of copy and erasure of information. Interestingly, we found that *U*_*X*_ → *U*_*Y*_ very rarely occurs (1.3 ± 0.4%; max *p*_FDR_ = 1.86 × 10^−45^), and when it does so, it is fleeting (1.49 ± 0.14 s; max *p*_FDR_ = 1.16 × 10^−41^), before yielding to another mode. Finally, *S* → *S* has a higher dwell time and frequency of occurrence than *R* → *R* (*p*_FDR_ = 8.85 × 10^−15^ and *p*_FDR_ = 2.14 × 10^−38^, respectively).

**FIG. 5.**
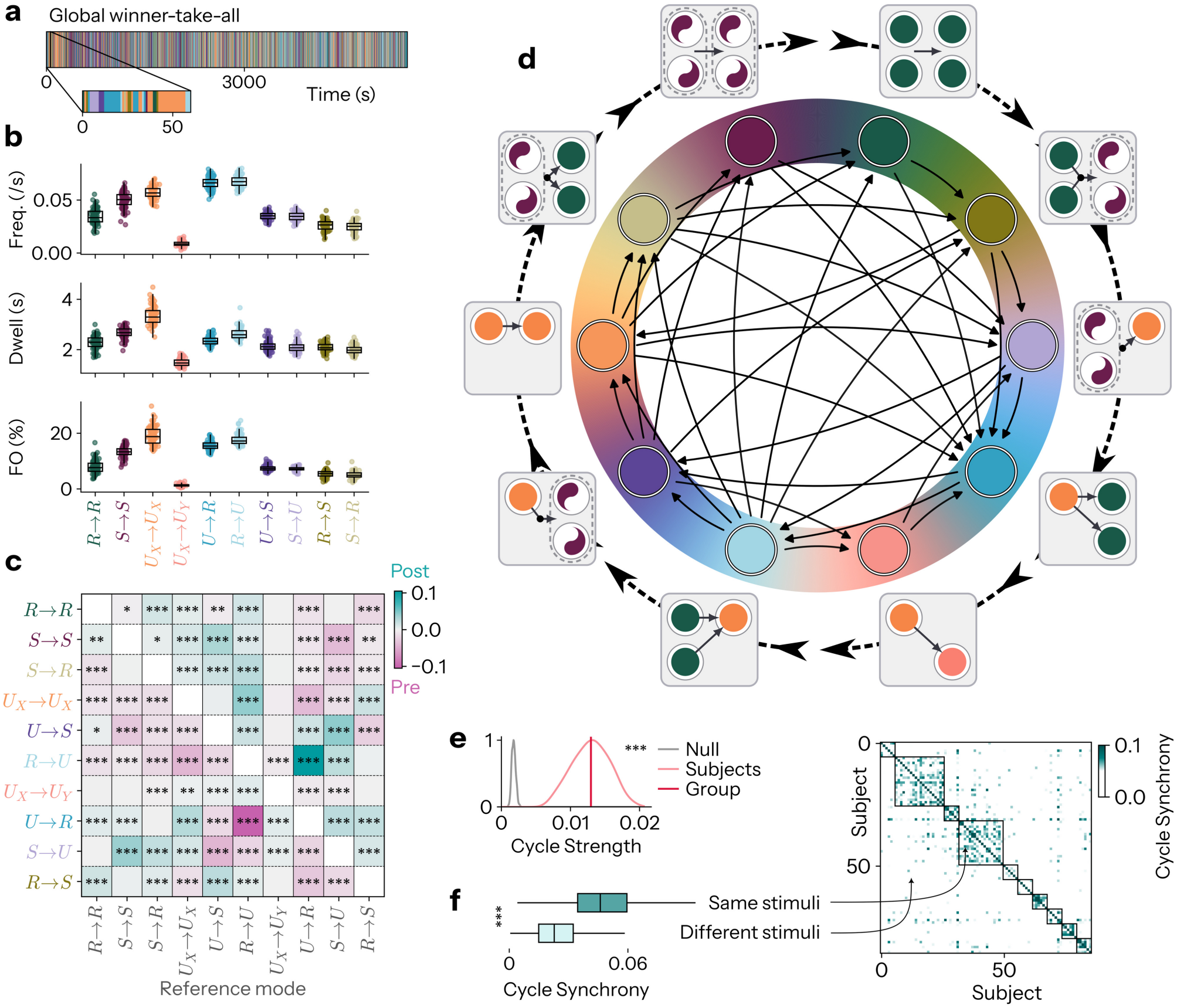
Global information dynamics evolve in a structured cycle synchronised by sensory stimuli. **a** Global winner-take-all time course of information dynamics for an example participant (`sub-1`). Each timepoint is assigned to the mode with the highest mean across all interactions, forming a discrete series. **b** Temporal properties of global information dynamics. First row: the frequency of occurrence or appearance rate of consecutive sequences of each mode. Second row: the mean duration or dwell time of sequences for each mode. Third row: the fractional occupancy (FO) of each mode. **c** Group-level fractional occupancy asymmetry matrix, showing the propensity of each mode (y-axis) to occur in the lead-up to (pink), versus following (teal), each reference mode (x-axis). For each reference mode, the interval between its successive visits is split in half, and the FO of every other mode in the second half is subtracted from its FO in the first half, such that the matrix vanishes if transitions are memoryless and obey detailed balance. Both axes are ordered by the cycle in **d**. Asterisks indicate significant asymmetries in the occurrence of modes before versus after each reference mode (one-sample *t*-test against zero across participants, FDR-corrected Benjamini-Hochberg procedure). **d** Cyclical evolution structure of global information dynamics. Nodes are the ten modes of information dynamics, positioned on the circle so as to maximise the net rotational flow of the FO asymmetry in **c**, thereby recovering the ordering that best expresses the temporal sequencing of global information states (see Methods). Arrows are drawn for FO asymmetry edges surviving *p*FDR *<* 10^−4^, and point from the mode that leads to the mode that lags, such that the net circulation runs clockwise. **e** Cycle strength *S*, the net rotational momentum of the FO asymmetry matrix around the ordering in **d**, at group- and subject-level, against a null (grey) of 1000 permutations that independently relabel the states within each participant and re-optimise the ordering. **f** Cycle synchrony between pairs of participants, quantified as the time-averaged phase-locking value (PLV) between their cycle-phase time series. Pairs are split by whether the two participants watched the same film (dark) or different films (light). Right shows the full matrix of PLV values ordered by film, with black squares delineating the ten films. (*,**,*** indicates p<0.05,0.01,0.001 FDR-corrected Benjamini-Hochberg procedure).

That the brain exhibits different temporal properties for these states raises the question of whether it wanders between them at random, or whether their succession obeys a grammar. We hypothesised that global information dynamics would occur in a cyclical fashion, with certain states predictably preceding or succeeding others. To test this, we used the recently developed temporal interval network density analysis (TINDA)^45^ (see Methods, Supplementary Fig. S11). First, we calculated the asymmetry of fractional occupancies for states over the first and second half of the interstate interval of each state. Put simply, we measure the likelihood that a mode of information dynamics tends to be occupied before versus after another specific state, forming a matrix of transitional asymmetries. Through this, we find that over 80% of the possible edges are significantly asymmetric (74/90 *p*_FDR_ *<* 0.05), suggesting the existence of a conserved “forward” trajectory over time across subjects.

Treating the asymmetry matrix as a weighted directed graph, and positioning modes on a unit circle so as to maximise the net rotational flow between them (see Methods), exposed a single dominant cyclical itinerary (Fig. 5d): *S* → *R* ≺ *S* → *S* ≺ *R* → *R* ≺ *R* → *S* ≺ *S* → *U* ≺ *U* → *R* ≺ *U*_*X*_ → *U*_*Y*_ ≺ *R* → *U* ≺ *U* → *S* ≺ *U*_*X*_ → *U*_*X*_ , and back again. Consistent with the idea that the brain switches between periods of integration and segregation, this itinerary cleaves into two contiguous arcs: one traversing precisely those four modes defined over shared representations alone (*S* → *R, S* → *S, R* → *R, R* → *S*), the other precisely those six that involve differentiated, regionunique information (*S* → *U* , *U* → *R, U*_*X*_ → *U*_*Y*_ , *R* → *U* , *U* → *S, U*_*X*_ → *U*_*X*_). This cycle reveals a repetitive global computational scheme in the naturalistic brain: synergistic information gets dissolved into unique information (*S* → *U*), information is then copied (*U* → *R*) and migrates (*U*_*X*_ → *U*_*Y*_); redundant information is then erased (*R* → *U*), only to be unified back into synergy (*U* → *S*) while unique information persists (*U*_*X*_ → *U*_*X*_). Synergy is then decrypted into redundancy (*S* → *R*), both collective formats are stored (*S* → *S, R* → *R*), and redundancy is finally re-encrypted into synergy (*R* → *S*), whereupon the cycle repeats.

This cycle was slow, strong, ubiquitous, and its structure indifferent to the different films watched by participants. Participants completed a full revolution of the cycle roughly every 13s (12.6 ± 1.1 s; 0.080 ± 0.007 Hz; Supplementary Fig. S12a). Its strength (*S* = 0.013) far exceeded a null distribution in which states were relabelled independently within each participant and the ordering re-optimised (0.0018 ± 0.0003; 1000 permutations, *p*_perm_ *<* 10^−3^, at the permutation floor) (Fig. 5e). It was also present in every single individual: all 86 participants had a positive cycle strength on the group ordering (0.0130 ± 0.0026), with even the weakest (*S* = 0.0076) exceeding the strongest null permutation (*S* = 0.0028). Consistent with this, asymmetry matrices were highly similar between participants (mean pairwise *r* = 0.71 ± 0.09, positive for all 3655 pairs), and each mode held a reliable position in the cycle across individuals (Rayleigh test of phase concentration: all ten modes *p <* 10^−12^, mean *R* = 0.85; Supplementary Fig. S12b). Nor was the cycle a signature of the stimulus itself. Taken separately, each film expressed the same cycle, at the same strength (within- film *S* = 0.0112 to 0.0145, each *p <* 10^−3^; between films, *F* (9, 76) = 1.199, *p* = 0.31), and with the same structure: a distance-based PERMANOVA on the participant-by- participant similarity of asymmetry matrices found no effect of film (pseudo-*F* = 1.184, *p*_perm_ = 0.18), and participants watching the same film were no more similar to one another than to participants watching a different one (*r* = 0.717 ± 0.052 vs 0.715 ± 0.052; *T* = 0.840, *p* = 0.40, *d* = 0.046, *BF*_01_ = 5.99; Supplementary Fig. S12c).

These results show the prevalence of the cycle, but what determines where one is at a particular moment? We predicted that stimuli may entrain the cycle such that it synchronises between individuals observing the same stimulus i.e film. Assigning each mode its position on the cycle and mapping each participant’s state sequence into a cycle-phase time series (see Methods), we found that participants watching the same film were markedly more phase-locked to one another than participants watching different films (PLV = 0.047 vs 0.025; *T* = 18.92, *p* = 3.23×10^−32^, *d* = 2.47, *BF*_10_ = 8.96×10^28^; permutation of film labels, *p*_perm_ *<* 10^−4^, at the permutation floor of 10,000 permutations) (Fig. 5f). This phaselocking also exceeded a circular-shift null that preserves each individual’s occupancy and autocorrelation (0.024; *T* = 19.19, *p* = 1.20×10^−32^, *d* = 2.43, *BF*_10_ = 2.37×10^29^; *p*_perm_ *<* 10^−3^, at the permutation floor of 1,000 surrogates), confirming that it reflects genuine stimulus-driven alignment rather than the mere statistics of the state sequences.

Taken together, we establish the cyclical temporal structure of global information dynamics in the naturalistic brain. This cycle is universal, being present in every individual and identical across films, yet it is not autonomous: environmental context dictates when each individual enters each stage of the cycle.

### Cognitive profile of global information dynamics

Finally, having established that films synchronise the cycle between viewers, we asked what it is about external stimuli that the global mode of information dynamics may be responsive to. To do so, we comprehensively annotated all ten films, describing every TR by what was said, what was shown, and what could be heard, and fusing the three streams into a single description of that moment (see Methods, QC in Fig. S13).

We first scored every TR against a battery of 24 lexical norms spanning sensorimotor strength, concreteness, socialness, affect, motivation and word frequency, and asked whether the moments occupied by each mode departed from a participant-wise circular-shift null (see Methods). Reliable structure was widespread: 62 of the 240 modeby-norm cells survived FDR correction (26%), and every one of the ten modes of information dynamics carried at least one significant norm (Fig. 6a). However, effects were uniformly small (largest 0.046 SD, median 0.023 SD). Strikingly, these profiles respected the two arcs of the cycle. Every one of the four modes defined over shared representations (*R* → *R, R* → *S, S* → *R, S* → *S*) accompanied concrete, visible, and bodily specified content during quiet, low-arousal moments. The copy (*U* → *R*) and erasure (*R* → *U*) of information showed the inverse, occupying abstract, aroused, auditory and motivationally loaded moments. The interconvert synergy and unique information (*S* → *U* , *U* → *S*), on the other hand, were geared towards locomotion.

**FIG. 6.**
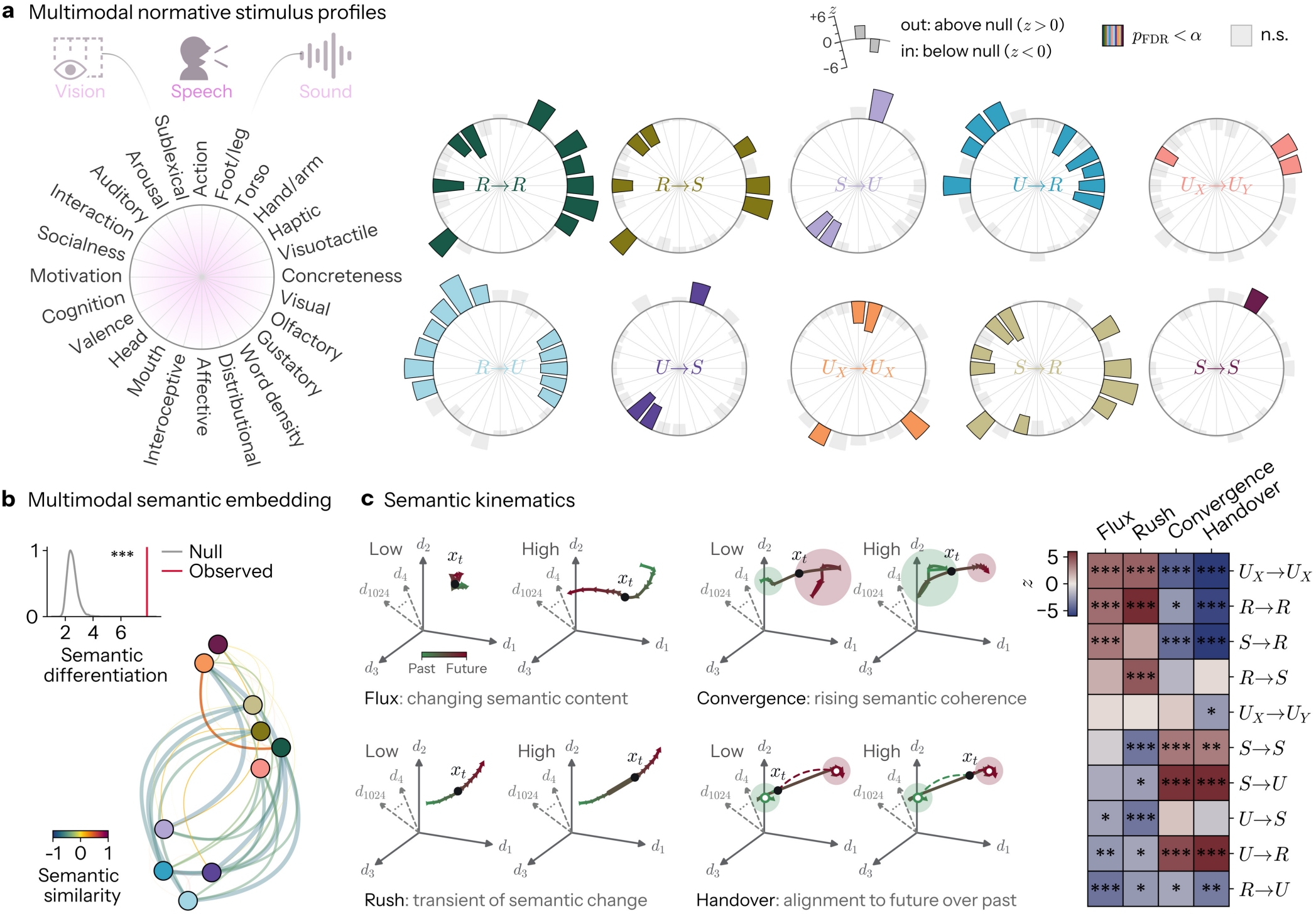
Global information dynamics are associated with distinct psycholinguistic features and semantic kinematics. **a** Normative profiles of each mode of information dynamics. Every TR of all ten films was described by a multimodal annotation of its speech, vision and sound, and scored against 24 lexical norms (see Methods). Each dial shows, for one mode, how far that mode’s mean score on every norm departs from a participant-wise circular-shift null, in units of the null standard deviation. Bars extend outward where a norm is elevated during that mode and inward where it is depressed. Coloured bars survive statistical testing *p*FDR *<* 0.05; grey bars do not. Spokes are ordered by seriating the norms according to how they covary across TRs. **b** Top: semantic differentiation between modes, the pseudo-*F* of PERMANOVA-style partition of the multimodal semantic embedding by mode identity (red), against 5,000 circular-shift nulls (grey). Bottom: semantic similarity network of the modes. Nodes are the ten modes of information dynamics, edges are the cosine similarity between their film-residualised semantic centroids, with width scaled by magnitude and colour by sign. Edges are faded where the pair does not separate from the null (*p*FDR *≥* 0.05). **c** Semantic kinematics. Left: schematics of the four kinematic axes, each contrasting a low against a high value, with the trajectory of the film through the embedding, coloured from past (green) to future (dark red) and *x*_*t*_ marking the present moment. Flux is the rate at which meaning is changing, Rush the excess of that rate over its own recent neighbourhood, Convergence the degree to which the film is settling into semantic coherence, and Handover the degree to which the present moment is semantically closer to what comes next than to what has come before (see Methods). Right: *z* of each mode on each axis against the circular-shift null, with modes ordered by their loading on flux. (*,**,*** indicates p<0.05,0.01,0.001 FDR-corrected Benjamini-Hochberg procedure).

However, content is likely more nuanced and high- dimensional than these 24 normative categories suggest. We therefore embedded each TR’s multimodal description in a dense 1024-dimensional semantic space (see Methods). Asking again whether modes reliably reflect distinguishable stimulus features, but instantiated here as regions of semantic space, we paired each participant’s mode at every TR with the film’s embedding at that moment, and asked whether the moments occupied by each mode cluster apart from one another. Following the logic of PERMANOVA, we partitioned the total semantic variance into a between-mode component (the spread of the ten mode centroids about the grand mean) and a within-mode component (the spread of moments about their own mode’s centroid), summarised as a pseudo-*F* , Indeed, we find the empirical semantic differentiation to exceed all nulls (pseudo-*F* = 7.88, *p*_perm_ *<* 2 × 10^−4^, the permutation floor of 5,000 nulls, Fig. 6b). Though again, the fraction of semantic variance involved was small (*R*^2^ = 1.23 × 10^−4^, 0.012%). This space permits examination of pairwise semantic similarities of global modes of information dynamics (Fig. 6b), of which 30 of the 45 pairs were reliable (*p*_FDR_ *<* 0.05) after FDR correction. The six most semantically remote pairs opposed precisely the two groupings that the normative profiles had already exposed, setting *U*_*X*_ → *U*_*X*_ , *R* → *R* and *S* → *R* against *S* → *U* , *U* → *R, U* → *S* and *R* → *U* (cosine similarity −0.79 to −0.63).

Not only is movie content high-dimensional, but it is deliberately dynamically embedded in the viewer’s cognitive processing window, with curated short-horizon semantic trajectories dictating local, moment-to-moment emotional and thematic turbulence, tension, surprise, and local narrative engagement. Put succinctly, film semantics have their own kinematics. To formalise different types of trajectory through semantic space, we conceive of four constructs, each operationalised with a family of measures: Semantic *Flux, Rush, Convergence*, and *Handover* (see Methods; Fig. 6c). Flux is the rate at which meaning is changing, Rush the excess of that rate over its own recent neighbourhood, Convergence the degree to which the film is settling into semantic coherence, and Handover the degree to which the present moment is semantically closer to what comes next than to what has come before. All four varied reliably across modes (all *p*_perm_ *<* 2.00 × 10^−4^, at the permutation floor of 5,000 nulls), with handover the most strongly differentiated (*SS* = 243.0, against 144.2, 138.0 and 95.2 for convergence, flux, and rush, respectively). Of the 40 mode-by-axis cells, 29 survived *p*_FDR_ *<* 0.05, spanning all ten modes of information dynamics, again with small effects (largest 0.037 SD, median 0.016 SD).

At one pole, *U* → *R, S* → *U* and *S* → *S* held moments whose meaning already anticipated what was coming rather remaining consistent with what had been (*z* = 10.78, 8.70 and 3.33), and at which the film was settling into a more stable semantic position (*z* = 5.00, 5.83 and 3.54 on convergence; all *p*_FDR_ ≤ 0.002). At the other, *R* → *R, S* → *R* and *U*_*X*_ → *U*_*X*_ sat at moments still tethered to what had passed (*z* = −5.54, −6.25 and −6.07), while which meaning was moving quickly (*z* = 3.88, 3.56 and 3.94 in flux; all *p*_FDR_ ≤ 0.002), and, for *R* → *R* and *U*_*X*_ → *U*_*X*_ , accelerating as it went (*z* = 6.30 and *z* = 4.26 on rush).

We confirm that these axes track their intended constructs via three checks (see Methods). Agglomerative clustering of the 45 measures recovered the four assigned families almost exactly (44 of 45 measures; Supplementary Fig. S14); the four axes were close to independent (largest absolute correlation *r* = 0.398; two correlation-matrix eigenvalues above one); and every measure, tested on its own against the leave-one-out composite of its family, reproduced the profile of its axis across modes (mean *r* = +0.71 for flux, +0.82 for convergence, +0.98 for handover and +0.85 for rush; Supplementary Fig. S15).

Taken together, we provide evidence that global information states do not just appear to be merely entrained by a stimulus, but are significantly associated with interpretable features of it: to how abstract and arousing its content is, and to whether the meaning of the moment is settling into what follows or still trailing what came before. These relationships are steady but small: in other words, information dynamics are clearly contingent upon the stimulus, but far from wholly explained by it.

### Replication

To test the robustness of our findings, we repeated every analysis end to end in an independent cohort that we recently acquired at 3T (NNDb-3T+, *n* = 33, all watching *Back to the Future*; see Methods), which differs from the original in field strength, repetition time, spatial resolution, preprocessing, cohort size and design (Extended Data Figs. 1–6; Supplementary Note 2). The information architecture transfers with high fidelity, including the regional and network topography of every mode, the redundancy-to-synergy gradient, individual fingerprints, molecular and cognitive matching, the dynamics of the global information states, their semantic differentiation and their association with semantic kinematics, and, most strikingly, their cyclical traversal, which recurs in every one of the 33 participants. A few results do not carry over: the direct versus indirect anatomy preference of half the modes (*U*_*X*_ → *U*_*Y*_ , *U* → *R, R* → *U* , *R* → *S* and *S* → *R*), two participation maps (*U* → *R* and *R* → *U*), the ranking of modes by reliability and identifiability, and the normative stimulus profiles of the five modes not involving redundancy; computational diversity replicates once both cohorts’ atoms are placed on a common scale via a spatially uninformative correction. Full details of these results are reported in Supplementary Note 2, with further demonstration of the replication in Supplementary Figs. S16–S25.

## III. DISCUSSION

Our findings provide a unified portrait of naturalistic human brain function. By systematically charting the topography, network topology, idiosyncrasy, dynamics, and cognitive relation of the fundamental schemes by which information is stored, transferred, and transformed, we expose the replicable and multidimensional organisation of large-scale neural computation under ecological conditions.

A dominant view in systems neuroscience holds that, in contrast to the sparse code of single neurons, population codes are dense and degenerate, an impression induced by the strong correlation structures that emerge at coarser recording apertures^46–50^. On this account, large-scale brain function is engineered toward resilience and robustness, qualities it purchases by boosting the redundancy of information^51^. Our results temper this picture: among the modes that store information in the same format across time, redundancy is the least abundant vehicle of all, trailing slightly below synergy and drastically below unique information. Thus, information may be more susceptible to the loss of individual regions than models born out of pairwise correlation-based statistics would predict. Along similar lines, a recurring debate in neuroscience concerns whether patches of cortex can be considered functionally specialised^52–55^, with arguments ranging from strict modularity, in which distinct computations are dedicated to distinct areas^56^, to equipotentialism, in which everywhere can do everything in uniform computational capacity^57^. Progress in quantifying the functional repertoire of individual regions has been made by surveying the domain-generality of their response profiles or connectivity across tasks designed to elicit different forms of information processing^58–60^. Here, however, we developed an alternative approach to this problem by introducing a measure of “computational diversity”, capturing how multifaceted the disjoint modes of information dynamics sustained by an interaction are: in other words, how wide their spectrum of computation is in a single context. Our results revealed how every area showed appreciable computational diversity, but this diversity varied systematically across the brain, establishing thalamic, medial, and limbic regions as the most computationally general, and primary sensory regions the most computationally specialised. So, as just detailed, while it is true that the brain is prosperous in persistently regionally unique information, it is far from Balkanised^61^. Moreover, where the brain may lie on the continuum from generality to specificity is contingent upon which brain area is being discussed in the first place.

Macroscale cortical organisation is routinely viewed through the lens of a single principal axis, running from “sensorimotor” to “association” regions^62,63^, and at rest the relative balance of redundancy and synergy follows this trend fairly faithfully^16,17^. By contrast, under naturalistic conditions, the brain’s redundancy-to-synergy gradient spans a novel orthogonal limbic-to-extrastriate axis. Within this gradient, we saw how relative synergy dominance steeply progressed within the dorsal and lateral visual system as one ascended from early retinotopic fields to motion-selective cortex, while the ventral stream at matched remove showed no such ascent, and the frontal eye fields, at the dorsal stream’s terminus, reverted to redundancy dominance outright. Under naturalistic conditions, synergy therefore accrues not with distance from the senses, but with the demand to bind a dynamic compositional scene from parts that no region holds alone; it recedes exactly where that stream ceases to elaborate the scene and begins to select within it, converging upon a single saccadic command. These findings emphasise the distinction between two senses of “association” that macroscale accounts of cortex often employ interchangeably: association as a position in a cortical hierarchy, remote from sensory input, and association as an operation, the conjoining of signals into a representation that none of its parts carries.

Paralimbic and limbic cortices, often left by the wayside in contemporary functional models^62,64,65^, occupy the upper echelons of structural models by virtue of their laminar consistency and numerous bidirectional projections with granular cortex, positioning them as both source and sink of sensory signals^3,66,67^. Recent work has argued for their reinstatement, casting them as the coordinators of interoceptive and exteroceptive inference, through a capacity to move representations between compressed and decompressed formats^68^. Our findings lend this proposal an information-theoretic substrate. It is precisely these regions that host the greatest interchange between synergistic and redundant codes, as well as the greatest migration of unique information. Thus, paralimbic cortices, as well as being at the start of the redundancy-to-synergy gradient, are cardinal engines of moving and transcoding information dynamically between the distributed and harmonised format of synergy and the accessible but excessive format of redundancy.

How the fixed scaffold of white matter gives rise to diverse spatiotemporal patterns of brain activity is an enduring problem in neuroscience^37,69–72^. We add another layer to this question by inferring the latent suite of computations instantiated by this activity. We showed how such information dynamics were partitioned across spatial scales: local shared information storage, intermediate or extremal information modification, and long-range differentiation of informational niches. Given this strong spatial stratification, we found that the communication protocol on the connectome with the most predictive power, for every mode of information dynamics, was navigation, the only model of the four to use physical position rather than topology alone. Thus, geometry is clearly a key and potentially unifying ingredient in explaining the connectomic control of computation. Additionally, we found that broadcasting reliably explained patterns of information dynamics better than strength sequence-preserving nulls, consistent with recent work showing how broadcasting aligns with the network structure of game-theoretic influence^73^. In sum, our results espouse a view where the connectome’s role is not simply “connecting” regions, but to spread signals and multiplex distinct computational regimes across distinct spatial arrangements.

If the brain’s computations are this multifold, what makes an individual’s brain function recognisably theirs? We found a clean dissociation in the means of brain identifiability across modes of information dynamics. Modes of information storage were reproducible within a person, yet similar between people; modes of information transfer and transformation were the least self-consistent, yet the most idiosyncratic. Every mode thereby allowed significant identifiability over surrogate networks, but through opposing means: storage individuates through stability atop a largely shared architecture, transformation through the sheer dispersion of signatures that no two people, and no one person twice, quite repeat. Thus what individuates a brain need not be what endures in it.

We showed how the brain’s information architecture is a plastic functional form that repurposes itself second- to-second in the service of cognition. Our results consequently espouse a view of the brain whereby, at the resolution of a single timepoint, a lone dyadic engagement between two regions can simultaneously instantiate several qualitatively distinct computations. Tracking these interactions, we observed that brain regions display a marked lack of computational use-selectivity, whereby an arbitrary region’s behaviour at one timepoint that strongly takes the form of copying information can take the form of encrypting it. We therefore find a strong computationally primitive form of neural “reuse” ^74^, or “recycling”^75^, that operates on the order of seconds rather than on phylogenetic or ontogenetic timescales.

Seminal formulations in neuroscience, such as Hebb’s assembly phase sequences^76^, and consequently Buzsáki’s neural syntax^77^, posit that cortical processing obeys a grammar, in which basic motifs are chained together in specific temporal orderings with compositional rules^78–81^.

We showed how global information dynamics possess precisely this kind of iterative dynamical structure, instantiated as a directed cycle^45^. However, we observe no evidence that the ordering of this cycle is remixed and repurposed according to differing structures of sensory stimuli. It was universal across films, indicating a canonical ordering to computation. Instead, sensory stimuli appeared to entrain where an individual resided within the universal cycle at each moment. The naturalistic brain thus carries an endogenous “computational clock” whose intrinsic sequence the world cannot rewrite but whose momentary position the world can set, a potential reconciliation of intrinsic dynamics with stimulus-driven processing.

Naturalistic work often reads stimulus-brain relations in terms of what is seen or heard at a specific moment^82^, yet a film is authored as a trajectory, and the meaning of a moment carries a direction as well as a subject. What most sharply separated global modes of information dynamics was accordingly not the semantic position of a moment, nor the rate at which meaning moved through it, but its handover: whether the moment already belonged to what was coming or was still tethered to what had been. We found that unpacking a combined representation into parts that single regions can carry, or duplicating a region’s own representation into a shared one, arrived as the film settled onto its next meaning. The conservation of a representation in the format it already occupies, or decryption of a jointly expressed format to a doubly expressed one, held moments still trailing what had passed, through which meaning was moving quickly and accelerating as it went, where nothing has yet settled that is worth committing and the remaining task is to protect what one has.

Our conclusions are bounded in several ways. The decomposition is bivariate and evaluated at a single lag, so the computations we report leave information dynamics distributed over three or more regions and unfolding over longer horizons beyond its reach. Our time-resolved atoms are likewise pointwise quantities evaluated against a covariance estimated over the whole scan, so like other time-resolved approaches, what fluctuates is a region pair’s momentary position within a stationary model^44,83–85^. Because the atoms are computed upon BOLD, regional differences in haemodynamic timing and signal quality can imprint themselves upon topographies we read as computational, though their transfer across field strength, resolution and preprocessing argues against a purely vascular origin. More fundamentally, information-theoretic quantities are observer-relative: they measure the reduction of our own uncertainty under a probability model that we impose upon variables, so the atoms describe the brain but cannot be claimed to be intrinsic to it. The fraction of stimulus variance the modes of information dynamics command, though reliable, is very small, and we cannot say whether this reflects a genuinely loose coupling to content, or the attenuation imposed by our measurements and automated annotation. With no behaviour recorded or experience sampling alongside our brain data, every cognitive reading we offer is anchored on the stimulus side alone, meaning that we do not show how this architecture matters from the perspective of the subject. Finally, our participants lay motionless inside a large magnet in a basement, watching curated stories upon which they could not act, and to which they need not respond^86^. This is naturalistic in the sense that participants were not performing an abstract task on abstract stimuli, or being encouraged to perform no identifiable task at all as in rest, but it is not yet truly naturalistic.

To conclude, in deciphering the spatiotemporal layout of information dynamics, we furnish the first encompassing map of not just where information resides in the human brain, but of what becomes of it. In doing so, we offer an account of the computational principles subserving adaptive cognition in our weird and wild world.

## IV. METHODS

### Participants

The human MRI data in this study is from the “Naturalistic Neuroimaging Database” (NNDb). Full details can be found in our previous publication^25^, but are reiterated here in brief. The database contains fMRI scans from (*N* = 86) participants who watched one of ten featurelength films they had not previously seen (500 Days of Summer, *n* = 20; Citizenfour, *n* = 18; and *n* = 6 each for The Usual Suspects, Pulp Fiction, The Shawshank Redemption, The Prestige, Back to the Future, Split, Little Miss Sunshine, and 12 Years a Slave). All subjects were native English speakers who were right-handed, had no visual or auditory deficits, and no history of psychiatric or neurological illness. The participant pool included 42 females and 44 males, with ages spanning from 18 to 58 years. Each participant gave written consent for their anonymised data to be used and shared.

### MRI acquisition

Functional and anatomical images were acquired using a 1.5T Siemens MAGNETOM Avanto scanner equipped with a 32-channel head coil. The scanning protocol utilised a multiband EPI sequence^87^ (TR = 1 s, TE = 54.8 ms, 75^°^ flip angle, 40 interleaved slices, and 3.2 mm isotropic resolution). The sequence employed a 4x multiband factor, without in-plane acceleration and with “leak block”, to minimise cross-slice aliasing^88,89^. Following the functional scans, anatomical images were obtained using a 10-minute high-resolution T1-weighted MPRAGE sequence (TR = 2.73 s, TE = 3.57 ms, 176 sagittal slices, 1.0 mm resolution). During scanning, movies were displayed on a 22.5 x 42cm screen that was back-projected and mirrored, positioned behind the head coil. Participants viewed the screen through a mirror mounted above their eyes at a 19.0^°^ viewing angle. Sound was delivered through MRI-compatible scanner noise-attenuating headphones^25^.

### Preprocessing

Anatomical images were corrected for image intensity non-uniformity, deskulled, and non-linearly aligned to the MNI-N27 template^25^. This was followed by inflation, registration and segmentation using FreeSurfer’s recon-all with standard settings^90^. White-matter and ventricle regions of interest (ROIs) were extracted for use as noise regressors. Functional images were first corrected for slice-time differences and de-spiked. EPI volumes were aligned to the volume with minimal motion and then co-registered to their respective anatomical scans. These aligned time series were then warped to the MNI template space. Time series were then spatially smoothed with 6mm full-width half-maximum^91^, normalised to have a sum of squares of one, and detrended with six motion regressors, along with the previously defined white matter and cerebrospinal fluid regressors^92^. The detrended time series were subsequently concatenated and analysed using spatial independent components analysis across 250 dimensions^93^. Data was also band-passed between 0.008– 0.09 Hz to remove high-frequency noise and low-frequency drifts. To ensure the computational feasibility of our analysis, we parcellated the functional data by averaging all the voxels in an area prescribed by combining the 1000 cortical ROIs from the Schaefer atlas^94^ and the 54 subcortical ROIs from the Tian atlas^95^.

### Structural connectome

Owing to the lack of diffusion MRI (dMRI) data for the participants from the NNDb, we used dMRI data from the 1,065 participants of the Human Connectome Project (HCP)^96^. The dMRI acquisition for the source data is described in detail elsewhere in consortium papers^96–98^, but in brief, scans were acquired with a Siemens 3T Skyra using a 2D EPI sequence featuring spin-echo contrast and single-shot acquisition, combined with triple-band slice excitation and monopolar diffusion gradient pulse. The acquisition parameters were: 1.25 mm isotropic resolution, TR = 5500 ms, TE = 89.5 ms, with *b*-values of 1000, 2000, and 3000 s/mm^2^. The total number of diffusion sampling directions was 90 for each shell, in addition to 6 *b*_0_ images^96–98^.

We adopted previously reported procedures to accurately reconstruct the structural connectome^99,100^. dMRI data were first corrected for eddy currents and susceptibility artefacts, and model-free q-space diffeomorphic reconstruction (QSDR) was employed to perform a diffusiblespin-conserving transformation and non-linear warping to MNI space^101^. Per-subject spin distribution functions were aggregated across the 1,065 participants to yield the population-averaged template. Whole-brain deterministic tractography was performed using a modified fibre assignment by continuous tracking (FACT) algorithm, with an angular cut-off of 55^°^, a step size of 1.0 mm, a minimum streamline length of 10 mm, a maximum length of 400 mm, and no spin-distribution-function smoothing, until 10,000,000 streamlines were reconstructed^102^. Streamlines were rejected if they exhibited improper termination locations, based on a white-matter mask created by applying an anisotropy threshold of 0.6 times Otsu’s threshold to the spin distribution functions. The region-to-region structural connectome was then extracted by projecting the tractogram onto the same volumetric parcellation used for the functional data, under an endpoint-based assignment rule whereby a streamline contributes to an edge only if both of its terminations fall within the two corresponding regions. To account for the well-known dependence of streamline counts on region size and tract length, edge weights were normalised so that the weight between regions *i* and *j* was given by the streamline count divided by the product of the mean of the two regional volumes, (*V*_*i*_ + *V*_*j*_)*/*2, and the mean streamline length connecting the two regions.

### Integrated information decomposition via information geometry

Consider a bivariate stochastic process **Z** = (*Z*^1^, *Z*^2^) that evolves from a past state 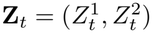 to a future state 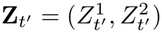. In the applications considered here, *Z*^1^ and *Z*^2^ represent the fluctuating amplitude of the BOLD response in two brain regions, and *t* and *t*^′^ *> t* two consecutive timesteps. The total amount of information shared between the past and future is quantified by the time-delayed mutual information (TDMI):

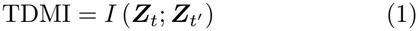

The TDMI is a gross statistic, as it combines qualitatively distinct forms of information into a single scalar quantity. Our aim is therefore to decompose the TDMI into its constituent and most fundamental dynamical information components.

To achieve this, we use the recently introduced Integrated Information Decomposition (ΦID)^4^. Unlike previous information decomposition approaches, ΦID enables the simultaneous differentiation of both parts and wholes with respect to the past and future. The TDMI can consequently be expressed as a sum of non-overlapping dynamical information “atoms”:

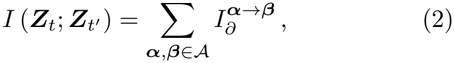

where *A* denotes the double-redundancy antichain lattice introduced in Ref.^4^. Here, ***α*** → ***β*** specifies the dependency mode of an information component in the past and future, respectively, with ***α, β*** ∈ {*R, U*_1_, *U*_2_, *S*} denoting redundancy, the two unique components, and synergy. For example, *S* → *R* represents the information that is synergistic in the past and gets decrypted or multilocalised, becoming redundant in the future of the two variables. Hence, pairing each of the four past dependency modes with each of the four future modes gives a total of sixteen distinct information atoms.

Each mutual information between a subset of the past variables and a subset of the future variables can itself be expressed as a sum of the corresponding ΦID atoms^4,5^. However, for a bivariate process, there are sixteen unknown atoms, and only nine such source-target mutual information quantities, meaning the decomposition is alge- braically underdetermined, and cannot be solved without additionally defined constraints.

To solve this, we use the information-geometric decom- position we recently introduced in Ref.^32^. This framework supplies a geometric definition of synergy at multiple orders, establishing a direct correspondence between these quantities and the ΦID atoms. Below, we provide a brief explanation of this approach, referring to the original publication for the full mathematical treatment and geometric interpretation^32^.

In information geometry, probability distributions are represented as points in a statistical manifold, whose coordinates can be chosen to isolate distinct statistical interactions among the variables. For the present bivariate system **Z** = (*Z*^1^, *Z*^2^), let 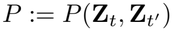 denote the full joint distribution. The corresponding statistical manifold *M* can be written as

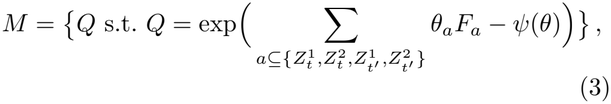

where *F*_*a*_ is the sufficient statistic associated with the subset *a, θ*_*a*_ is its corresponding natural parameter, and *ψ*(*θ*) is the log-partition function ensuring normalisation. In these coordinates, each *θ*_*a*_ encodes a specific statistical interaction. Consequently, setting *θ*_*a*_ = 0 removes that interaction while leaving the remaining interaction coordinates unchanged.

This representation allows higher-order dependencies to be quantified geometrically. Specifically, removing a set of interaction terms defines a submanifold of *M* , and the contribution of the removed interactions can be quantified by the Kullback–Leibler (KL) divergence between the original distribution and its projection onto that submanifold. In particular, higher-order synergy corresponds to information that cannot be accounted for by lower-order source-target dependencies. It can therefore be isolated by constructing distributions in which the relevant pairwise pathways have been removed and measuring the residual distance from the original distribution ^32^.

To make this construction explicit, let’s consider a pair of indices *i, j* ∈ {1, 2} and define the following submanifold of *M* :

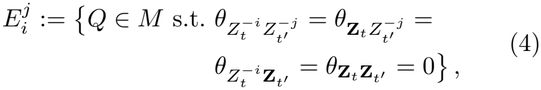

where −*i* denotes the index complementary to *i*. In 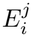, all interactions involving both 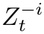 and 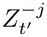 are switched off, leaving only the dependencies mediated by 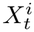 and 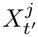 . In other words, this submanifold describes the class of distributions where 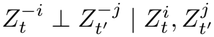. Denoting as 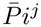 the *m*-projection of *P* onto 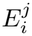, we can then quantify the information removed by this projection by calculating the Kullback-Leibler distance between the original distribution *P* and its projection 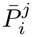 onto the submanifold 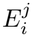, which corresponds to the conditional mutual information

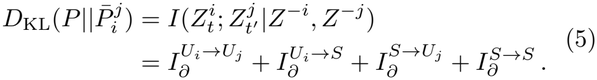

Thus, each projection isolates the subset of ΦID atoms that involve unique or synergistic information on the selected source and target components.

Considering Eq. (5) for all four combinations of *i, j* ∈ 1, 2 reveals that the only atom common to all four conditional mutual information values is 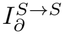. Consequently, the distributions generated by interpolating between the four projected distributions 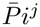 span a family of models that can account for pairwise and three-variable pathway contributions, but not the fully synergistic 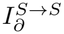 atomic dependency.

Therefore, the highest-order synergy is the residual distance of the true distribution from this set of reducible models, namely the smallest KL divergence between *P* and the interpolated projections:

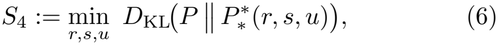

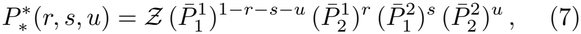

where *Z* is a normalising constant. Following from the construction above, this quantity corresponds exactly to the ΦID atom that is synergistic in both the past and future, i.e. 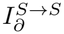.

The same geometrical construction can be applied to the three-variable marginal distributions 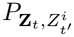 and 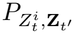 yielding the third-order synergies associated with each source-target combination, each of which is identified as a pair of atoms in the ΦID algebra:

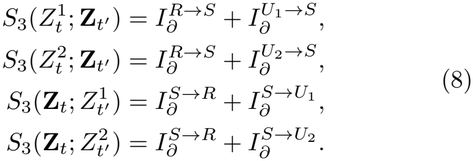

Finally, applying the construction with either the source or target pair treated as a single bivariate group gives

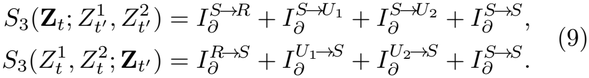

Together with the nine source-target mutual informations, these seven geometric synergy measures provide the seven additional constraints required to close the ΦID system. Hence, the resulting sixteen independent equations uniquely determine all sixteen information atoms.

In practice, we applied this estimator independently to every pair of brain regions, treating their standardised BOLD time series as the two components of the bivariate process **Z**. The past and future states were defined as the BOLD activity at consecutive time points, such that the decomposition characterises the information shared between the activity of the two regions at one time point and their activity at the next timestep.

To make modes more directly comparable, ensuring symmetry and invariance under relabelling, we combine unique contributions that differ in identity. For ease of notation and additional clarity, throughout this manuscript we used the following shorthands to express the ΦID atoms:

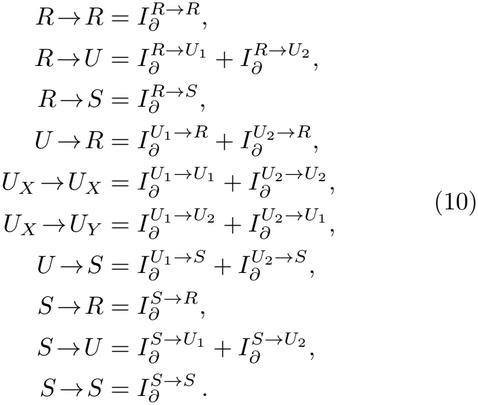

Here, the symbols *X* and *Y* are employed to distinguish whether unique information is transferred to the same or the other brain area, respectively.

### Time-resolved information dynamics

The information-theoretic framework described above characterises temporal dependencies by quantifying the statistical interdependence between past and future states. However, these quantities are inherently time-averaged: they are defined as expectations over a joint probability distribution. To obtain a temporally-resolved characterisation, we consider the pointwise (or local) versions of the information-theoretic quantities, which assign an information value to each individual observation of the brain.

More generally, a pointwise information quantity is obtained by removing the expectation from the corresponding information-theoretic expression, and evaluating the resulting information content for a particular realisation 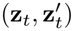. In such a case, the pointwise time-delayed mutual information is given by

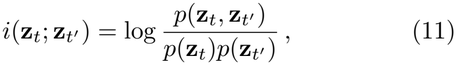

where **z**_*t*_ and 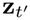denote particular observed states. The usual mutual information is recovered by averaging this quantity over the joint distribution:

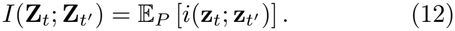

The same construction applies to the Kullback–Leibler divergence. For two probability distributions *P* and *Q* defined over the same space *Z*, the pointwise KL divergence is

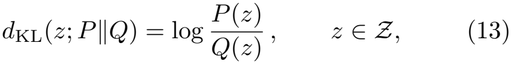

and the corresponding KL divergence is recovered by averaging over *P* :

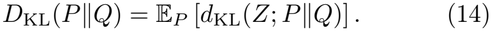

In particular, for the Gaussian distributions considered here, the pointwise quantities are evaluated directly as log-density ratios at each observed state.

In this work, we extended the ΦID^IG^ framework to this local setting, enabling the decomposition of information dynamics at the level of individual observations. Once the relevant probability distributions have been estimated and the ΦID^IG^ construction performed, all quantities entering the resulting system of equations can be evaluated instead using the pointwise mutual information and KL divergences defined above, yielding solutions to each ΦID atom at every time point. Averaging these local atoms over time recovers the corresponding global ΦID^IG^ decomposition.

### Computational diversity

To assess the extent to which an interaction hosts a diversity of modes of information dynamics, we introduce a novel quantity, computational diversity (*D*).

Collecting the *n* = 16 atoms of a region pair into a vector **a** = (*a*_1_, … , *a*_*n*_), with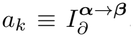, we form the normalised squared weights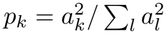, a probability distribution describing each atom’s fractional contribution to the profile, and take its participation ratio , normalised by the number of atoms,

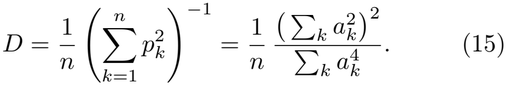

It ranges from 1*/n*, when a single atom carries all the information (a maximally specialised computational profile), to 1, when all *n* atoms contribute equally (a maximally diverse one). Equivalently, *D* is the Rényi- 2 (inverse-Simpson) effective number of atoms divided by *n*. We computed *D* for every region pair, yielding a symmetric 1054 × 1054 diversity matrix per participant.

### Redundancy-to-synergy rank gradient

Following prior work^16^, after obtaining whole-brain networks for each information atom, we determine the position of each ROI in a gradient of relative engagement in *S* → *S* or *R* → *R* representation. We first calculate the mean of each brain region’s global interactions for *S* → *S* and *R* → *R*, and rank them in descending order, respectively. Then we subtract the *R* → *R* rank from the *S* → *S* rank, yielding a gradient from negative (relative *R* → *R* >*S* → *S*) to positive (relative *R* → *R* <*S* → *S*), that represents ROI-wise bias relative to the rest of the brain toward a particular mode of information integration.

### Networks

#### Network normalisation

To avoid issues with negative weights and scaling differences between types of information, we normalise the weights in each network via an empirical probability integral transform, akin to the copula transform in statistics. Concretely, for a network with *m* weights {*w*_*ij*_}, every weight is replaced by the empirical cumulative distribution function evaluated at it,

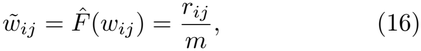

where *r*_*ij*_ is the rank of *w*_*ij*_ among all *m* weights, with ties assigned their average rank. In other words, the smallest weight is mapped to 1*/m*, the largest to 1, and all others are linearly interpolated in between. This is a non-parametric, rank-based procedure that maps the weights of each network onto the uniform ordinal distribution on the unit interval, [0, 1].

#### Global efficiency

We calculate the capacity for topological integration of each network via the average of the inverse distances between nodes, its global efficiency:

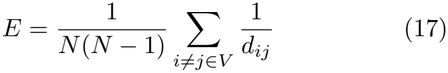

where *d*_*ij*_ is the distance between the i-th and j-th nodes.

#### Complexity

We quantify the structural complexity of each network through its graph von Neumann (entanglement) entropy^103,104^. The combinatorial graph Laplacian *L* = *D* − *W* , where *W* is the weight matrix and *D* the diagonal matrix of node strengths, is treated as an unnormalised density operator; rescaling by its trace yields a density matrix *ρ* = *L/*tr(*L*) with unit trace and non-negative eigenvalues *λ*_*i*_ = *µ*_*i*_*/* ∑_*k*_ *µ*_*k*_, where *µ*_*i*_ are the eigenvalues of *L*. The von Neumann entropy of this operator is then simply:

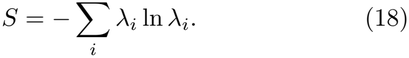

Akin to the Shannon entropy. This entropy is thus interpreted as the level of entanglement of the statistical ensemble of edges. A high entropy reflects a complex, heterogeneous architecture whose connectivity is distributed across many spectral modes, whereas a low entropy indicates a more regular and mirrored organisation dominated by a few modes.

#### Hubness

To understand structural inhomogeneity in the network, we characterise the extent to which it exhibits a hierarchical heavy-tailed organisation by computing the s-metr ^105^:

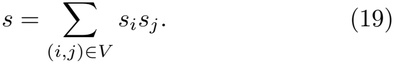

Where *s*_*i*_ is the weighted degree (strength) of node *i*. A high *s* value indicates that connectivity is concentrated, with high-degree nodes predominantly connecting to other high-degree nodes, and low-degree nodes connecting with other low-degree nodes. In contrast, a low *s* value indicates that connectivity is uniform across the network.

#### Modularity

We calculate the modularity of each network via the classic quantity *Q*:

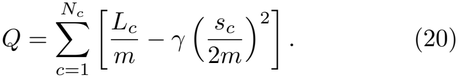

Where the sum iterates over the *N*_*c*_ communities, *L*_*c*_ is the total intra-community weight, *m* is the total edge weight, *s*_*c*_ is the total strength of community *c*, and *γ* is the resolution parameter which we set to 1^106^.

#### Participation

To capture the extent to which a region’s connections are distributed across communities rather than concentrated within its own, we computed the participation coefficient. Given a partition of the network into *N*_*c*_ communities, the participation coefficient of node *i* is

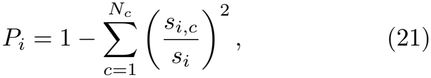

where *s*_*i,c*_ is the summed weight of node *i*’s connections to community *c*, and *s*_*i*_ = ∑_*c*_ *s*_*i,c*_ is its total strength. *P*_*i*_ tends to 0 for a node whose connections lie almost entirely within its own community (a provincial node), and towards 1 for a node whose connections are evenly spread across all communities (a connector node).

#### Community detection

To identify distinct subsystems of each network for the computation of modularity and participation, we performed community detection with multiresolution consensus clustering (MRCC)^107^. In a first pass, the network was partitioned with the Leiden algorithm under a Constant Potts Model (CPM) objective^108^ across 20 linearly spaced values of the resolution parameter *γ*, and the range of *γ* yielding non-trivial partitions (more than one community but fewer than *N*) was retained. A second, denser sweep of 30 resolutions spanning this range was then used to build a consensus co-assignment matrix, whose entries record the frequency with which each pair of regions was placed in the same community. This consensus matrix was itself partitioned with Leiden under a CPM objective (resolution parameter 0.1), and any communities smaller than two nodes were merged into their highest-affinity neighbour to give the final partition.

#### Multiplexing

By framing each network of information dynamics as a layer in a multilayer network^109^, we can characterise how the brain’s connectivity is shared across, or specialised between, these layers. Taking the rank-normalised groupaverage network of each mode of information dynamics as a layer over the same set of regions, every connection (*i, j*) has a profile of its strength across modes, which we summarise by its mean, the consensus *µ*_*ij*_, and its standard deviation, the contrast *σ*_*ij*_. A high consensus marks a connection that is consistently expressed by every mode of information dynamics, whereas a high contrast marks one that is strongly expressed in some modes and absent in others.

#### Mode invariance

At the level of nodes, we instead ask whether the connectivity pattern of each region is preserved across modes. For region *r*, let 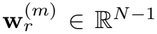 denote its connectivity profile, the vector of its edge weights to all other regions, in mode *m*. We quantify its mode invariance *v*_*r*_ as the mean absolute cosine similarity between these profiles over all pairs of the *A* = 10 modes of information dynamics,

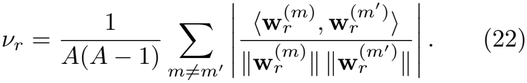

A high *v*_*r*_ indicates that a region connects to the same partners irrespective of the mode, its connectivity being multiplexed across modes, whereas a low value indicates that it reconfigures its connections from one mode to the next.

#### Surrogate networks

To establish whether an observed relationship between two networks exceeded that expected from their basic geometry, we generated randomised surrogates that preserve the network’s size, density, weighted degree (strength) sequence, and weight distribution. Starting from a fully randomised network, following the procedure of Milisav et al^39^, we perform simulated annealing to minimise the energy function 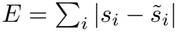, where *s*_*i*_ and 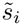 are the empirical and surrogate strengths of node *i*, respectively. At each iteration a pair of weights is swapped, accepted if it lowers the energy and otherwise with the Metropolis probability exp(−Δ*E/T*). Temperature *T* was initialised at 100 and reduced by 25% over each of 20 annealing stages of 5,000 permutations. For each network we generated 1,000 surrogates, yielding a null distribution for permutation *p*-value extraction.

### Communication models

To relate the large-scale information dynamics to different protocols the brain may deploy for polysynaptic signalling, we computed a spectrum of network communication models on the structural connectome^71^. For models that operate on connection costs, the edge weights *w*_*ij*_ were first remapped to lengths through the monotonic transform *l*_*ij*_ = − ln (*w*_*ij*_*/* max(*w*)) , so that stronger connections form shorter, less costly channels; the Euclidean distance between regional centroids was used where a spatial metric was required. We considered four models spanning the routing-to-diffusion spectrum^37^.

#### Routing

Routing protocols assume that signals travel along the single most efficient path between two regions. Writing a path from *i* to *j* as the sequence of nodes it visits, Ω_*ij*_ = {*i, u*_1_, … , *u*_*k*_, *j*}, its cost is the sum of the connection lengths traversed, and the shortest-path length is the minimum of this cost over all paths,

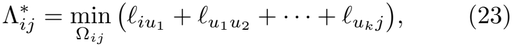

which we obtain with the Floyd–Warshall algorithm. The shortest-path efficiency is its reciprocal,

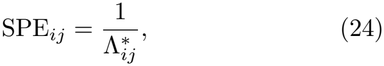

so that regions joined by short, strong paths communicate most efficiently^110^.

#### Navigation

Navigation is a decentralised, geometrically greedy routing strategy that requires only local knowledge of node positions^111,112^. Let *D*_*ij*_ denote the Euclidean distance between regions *i* and *j*. Navigating from a source *i* towards a target *j*, the signal advances at each step from the current node *u* to whichever of its neighbours lies spatially closest to the target,

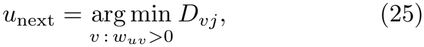

terminating successfully when *j* is reached and failing (length ∞) if a node is revisited. The navigation path length is the total Euclidean distance traversed along the resulting path 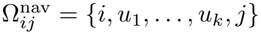,

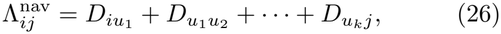

and navigation efficiency is its reciprocal,

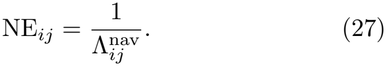

Broadcasting assumes that signals diffuse simultane-ously along all walks of the network at once. Communicability sums the contributions of all walks between two regions, weighting each walk of length *n* by 1*/n*! so that short walks dominate; for the weighted connectome *W* this reduces to the matrix exponential:

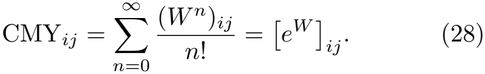

High communicability indicates that two regions are linked by many short, parallel walks^113^.

#### Diffusion

Diffusion models signalling as an unbiased random walk that explores the connectome with no preferred direction and a large propensity for recurrence or retransmission. The walk is governed by the transition matrix Ŵ , obtained by row-normalising the connectome, ŵ_*ij*_ = *w*_*ij*_*/* ∑_*u*_ *w*_*iu*_, with stationary distribution ***ω*** satisfying ***ω***Ŵ = ***ω***. Stacking copies of ***ω*** into the matrix 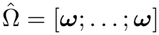 and forming the fundamental matrix 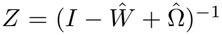, the mean first-passage time, the expected number of steps for a walker departing region *i* to first reach region *j*, is

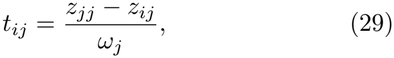

and diffusion efficiency is its reciprocal,

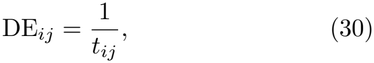

so that regions between which a walker passes quickly are taken to communicate efficiently^114,115^.

### Individual differences

To characterise the within- and between-subject variability of our different modes of information dynamics, we first quantified the reliability of each network edge via the intraclass correlation coefficient ICC(1, 1). Treating the first and second halves of the scan as *k* = 2 repeated measurements of an edge across the *N* subjects,

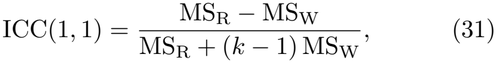

where MS_R_ is the between-subject mean square and MS_W_ the within-subject mean square; with *k* = 2 the denominator reduces to MS_R_ + MS_W_. Writing the two half-scan weights of an edge for subject *n* as 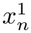 and 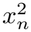 , with subject mean 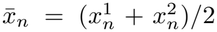, these are 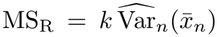, the unbiased varia nce of the subject means, and 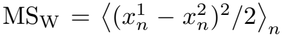, the average within-subject squared difference. ICC(1, 1) approaches one when an edge varies far more between subjects than between the two halves of a given subject, and we summarised reliability at each node as the mean ICC across its edges.

Next, we performed “brain fingerprinting”^116–118^. Constructing a non-symmetric identifiability matrix, containing for each mode of information dynamics, the Pearson correlation between network edges in the first- and second-half of the scan within subjects (diagonal), and the correlation between edges in the first- and second-half of the scan between subjects (off-diagonals). For each mode of information dynamics, and each individual, *Self similarity* (I_self_) is therefore the diagonal element (correlation between 1st and 2nd half of their scan), and *Other similarity* (I_other_) is the average of the off-diagonal elements in its corresponding row and column, (average correlation between 1st half of their scan and 2nd half of everyone else’s scan). I_self_ coarsely represents how stable a particular whole-brain configuration of information is over time, whereas I_other_ represents how indistinguishable it is from other subjects. Identifiability then is taken as I_self_ − I_other_, and quantifies the extent to which a mode of information dynamics is consistent intra-subjectively, but distinctive inter-subjectively.

We assessed the significance of identifiability against a null distribution obtained by reassigning each subject’s self-comparison to a randomly chosen other subject (1000 label derangements) and recomputing I_self_ , I_other_ and their difference for each permutation.

### Entropy rate

To quantify how temporally diverse each region’s engagement in a mode of information dynamics is, we estimate the state-space entropy rate of its global, time-resolved fluctuations *x*_*t*_ in that mode. The entropy rate is the uncertainty that remains in the present sample once its entire past is known,

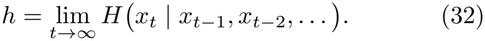

To estimate it, we follow the procedure of Mediano et al.^44^. Treating *x*_*t*_ as a noisy read-out of a hidden state **z**_*t*_ with linear, Gaussian dynamics, written in innovations form as:

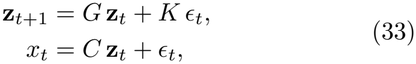

where *G* is the state-transition matrix, *C* the observation matrix, *K* the Kalman gain, and *ϵ*_*t*_ the innovations: the white, zero-mean one-step-ahead prediction errors with variance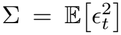. Since the hidden state captures everything the past reveals about the present, the entropy rate reduces to the entropy of these innovations, which for a Gaussian process is simply:

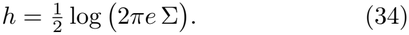

We identified the state-space model by a subspace method, setting the past and future regression horizons to twice the order of an autoregressive model chosen by the Hannan-Quinn criterion, and fixing the dimension of **z**_*t*_ by the singular-value criterion.

Thus, we characterise how much a region explores possible trajectories in a mode of information dynamics, or alternatively, the intrinsic unpredictability or prediction error of its global engagement in that mode^44^.

### AHBA data

Regional microarray expression data were obtained from 6 post-mortem brains (1 female, ages 24.0–57.0, 42.50 ± 13.38) provided by the Allen Human Brain Atlas (AHBA, human.brain-map.org;^119^). Data were processed with the abagen toolbox (version 0.1.3; github. com/rmarkello/abagen)^120^.

First, microarray probes were reannotated using data provided by^120^; probes not matched to a valid Entrez ID were discarded. Next, probes were filtered based on their expression intensity relative to background noise^121^, such that probes with intensity less than the background in ≥50.00% of samples across donors were discarded, yielding 31,569 probes. When multiple probes indexed the expression of the same gene, we selected and used the probe with the most consistent pattern of regional variation across donors (i.e., differential stability;^122^), calculated with:

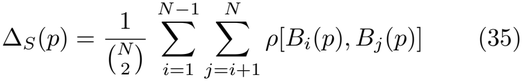

where *ρ* is Spearman’s rank correlation of the expression of a single probe, *p*, across regions in two donors *B*_*i*_ and *B*_*j*_, and *N* is the total number of donors. Here, regions correspond to the structural designations provided in the ontology from the AHBA.

The MNI coordinates of tissue samples were updated to those generated via non-linear registration using the Advanced Normalization Tools (ANTs; github.com/chrisfilo/alleninf). Samples were assigned to brain regions in the extended Schaefer-1054 atlas used herein if their MNI coordinates were within 2 mm of a given parcel. All tissue samples not assigned to a brain region in the provided atlas were discarded.

Inter-subject variation was addressed by normalizing tissue sample expression values across genes using a robust sigmoid function^123^:

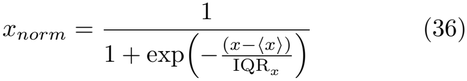

where ⟨*x*⟩ is the median and IQR_*x*_ is the normalized interquartile range of the expression of a single tissue sample across genes. Normalized expression values were then rescaled to the unit interval:

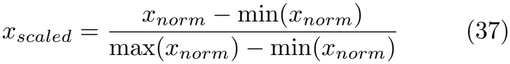

Gene expression values were then normalised across tissue samples using an identical procedure. Samples assigned to the same brain region were averaged separately for each donor and then across donors. To ensure spatially dense expression values, we followed the procedure of Wagstyl et al.^124^, performing nearest-neighbour interpolation of missing parcels, and smoothening with a 20mm full-width at half maximum Gaussian kernel.

We then subsetted the 15,633 genes from the AHBA library to those of neurobiological relevance. Specifically, we followed the categorical gene sets used by Burt et al.^125^:

- *Brain-specific*: 2,412 genes from supplementary dataset 1 of Fagerberg et al.^126^: those whose expression in brain tissue was four times higher than the median across all 27 tissues sampled^127^. 1,905 genes retained.
- *Neuron-specific*: 2,529 genes from supplementary dataset S3b of Cahoy et al.^128^: those whose log-expression in P7n neurons was 0.5 times greater than the median log-expression across the 11 CNS cell types^127^. 2,319 genes were retained.
- *Synaptic*: 1,885 genes aggregated from Synap-tomeDB^129^ encoding proteins of the presynaptic nerve terminal, presynaptic active zone, synaptic vesicles, and postsynaptic density. 1,746 genes retained.

### Neurosynth data

Continuous measures of the association between brain areas and cognitive categories were taken from the whole-brain automated term-based meta-analytic tool Neurosynth^130^, based on >15,000 published functional MRI studies. Specifically, we used the volumetric *P* (function|activation) association test maps for the 123 terms that overlap between Neurosynth and the Cognitive Atlas, a public ontology of cognitive science^131^. These range from umbrella terms (e.g perception or attention), to specific processes (e.g reinforcement learning or mental imagery), behaviours (e.g navigation or sleep), and emotional states (e.g fear or anxiety). These maps therefore measure the prevalence of reporting for a given cognitive domain or behaviour in the functional neuroimaging literature with respect to alterations in a given voxel’s activity.

### Matching

To investigate the extent to which information dynamics reflect the contribution of lower-level biophysical details, or contribute to higher-level functional properties, we performed spatial correlations between the topographies of mode of information dynamics, for each participant at each time point, with each map in the refined the AHBA and Neurosynth datasets, parcellated to the same combined atlas. For each participant, we then take the maximum value for each timepoint, for each mode of information dynamics, as their quality of matching with chemoarchitecture, and cognition, which are respectively averaged across time to yield scalar scores. The former we term its “molecular match”, and the latter following Luppi et al. its “cognitive match” ^132^.

### Global information states

To gain a simple and interpretable global view on the status of representation and computation at each moment, we use a winner-take-all approach. After z-normalising each region’s global interactions for each mode of information dynamics, we average across all regions at each time point and assign each timepoint the categorical label (state) of the mode of information dynamics that is strongest.

To characterise the temporal dynamics of each state, we take the subject-specific state sequences and compute (1) the fractional occupancy, the proportion of timepoints occupied by the state, (2) the appearance rate, the rate at which a consecutive sequence of state occupation appears, and (3) the dwell time and dwell variance, the mean and variation in durations of continuous occupations of the state^43^.

### Structured cycles

To investigate the temporal structure of whole-brain information states and their transitions, we turn to the recently developed temporal interval network density analysis (TINDA)^45^. We first partitioned all inter-state intervals in half and compute the FO asymmetry matrix:

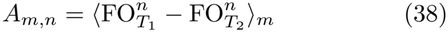

where *T*_1_ are the timepoints in the first half of the interval, *T*_2_ the second, where ⟨… ⟩_*m*_ denotes the average FO asymmetry for state *n* over reference state *m*’s intervals. Put simply, a positive value at index *A*_*m,n*_ indicates that state *n* occurs more dominantly succeeding rather than preceding state *m*. The same is true for the inverse: a negative value indicates that state *n* occupies a greater proportion of the period prerather than post-state *m*^45^. To discover the natural ordering of states in the cycle, we position nodes equidistantly on a unit circle spanning [0, 2*π*], thus providing each node an associated phase *q*. The transition from state *n* to *m* can be represented as a vector in the complex plane of phase differences 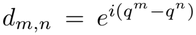. Treating *A* as a weighted directed graph we can scale the vectors according to the strength (and direction) of the corresponding FO asymmetry. The sum of the imaginary components then comes to represent the overall level of direction or broken detailed balance in the transition structure^45^. Accordingly, we have the cycle strength as:

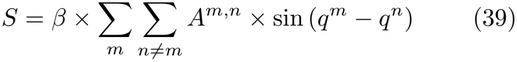

where *β* is a normalization factor based on the theoretical maximum cycle strength for *K* states, such that *S* is constrained to [−1, 1]. The polarity of *S* represents the net direction, being positive for clockwise flow and negative for counterclockwise flow. The 9!*/*2 distinct circular orderings of *K* = 10 states were searched by a stochastic hill-climber that proposes random transpositions of two nodes and accepts only those that increase *S*. Because such a search could potentially settle in a local optimum, we restarted it 10 times with 5 × 10^4^ proposals per restart and kept the ordering with the largest *S*; within each null permutation the optimiser was run with 2 × 10^4^ proposals^45^.

To assess the significance of the observed cycle strength, we generated null distributions of 1000 permutations, each independently relabelling the states within every subject, re-optimising the ordering, and recomputing *S*.

### Cycle synchrony

Assigning each state *k* the phase *q*^*k*^ of the cycle, we mapped each subject’s state sequence into a unit-modulus phase time series 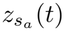, equal to 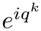 whenever subject *s*_*a*_ occupied state *k* at time *t*, truncated to the shortest film length *T* . The synchrony between two subjects *s*_*a*_ and *s*_*b*_ is simply their time-averaged phase-locking value

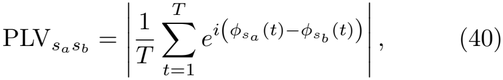

where 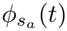(*t*) is the phase of 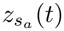, so that 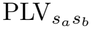 approaches 1 when two cycle-phase time courses are perfectly locked and 0 when their phase difference is uniform over time.

To assess the significance of the observed synchrony, we generated a null distribution of PLVs by circularly shifting each subject’s phase time series by a random temporal offset (1000 surrogates), preserving each subject’s marginal dynamics and autocorrelation.

### Multimodal stimulus extraction

To quantitatively parse the content of the movie stimuli, we performed a window manifest partition of each film into consecutive non-overlapping core windows of 1 s, each nested within a 3 s context window (±1 TR). Three annotation streams were generated for every window. First, *speech*: the soundtrack was tran-scribed with Whisper large-v3 (faster-whisper, beam size 5: github.com/SYSTRAN/faster-whisper)^133^, returning word-level onsets, offsets and posterior probabilities, and speaker turns were recovered by diarisation^134^. Each word was assigned to the window containing its onset, yielding a per-TR dialogue string, together with a wider dialogue string spanning the context window. Second, *vision*: eight frames sampled uniformly across the core window were passed to a vision-language model (`Qwen3-VL-8B-Instruct`)^135^, instructed to describe only what was directly visible and not to infer plot, identity, relationships or intentions, under a fixed set of eight fields (actions, alcohol, objects, people, setting, shot, text, and uncertain). Third, *sound* : an audio-language model (`Qwen2.5-Omni-7B`)^136^ was queried twice per window, once on the bare core clip for discrete non-speech sound events, and once on the core clip padded by 3.75s on either side for the slower-varying ambience and music, in both cases instructed to ignore speech. Field-structured captions were parsed back into their slots, and the concatenation of the content-bearing slots forms the per-TR *embedding text* of that stream.

Quality control was performed per stream and per window. Windows were flagged as failed when the caption of a stream could not be parsed into its fields, when the frame yield fell below 0.75, or when the resulting caption was empty; the union of these flags forms a per-film failed mask. For all analyses we additionally required that a TR carry a non-degenerate embedding and a non-empty caption, which removes title cards, black frames and other content-free intervals that would otherwise occupy an arbitrary position in the semantic space. Stimulus annotations were aligned to the fMRI data by pairing “brain time” *t* with the annotation of the film at *t* − 5 s. Because films differ in genre, pace and dialogue density, every stimulus feature was standardised within film before pooling, and, for the embedding analyses, the film mean was subtracted from each embedding, so that no reported effect can be an expression of film identity.

Statistical inference throughout used a participant-wise circular-shift null: each participant’s state sequence was shifted by an independent random offset, which preserves the first order statistics of the sequence. Observed and null statistics were pooled identically, so any shrinkage induced by averaging over hundreds of thousands of TRs is common to both. Unless stated otherwise, nulls comprise 5000 permutations and p-values are two-tailed and FDR-corrected by the Benjamini-Hochberg procedure.

### Normative profiles

To explore what kind of psycholinguistic content occurs during each mode of global information dynamics, we scored every TR against a battery of lexical norms. We used 24 dimensions drawn from eight normative corpora: the eleven Lancaster sensorimotor norms (auditory, foot/leg, gustatory, hand/arm, haptic, head, interoceptive, mouth, olfactory, torso, visual)^137^; concreteness^138^; socialness^139^; the six factor scores of the higher-order semantic space (sub-lexical, distributional, visuotactile, body action, affective, social interaction)^140^; valence and arousal from the NRC valence-arousal-dominance lexicon^141^; cognition^142^; reward^143^: motivation^143^; and Zipf log word frequency^144^.

Each norm dimension *j* was first z-scored across the entirety of its source table, so that a score retains its meaning relative to the external norm rather than to this stimulus set. The embedding text of a TR (speech, visual and audio slots concatenated) was lower-cased and tokenised on non-alphanumeric boundaries, and the TR was scored as the mean table-wide *z* of whichever of its tokens appear in that table,

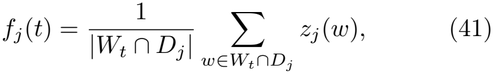

where *W*_*t*_ are the tokens of TR *t* and *D*_*j*_ the vocabulary of dimension *j*. Tokens absent from a table contribute nothing to that dimension and do not penalise the mean. The resulting *T* × 24 matrix was then *z*-scored column-wise within each film, with the rare undefined entries imputed to the column mean and therefore sitting exactly at zero after centring.

For each mode of information dynamics *a* and dimension *j* we accumulated the feature over every occurrence of that state, across all participants and films, and divided by the number of occurrences,

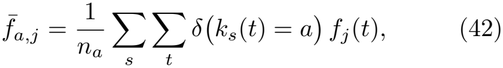

where *k*_*s*_(*t*) is the state of participant *s* at time *t* and *n*_*a*_ the total number of occurrences of state *a* (576,888 state occurrences in total). The same quantity was recomputed under the circular-shift null to give *z*-scores 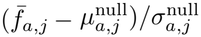 and permutation *p*-values, FDR-corrected over the state × dimension cells. An omnibus test per dimension, asking whether that dimension is distributed unevenly over states at all, used the between-state sum of squares 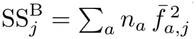 against the same null. Effects are reported in units of within-film standard deviations of the norm.

### Semantic embedding

To examine how global information dynamics are reflected in a more abstract semantic space, we created a multimodal embedding space. The embedding text of each stream was encoded independently with the multilingual text encoder (BGE-M3, dense-only mode)^145^, giving a unit-norm 1024-dimensional vector per stream per TR. The three streams were combined per TR with adaptive weights. Baseline weights were 0.7 for dialogue, 0.2 for vision and 0.1 for environmental audio. A salience gate then adjusted these per TR, raising the visual and environmental-audio weights when those streams were unusually salient. Each salience cue was robustly standardised within film and only positive deviations counted. The visual boost was a weighted sum of frame motion energy (0.08) and visual novelty, taken as one minus the cosine similarity between consecutive visual embeddings (0.06), capped at 0.15. The environmental boost was a weighted sum of the acoustic energy (0.06) and spectral centroid (0.04) of the core window and audio novelty (0.05), capped at 0.10. When more than 70% of the window’s energy was at high frequencies the environmental boost was multiplied by 0.3, which damps it in bright, broadband passages. Both boosts were taken out of the dialogue weight; if that drove the dialogue weight below a floor of 0.40 it was restored to the floor and the deficit reclaimed from the visual and environmental weights in proportion to their size. At TRs with no speech the dialogue weight was set to zero and the remaining two weights renormalised to sum to one, as were all weights after gating. Writing 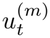 for the unit-norm embedding of stream *m* and 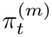 for its weight, the instantaneous embedding is 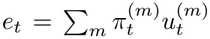, renormalised to unit length.

Because meaning at a given moment is carried partly by what has just happened, a temporal context was then applied. The context-weighted embedding places half of its weight on the present TR and distributes the remaining half over the preceding TRs with an exponential kernel,

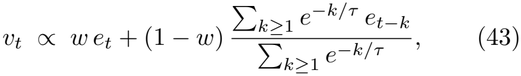

with *w* = 0.5 and *τ* = 9 s (half-life 6.24 s), renormalised to unit length. We retain both this context-weighted matrix and the raw per-TR matrix, the latter built from the baseline weights without the salience boosts; the former is used wherever the semantic content of a moment is at issue, the latter wherever the movement of meaning is at issue, since its exponential memory would otherwise impose autocorrelation on any measure of semantic change.

To see how global information dynamics explain semantic variance, within each film, embeddings were demeaned and the residualised vectors of all films pooled and reduced by principal component analysis to 100 components. With this, we computed a PERMANOVA-style partition of the total sum of squares about the grand mean into between-state and within-state components, summarised by

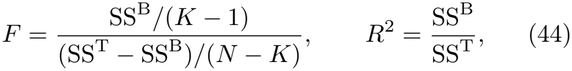

with *K* = 10 modes of information dynamics, and *N* the total number of state occurrences, tested against the circular-shift null. We then tested each pair of modes of information dynamics individually, taking the squared Euclidean distance between their centroids in this space as the statistic and correcting across pairs by FDR. As a diagnostic against occupancy-driven artefacts, the norm of each state’s centroid was also compared to its null, since a rarely visited state has a noisier and therefore longer centroid by construction.

### Semantic kinematics

To explore how information dynamics mirror semantic dynamics, we quantified properties of the stimulus’s trajectory through the semantic space, as opposed to its position in it, thus assaying how meaning evolved from moment to moment.

We conceived of semantic kinematics through the lens of four constructs: *Flux* is the overall rate at which meaning is changing, high when the film is moving quickly through semantic space and revisiting nothing. *Convergence* is the degree to which the film is settling, high when the is future more semantically coherent than the past. *Handover* is the degree to which the present moment is semantically closer to what comes next than what has come before. *Rush* is the excess of the current rate of change over its own recent neighbourhood, a local semantic acceleration.

For each construct we assembled a family of candidate measures of the local trajectory around each TR, 45 in total (26 for flux, 6 for convergence, 6 for handover, and 7 for rush). These were built from five primitives: the step size between consecutive states; the deviation of the current state from the mean of a neighbouring window; the distance to the state at a fixed lag; the coherence of a window (the norm of its mean unit vector, which approaches one when the film is parked in one region of semantic space); and recurrence, the distance to the nearest matching state within a lag band, whose lower bound of four TRs prevents the measure from rediscovering local smoothness rather than a genuine callback. Each primitive was taken over past and future windows spanning 3 to 120 TRs, together with the past-future asymmetries and local contrasts formed by differencing matched windows. Members were oriented by flipping the sign of any measure running opposite to its construct, coherence for instance entering flux negatively, and each was *z*-scored within film. All were computed on the raw per-TR embedding, which carries no built-in temporal memory, using cosine distance *d*(*x, y*) = 1 −cos(*x, y*) throughout. The full bank is given in Supplementary Table S1.

Each axis was then taken as the mean of its oriented members, and tested against states exactly as the norms were, giving *z*-scores and FDR-corrected permutation *p*-values over all state by axis cells.

Because the members of a family are distinct operationalisations of a single construct, they also serve as internal replications of the axis built from them: were an axis’s profile over states an artefact of averaging rather than a property of the construct, its individual measures would not reproduce it. We therefore re-tested every measure on its own. Since, a measure cannot be compared against a composite of which it is itself an ingredient, each measure was compared instead against the leave-one-out composite of its family, that is the mean of the other members. Members and their leave-one-out composites were carried through the same permutation pass, and agreement was quantified as the correlation across the ten states between a measure’s own *z*-profile and that of its leave-one-out composite.

### Replication

To demonstrate the robustness of our results, we employ our recently collected additional multimodal naturalistic neuroimaging dataset at 3T (NNDb-3T+)^146^.

#### Participants

In contrast to the original database, in which participants were distributed across ten films, every participant here watched the same feature-length film, Back to the Future. The dataset provides movie-watching scans from 40 participants (21 female; ages 18 to 45 years, mean 29.02 ± 6.20), predominantly right-handed native English speakers with no history of neurological illness. Seven participants were excluded for incomplete or interrupted runs, or a missing anatomical scan, retaining 33 participants for analysis. Each participant gave written consent for their anonymised data to be used and shared. The whole study including MRI was approved by the ethics committee at University College London (Reference number: fMRI/2023/003).

#### MRI acquisition

Functional and anatomical images were acquired on a 3T Siemens MAGNETOM Prisma scanner equipped with a custom 30 channel head coil (a standard 32 channel coil with the two eye elements removed to permit wide-angle viewing). The movie-watching protocol utilised a CMRR multiband EPI sequence^87^ (TR = 1.5 s, TE = 35.2 ms, 60^°^ flip angle, 72 interleaved slices, and 2 mm isotropic resolution), with a 4× multiband factor and anterior-to-posterior phase encoding. A reverse phase-encoded (posterior-to-anterior) scan was acquired after each run to permit susceptibility distortion correction. The film (duration 1 h 51 min) was presented in three consecutive runs of 1360, 1522 and 1608 volumes, the first 8, 16 and 16 volumes of which were discarded so that the retained time series matched the full film without overlap between runs. Following the functional scans, anatomical images were obtained using a 5-minute high-resolution T1-weighted MPRAGE sequence (TR = 2.3 s, TE = 2.98 ms, 208 sagittal slices, 1.0 mm resolution). During scanning, the film was displayed through a mirror-reversing LCD projector onto a 35.5 × 26 cm rear-projection screen, viewed through a mirror mounted on the head coil at a distance of 57.5 cm 28.9^°^ × 18.3^°^ visual angle. Sound was delivered through MRI-compatible noise-attenuating insert earphones, and head motion was further restrained with an inflatable head-immobilisation device.

#### Preprocessing

Anatomical images were skull-stripped and non-linearly aligned to the MNI152 template. This was followed by inflation, registration and segmentation using Freesurfer’s recon-all with standard settings^90^. Functional images were first stripped of the initial volumes of each run (8 from the first run and 16 from the second and third) and corrected for susceptibility-induced geometric distortion using the reverse phase-encoded field maps. EPI volumes were aligned to the volume with minimal motion by a two-pass registration, and then co-registered to their respective anatomical scans. These aligned time series were then warped to the MNI template space, and spatially smoothed with 4mm full-width half maximum^91^. The smoothed time series were then detrended with motion and ventricle regressors, together with a local white-matter regressor estimated by fast ANATICOR^147^. As for the original dataset, the concatenated time series were band-passed between 0.008–0.09 Hz to remove high-frequency noise and low-frequency drifts. To ensure the computational feasibility of our analysis, we parcellated the functional data by averaging all the voxels in an area prescribed by combining the 1000 cortical ROIs from the Schaefer atlas^94^, and the 54 subcortical ROIs from the Tian atlas^95^.

## V. DATA AVAILABILITY

Full-length movie watching fMRI data is publicly available at naturalistic-neuroimaging-database. org. The replication movie watching 3T fMRI dataset has been deposited on OpenNeuro at openneuro.org/datasets/ds006642. HCP DWI data is publicly available at brain.labsolver.org. AHBA mRNA data is publicly available at human.brain-map.org.

## VI. CODE AVAILABILITY

Code will be made publicly available upon acceptance of the manuscript for publication at github.com/gblackburne/natural-info.

## VII. ACKNOWLEDGEMENTS

All MRI scanning hours were donated by the Birkbeck-UCL Centre for NeuroImaging (BUCNI) as part of its commitment to open science resources. We would like to thank all staff at BUCNI for their support. G.B is supported by the Leverhulme Trust.

## VIII. CONTRIBUTIONS

G.B.: Conceptualization, Methodology, Software, Formal analysis, Data curation, Validation, Visualization, Project administration, Writing – original draft, Writing – review & editing. A.L.: Methodology, Software, Writing – review & editing. A.B.: Writing – review & editing. E.L.: Investigation, Writing – review & editing. G.C.: Writing – review & editing. F.D.: Investigation, Writing – review & editing. A.I.L.: Writing – review & editing. J.I.S.: Investigation, Resources, Supervision, Writing – review & editing. P.A.M.M.: Supervision, Writing – review & editing.

## IX. ETHICS DECLARATIONS

### Competing interests

The authors declare no competing interests.

## X. CITATION DIVERSITY

Recent work in several fields of science has identified a bias in citation practices such that papers from marginalised demographics are under-cited relative to the number of such papers in the field^148–150^. To report transparently on our own bias, first, we classified the gender of the first names of the first and last authors^148^. Our references contain 6.99% woman(first)/woman(last), 4.9% man/woman, 16.53% woman/man, and 71.59% man/man. Second, we performed a similar procedure for obtaining predicted racial or ethnic category. By this measure, splitting into inferred white authors (W) and authors of colour (C), our references contain 17.05% C/C, 16.46% W/C, 26.01% C/W, and 40.48% W/W. We note that these algorithmic proxies are limited, cannot infer the full spectrum of self-determined gender identity, structurally exclude non-binary, transgender, and intersex researchers, and cannot adequately account for Indigenous and mixed-race authors. We look forward to future work that could help us to better understand how to support equitable practices in science^151^.

## Notes

### Competing Interest Statement

The authors have declared no competing interest.

## References

1 Anderson, J. R. The Architecture of Cognition (Harvard Univ. Press, Cambridge, MA, USA, 1983).

2 McClelland, J. L. & Rumelhart, D. E. Parallel Distributed Processing (MIT Press, Cambridge, MA, USA, 1987).

3 Mesulam, M. M. Large-scale neurocognitive networks and distributed processing for attention, language, and memory. Ann. Neurol. 28, 597–613 (1990).

4 Mediano, P. A. M. et al. Toward a unified taxonomy of information dynamics via integrated information decomposition. Proc. Natl. Acad. Sci. U. S. A. 122, e2423297122 (2025).

5 Williams, P. L. & Beer, R. D. Nonnegative decomposition of multivariate information. arXiv [cs.IT] (2010). 1004.2515.

6 Gawne, T. J. & Richmond, B. J. How independent are the messages carried by adjacent inferior temporal cortical neurons? J. Neurosci. 13, 2758–2771 (1993).

7 McGill, W. J. Multivariate information transmission. Psychometrika 19, 97–116 (1954).

8 Luppi, A. I., Rosas, F. E., Mediano, P. A. M., Menon, D. K. & Stamatakis, E. A. Information decomposition and the informational architecture of the brain. Trends Cogn. Sci. 28, 352–368 (2024).

9 Nigam, S., Pojoga, S. & Dragoi, V. Synergistic coding of visual information in columnar networks. Neuron 104, 402–411.e4 (2019).

10 Varley, T. F., Sporns, O., Schaffelhofer, S., Scherberger, H. & Dann, B. Information-processing dynamics in neural networks of macaque cerebral cortex reflect cognitive state and behavior. Proc. Natl. Acad. Sci. U. S. A. 120, e2207677120 (2023).

11 Gelens, F. et al. Distributed representations of prediction error signals across the cortical hierarchy are synergistic. Nat. Commun. 15, 3941 (2024).

12 Greco, A., Moser, J., Preissl, H. & Siegel, M. Predictive learning shapes the representational geometry of the human brain. Nat. Commun. 15, 9670 (2024).

13 Combrisson, E. et al. Higher-order and distributed synergistic functional interactions encode information gain in goal-directed learning. Nat. Commun. 16, 7179 (2025).

14 Proca, A. M. et al. Synergistic information supports modality integration and flexible learning in neural networks solving multiple tasks. PLoS Comput. Biol. 20, e1012178 (2024).

15 Barack, D. L. & Krakauer, J. W. Two views on the cognitive brain. Nat. Rev. Neurosci. 22, 359–371 (2021).

16 Luppi, A. I. et al. A synergistic core for human brain evolution and cognition. Nat. Neurosci. 25, 771–782 (2022).

17 Varley, T. F., Pope, M., Puxeddu, M. G., Faskowitz, J. & Sporns, O. Partial entropy decomposition reveals higher-order information structures in human brain activity. Proc. Natl. Acad. Sci. U. S. A. 120, e2300888120 (2023).

18 Margulies, D. S. et al. Situating the default-mode network along a principal gradient of macroscale cortical organization. Proc. Natl. Acad. Sci. U. S. A. 113, 12574–12579 (2016).

19 Jo, Y. et al. The diversity and multiplexity of edge communities within and between brain systems. Cell Rep. 37, 110032 (2021).

20 Vidaurre, D. et al. Spontaneous cortical activity transiently organises into frequency specific phase-coupling networks. Nat. Commun. 9, 2987 (2018).

21 Dimakou, A., Pezzulo, G., Zangrossi, A. & Corbetta, M. The predictive nature of spontaneous brain activity across scales and species. Neuron 113, 1310–1332 (2025).

22 Luppi, A. I., Gellersen, H. M., Naci, L., Gozzi, A. & Bor, D. Unlocking spontaneous cognition in the age of data science. Nat. Rev. Neurosci. 27, 591–593 (2026).

23 Biswal, B. B. & Uddin, L. Q. The history and future of resting-state functional magnetic resonance imaging. Nature 641, 1121–1131 (2025).

24 Finn, E. S. Is it time to put rest to rest? Trends Cogn. Sci. 25, 1021–1032 (2021).

25 Aliko, S., Huang, J., Gheorghiu, F., Meliss, S. & Skipper, J. I. A naturalistic neuroimaging database for understanding the brain using ecological stimuli. Sci. Data 7, 347 (2020).

26 Wang, J. et al. Test-retest reliability of functional connectivity networks during naturalistic fMRI paradigms. Hum. Brain Mapp. 38, 2226–2241 (2017).

27 Vanderwal, T. et al. Individual differences in functional connectivity during naturalistic viewing conditions. Neuroimage 157, 521–530 (2017).

28 Finn, E. S. & Bandettini, P. A. Movie-watching outperforms rest for functional connectivity-based prediction of behavior. Neuroimage 235, 117963 (2021).

29 Wang, L., Mruczek, R. E. B., Arcaro, M. J. & Kastner, S. Probabilistic maps of visual topography in human cortex. Cereb. Cortex 25, 3911–3931 (2015).

30 Yeo, B. T. T. et al. The organization of the human cerebral cortex estimated by intrinsic functional connectivity. J. Neurophysiol. 106, 1125–1165 (2011).

31 Luppi, A. I. et al. Unravelling consciousness and brain function through the lens of time, space, and information. Trends Neurosci. 47, 551–568 (2024).

32 Liardi, A., Blackburne, G. & Mediano, P. A. M. An information-geometric formulation of distributed computation. arXiv [cs.IT] (2026). 2611.03545.

33 Liu, Z.-Q. et al. Benchmarking methods for mapping functional connectivity in the brain. Nat. Methods 22, 1593–1602 (2025).

34 Blackburne, G. et al. Psychedelics restructure information in the human brain. bioRxiv 2026.10.01.755915 (2026).

35 Bassett, D. S. & Sporns, O. Network neuroscience. Nat. Neurosci. 20, 353–364 (2017).

36 Sporns, O. Contributions and challenges for network models in cognitive neuroscience. Nat. Neurosci. 17, 652–660 (2014).

37 Seguin, C., Sporns, O. & Zalesky, A. Brain network communication: concepts, models and applications. Nat. Rev. Neurosci. 24, 557–574 (2023).

38 Zamani Esfahlani, F., Faskowitz, J., Slack, J., Mišić, B. & Betzel, R. F. Local structure-function relationships in human brain networks across the lifespan. Nat. Commun. 13, 2053 (2022).

39 Milisav, F., Bazinet, V., Betzel, R. F. & Misic, B. A simulated annealing algorithm for randomizing weighted networks. Nat. Comput. Sci. 5, 48–64 (2025).

40 Zalesky, A., Fornito, A., Cocchi, L., Gollo, L. L. & Breaks-pear, M. Time-resolved resting-state brain networks. Proc. Natl. Acad. Sci. U. S. A. 111, 10341–10346 (2014).

41 Shine, J. M. et al. Human cognition involves the dynamic integration of neural activity and neuromodulatory systems. Nat. Neurosci. 22, 289–296 (2019).

42 Betzel, R. F., Faskowitz, J. & Sporns, O. Living on the edge: network neuroscience beyond nodes. Trends Cogn. Sci. 27, 1068–1084 (2023).

43 Vidaurre, D., Smith, S. M. & Woolrich, M. W. Brain network dynamics are hierarchically organized in time. Proc. Natl. Acad. Sci. U. S. A. 114, 12827–12832 (2017).

44 Mediano, P. A. M. et al. Spectrally and temporally resolved estimation of neural signal diversity. eLife (2023).

45 van Es, M. W. J. et al. Large-scale cortical functional networks are organized in structured cycles. Nat. Neurosci. 1–11 (2025).

46 Ecker, A. S. et al. Decorrelated neuronal firing in cortical microcircuits. Science 327, 584–587 (2010).

47 Schneidman, E., Berry, M. J., 2nd, Segev, R. & Bialek, W. Weak pairwise correlations imply strongly correlated network states in a neural population. Nature 440, 1007–1012 (2006).

48 Averbeck, B. B., Latham, P. E. & Pouget, A. Neural correlations, population coding and computation. Nat. Rev. Neurosci. 7, 358–366 (2006).

49 Panzeri, S., Moroni, M., Safaai, H. & Harvey, C. D. The structures and functions of correlations in neural population codes. Nat. Rev. Neurosci. 23, 551–567 (2022).

50 Denève, S. & Machens, C. K. Efficient codes and balanced networks. Nat. Neurosci. 19, 375–382 (2016).

51 Munn, B. R. et al. Multiscale organization of neuronal activity unifies scale-dependent theories of brain function. Cell 187, 7303–7313.e15 (2024).

52 Bechtel, W. & Richardson, R. C. Discovering Complexity: Decomposition and localization as strategies in scientific research (The MIT Press, 2010).

53 McCaffrey, J. B. Evolving concepts of functional localization. Philos. Compass 18, e12914 (2023).

54 Rosen, M. C. & Freedman, D. J. How distributed is the brain-wide network that is recruited for cognition? Nat. Rev. Neurosci. 27, 138–150 (2026).

55 Hayden, B. Y., Heilbronner, S. R. & Yoo, S. B. M. Rethinking the centrality of brain areas in understanding functional organization. Nat. Neurosci. 29, 267–278 (2026).

56 Kanwisher, N. Functional specificity in the human brain: a window into the functional architecture of the mind. Proc. Natl. Acad. Sci. U. S. A. 107, 11163–11170 (2010).

57 Lashley, K. S. Brain mechanisms and intelligence (University of Chicago Press, Chicago, 1929).

58 Op de Beeck, H. P., Haushofer, J. & Kanwisher, N. G. Interpreting fMRI data: maps, modules and dimensions. Nat. Rev. Neurosci. 9, 123–135 (2008).

59 International Brain Laboratory et al. A brain-wide map of neural activity during complex behaviour. Nature 645, 177–191 (2025).

60 Anderson, M. L., Kinnison, J. & Pessoa, L. Describing functional diversity of brain regions and brain networks. Neuroimage 73, 50–58 (2013).

61 Mundale, J. Concepts of localization: Balkanization in the brain. Brain Mind 3, 313–330 (2002).

62 Huntenburg, J. M.Bazin, P.-L. & Margulies, D. S. Large-scale gradients in human cortical organization. Trends Cogn. Sci. 22, 21–31 (2018).

63 Sydnor, V. J. et al. Neurodevelopment of the association cortices: Patterns, mechanisms, and implications for psychopathology. Neuron 109, 2820–2846 (2021).

64 Wang, X.-J. Theory of the multiregional neocortex: Large-scale neural dynamics and distributed cognition. Annu. Rev. Neurosci. 45, 533–560 (2022).

65 Dehaene, S., Kerszberg, M. & Changeux, J. P. A neuronal model of a global workspace in effortful cognitive tasks. Proc. Natl. Acad. Sci. U. S. A. 95, 14529–14534 (1998).

66 Barbas, H. General cortical and special prefrontal connections: principles from structure to function. Annu. Rev. Neurosci. 38, 269–289 (2015).

67 Hilgetag, C. C. & Goulas, A. ‘hierarchy’ in the organization of brain networks. Philos. Trans. R. Soc. Lond. B Biol. Sci. 375, 20190319 (2020).

68 Chanes, L. & Barrett, L. F. Redefining the role of limbic areas in cortical processing. Trends Cogn. Sci. 20, 96–106 (2016).

69 Deco, G., Jirsa, V. K., Robinson, P. A., Breakspear, M. & Friston, K. The dynamic brain: from spiking neurons to neural masses and cortical fields. PLoS Comput. Biol. 4, e1000092 (2008).

70 Kumar, A., Rotter, S. & Aertsen, A. Spiking activity propagation in neuronal networks: reconciling different perspectives on neural coding. Nat. Rev. Neurosci. 11, 615–627 (2010).

71 Avena-Koenigsberger, A., Misic, B. & Sporns, O. Communication dynamics in complex brain networks. Nat. Rev. Neurosci. 19, 17–33 (2017).

72 Suárez, L. E., Markello, R. D., Betzel, R. F. & Misic, B. Linking structure and function in macroscale brain networks. Trends Cogn. Sci. 24, 302–315 (2020).

73 Fakhar, K. et al. A general framework for characterizing optimal communication in brain networks. Elife 13, RP101780 (2025).

74 Anderson, M. L. After phrenology: Neural reuse and the interactive brain (The MIT Press, 2014).

75 Dehaene, S. & Cohen, L. Cultural recycling of cortical maps. Neuron 56, 384–398 (2007).

76 Hebb, D. O. The organization of behaviour (John Wiley & Sons, Nashville, TN, 1949), 99 edn.

77 Buzsáki, G. Neural syntax: cell assemblies, synapsembles, and readers. Neuron 68, 362–385 (2010).

78 Ikegaya, Y. et al. Synfire chains and cortical songs: temporal modules of cortical activity. Science 304, 559–564 (2004).

79 Luczak, A., Barthó, P. & Harris, K. D. Spontaneous events outline the realm of possible sensory responses in neocortical populations. Neuron 62, 413–425 (2009).

80 Xie, W. et al. Neuronal sequences in population bursts encode information in human cortex. Nature 635, 935–942 (2024).

81 Driscoll, L. N., Shenoy, K. & Sussillo, D. Flexible multitask computation in recurrent networks utilizes shared dynamical motifs. Nat. Neurosci. 27, 1349–1363 (2024).

82 Huth, A. G., de Heer, W. A., Griffiths, T. L., Theunissen, F. E. & Gallant, J. L. Natural speech reveals the semantic maps that tile human cerebral cortex. Nature 532, 453–458 (2016).

83 Lurie, D. J. et al. Questions and controversies in the study of time-varying functional connectivity in resting fMRI. Netw. Neurosci. 4, 30–69 (2020).

84 Faskowitz, J., Esfahlani, F. Z., Jo, Y., Sporns, O. & Betzel, R. F. Edge-centric functional network representations of human cerebral cortex reveal overlapping system-level architecture. Nat. Neurosci. 23, 1644–1654 (2020).

85 Novelli, L. & Razi, A. A mathematical perspective on edge-centric brain functional connectivity. Nat Commun 13, 2693 (2022).

86 Yartsev, M. M. Natural behavior is what you do, not just where you are. Nat. Neurosci. 29, 763–766 (2026).

87 Feinberg, D. A. & Setsompop, K. Ultra-fast MRI of the human brain with simultaneous multi-slice imaging. J. Magn. Reson. 229, 90–100 (2013).

88 Todd, N. et al. Evaluation of 2D multiband EPI imaging for high-resolution, whole-brain, task-based fMRI studies at 3T: Sensitivity and slice leakage artifacts. Neuroimage 124, 32–42 (2016).

89 Cauley, S. F., Polimeni, J. R., Bhat, H., Wald, L. L. & Setsompop, K. Interslice leakage artifact reduction technique for simultaneous multislice acquisitions: Interslice leakage artifact reduction technique. Magn. Reson. Med. 72, 93–102 (2014).

90 Fischl, B. FreeSurfer. Neuroimage 62, 774–781 (2012).

91 Friedman, L., Glover, G. H., Krenz, D., Magnotta, V. & FIRST BIRN. Reducing inter-scanner variability of activation in a multicenter fMRI study: role of smoothness equalization. Neuroimage 32, 1656–1668 (2006).

92 Caballero-Gaudes, C. & Reynolds, R. C. Methods for cleaning the BOLD fMRI signal. Neuroimage 154, 128–149 (2017).

93 Smith, S. M. et al. Advances in functional and structural MR image analysis and implementation as FSL. Neuroimage S208–19 (2004).

94 Schaefer, A. et al. Local-global parcellation of the human cerebral cortex from intrinsic functional connectivity MRI. Cereb. Cortex 28, 3095–3114 (2018).

95 Tian, Y., Margulies, D. S., Breakspear, M. & Zalesky, A. Topographic organization of the human subcortex unveiled with functional connectivity gradients. Nat. Neurosci. 23, 1421–1432 (2020).

96 Van Essen, D. C. et al. The WU-minn human connectome project: an overview. Neuroimage 80, 62–79 (2013).

97 Glasser, M. F. et al. The minimal preprocessing pipelines for the human connectome project. Neuroimage 80, 105–124 (2013).

98 Sotiropoulos, S. N. et al. Advances in diffusion MRI acquisition and processing in the human connectome project. Neuroimage 80, 125–143 (2013).

99 Yeh, F.-C. et al. Population-averaged atlas of the macroscale human structural connectome and its network topology. Neuroimage 178, 57–68 (2018).

100 Yeh, F.-C. Population-based tract-to-region connectome of the human brain and its hierarchical topology. Nat. Commun. 13, 4933 (2022).

101 Yeh, F.-C. & Tseng, W.-Y. I. NTU-90: a high angular resolution brain atlas constructed by q-space diffeomorphic reconstruction. Neuroimage 58, 91–99 (2011).

102 Yeh, F.-C., Verstynen, T. D., Wang, Y., Fernández-Miranda, J. C. & Tseng, W.-Y. I. Deterministic diffusion fiber tracking improved by quantitative anisotropy. PLoS One 8, e80713 (2013).

103 Braunstein, S. L., Ghosh, S. & Severini, S. The laplacian of a graph as a density matrix: A basic combinatorial approach to separability of mixed states. Ann. Comb. 10, 291–317 (2006).

104 De Domenico, M., Nicosia, V., Arenas, A. & Latora, V. Structural reducibility of multilayer networks. Nat. Commun. 6, 6864 (2015).

105 Li, L., Alderson, D., Tanaka, R., Doyle, J. C. & Willinger, W. Towards a theory of scale-free graphs: Definition, properties, and implications (extended version). arXiv [cond-mat.dis-nn] (2005). cond-mat/0501169.

106 Newman, M. E. J. Equivalence between modularity optimization and maximum likelihood methods for community detection. Phys Rev E 94, 052315 (2016).

107 Jeub, L. G. S., Sporns, O. & Fortunato, S. Multiresolution consensus clustering in networks. Sci. Rep. 8, 3259 (2018).

108 Traag, V. A., Waltman, L. & van Eck, N. J. From louvain to leiden: guaranteeing well-connected communities. Sci. Rep. 9, 5233 (2019).

109 De Domenico, M. More is different in real-world multilayer networks. Nat. Phys. 19, 1247–1262 (2023).

110 Latora, V. & Marchiori, M. Efficient behavior of small-world networks. Phys. Rev. Lett. 87, 198701 (2001).

111 Boguñá, M., Krioukov, D. & Claffy, K. C. Navigability of complex networks. Nat. Phys. 5, 74–80 (2009).

112 Seguin, C., van den Heuvel, M. P. & Zalesky, A. Navigation of brain networks. Proc. Natl. Acad. Sci. U. S. A. 115, 6297–6302 (2018).

113 Estrada, E. & Hatano, N. Communicability in complex networks. Phys. Rev. E Stat. Nonlin. Soft Matter Phys. 77, 036111 (2008).

114 Zhou, H. Network landscape from a brownian particle’s perspective. Phys. Rev. E Stat. Nonlin. Soft Matter Phys. 67, 041908 (2003).

115 Goñi, J. et al. Exploring the morphospace of communication efficiency in complex networks. PLoS One 8, e58070 (2013).

116 Finn, E. S. et al. Functional connectome fingerprinting: identifying individuals using patterns of brain connectivity. Nat. Neurosci. 18, 1664–1671 (2015).

117 Amico, E. & Goñi, J. The quest for identifiability in human functional connectomes. Sci. Rep. 8, 8254 (2018).

118 Tolle, H. M. et al. The unique neural signature of your trip: Functional connectome fingerprints of subjective psilocybin experience. Netw. Neurosci. 8, 203–225 (2024).

119 Hawrylycz, M. J. et al. An anatomically comprehensive atlas of the adult human brain transcriptome. Nature 489, 391–399 (2012).

120 Arnatkeviciute, A., Fulcher, B. D. & Fornito, A. A practical guide to linking brain-wide gene expression and neuroimaging data. Neuroimage 189, 353–367 (2019).

121 Quackenbush, J. Microarray data normalization and trans-formation. Nat. Genet. 32 Suppl, 496–501 (2002).

122 Hawrylycz, M. et al. Canonical genetic signatures of the adult human brain. Nat. Neurosci. 18, 1832–1844 (2015).

123 Fulcher, B. D., Little, M. A. & Jones, N. S. Highly comparative time-series analysis: the empirical structure of time series and their methods. J. R. Soc. Interface 10, 20130048 (2013).

124 Wagstyl, K. et al. Transcriptional cartography integrates multiscale biology of the human cortex. eLife (2024).

125 Burt, J. B. et al. Hierarchy of transcriptomic specialization across human cortex captured by structural neuroimaging topography. Nat. Neurosci. 21, 1251–1259 (2018).

126 Fagerberg, L. et al. Analysis of the human tissue-specific expression by genome-wide integration of transcriptomics and antibody-based proteomics. Mol. Cell. Proteomics 13, 397–406 (2014).

127 Genovese, G. et al. Increased burden of ultra-rare protein-altering variants among 4,877 individuals with schizophrenia. Nat. Neurosci. 19, 1433–1441 (2016).

128 Cahoy, J. D. et al. A transcriptome database for astrocytes, neurons, and oligodendrocytes: a new resource for understanding brain development and function. J. Neurosci. 28, 264–278 (2008).

129 Pirooznia, M. et al. SynaptomeDB: an ontology-based knowledgebase for synaptic genes. Bioinformatics 28, 897–899 (2012).

130 Yarkoni, T., Poldrack, R. A., Nichols, T. E., Van Essen, D. C. & Wager, T. D. Large-scale automated synthesis of human functional neuroimaging data. Nat. Methods 8, 665–670 (2011).

131 Poldrack, R. A. et al. The cognitive atlas: toward a knowledge foundation for cognitive neuroscience. Front. Neuroinform. 5, 17 (2011).

132 Luppi, A. I. et al. General anaesthesia decreases the uniqueness of brain functional connectivity across individuals and species. Nat. Hum. Behav. 9, 987–1004 (2025).

133 Radford, A. et al. Robust speech recognition via large-scale weak supervision. arXiv [eess.AS] (2022). 2212.04356.

134 Bredin, H. et al. Pyannote.audio: Neural building blocks for speaker diarization. In ICASSP 2020 - 2020 IEEE International Conference on Acoustics, Speech and Signal Processing (ICASSP), 7124–7128 (IEEE, 2020).

135 Bai, S. et al. Qwen3-VL technical report. arXiv [cs.CV] (2025). 2511.21631.

136 Xu, J. et al. Qwen2.5-omni technical report. arXiv [cs.CL] (2025). 2503.20215.

137 Lynott, D., Connell, L., Brysbaert, M., Brand, J. & Carney, J. The lancaster sensorimotor norms: multidimensional measures of perceptual and action strength for 40,000 english words. Behav. Res. Methods 52, 1271–1291 (2020).

138 Brysbaert, M., Warriner, A. B. & Kuperman, V. Concreteness ratings for 40 thousand generally known english word lemmas. Behav. Res. Methods 46, 904–911 (2014).

139 Diveica, V., Pexman, P. M. & Binney, R. J. Quantifying social semantics: An inclusive definition of socialness and ratings for 8388 english words. Behav. Res. Methods 55, 461–473 (2023).

140 Diveica, V., Muraki, E. J., Binney, R. J. & Pexman, P. M. Mapping semantic space: Exploring the higher-order structure of word meaning. Cognition 248, 105794 (2024).

141 Mohammad, S. Obtaining reliable human ratings of valence, arousal, and dominance for 20,000 english words. In Proceedings of the 56th Annual Meeting of the Association for Computational Linguistics (Volume 1: Long Papers), 174–184 (Association for Computational Linguistics, Stroudsburg, PA, USA, 2018).

142 Corenblum, H. T. & Pexman, P. M. Cognition ratings for 8826 english words. Behav. Res. Methods 57, 181 (2025).

143 Giurgea, D.-I., Pexman, P. M. & Binney, R. J. Reward as a facet of word meaning: Ratings of motivation for 8,601 english words. Behav. Res. Methods 57, 242 (2025).

144 van Heuven, W. J. B., Mandera, P., Keuleers, E. & Brysbaert, M. SUBTLEX-UK: a new and improved word frequency database for british english. Q. J. Exp. Psychol. (Hove) 67, 1176–1190 (2014).

145 Chen, J. et al. M3-embedding: Multi-linguality, multi-functionality, multi-granularity text embeddings through self-knowledge distillation. arXiv [cs.CL] (2024). 2402.03216.

146 Levchenko, E. et al. A neuroimaging dataset combining movie-watching, eye-tracking, sensorimotor mapping, and cognitive tasks. Sci. Data 1–19 (2026).

147 Jo, H. J., Saad, Z. S., Simmons, W. K., Milbury, L. A. & Cox, R. W. Mapping sources of correlation in resting state FMRI, with artifact detection and removal. Neuroimage 52, 571–582 (2010).

148 Dworkin, J. D. et al. The extent and drivers of gender imbalance in neuroscience reference lists. Nat. Neurosci. 23, 918–926 (2020).

149 Caplar, N., Tacchella, S. & Birrer, S. Quantitative evaluation of gender bias in astronomical publications from citation counts. Nat. Astron. 1, 0141 (2017).

150 Dion, M. L., Sumner, J. L. & Mitchell, S. M. Gendered citation patterns across political science and social science methodology fields. Polit. Anal. 26, 312–327 (2018).

151 Dworkin, J., Zurn, P. & Bassett, D. S. (in)citing action to realize an equitable future. Neuron 106, 890–894 (2020).

