## Supplementary Material for "The information architecture of the naturalistic human brain"

#### Supplementary Material
### The information architecture of the naturalistic human brain

George Blackburne,[1,2,*] Alberto Liardi,[2] Andreea Bacalum,[2] Egor Levchenko,[3,4] Greg Cooper,[5] Fred Dick,[1,6] Andrea I. Luppi,[7,8,9,10] Jeremy I. Skipper,[1,†] and Pedro A. M. Mediano[2,11,†]

[1] *Department of Experimental Psychology, University College London*
[2] *Department of Computing, Imperial College London*
[3] *Department of Psychological Sciences, Birkbeck, University of London*
[4] *Department of Cell and Developmental Biology, University College London*
[5] *Department of Clinical, Educational and Health Psychology, University College London*
[6] *Birkbeck-UCL Centre for Neuroimaging, University College London*
[7] *Centre for Eudaimonia and Human Flourishing, University of Oxford*
[8] *Department of Psychiatry, University of Oxford*
[9] *Montreal Neurological Institute, McGill University*
[10] *Division of Information Engineering and St John's College, University of Cambridge*
[11] *Division of Psychology and Language Sciences, University College London*

#### CONTENTS

S1. Extended Data

S2. Supplementary Note 1: Complexity and diversity

S3. Supplementary Note 2: Replication

S4. Supplementary Methods

S5. Supplementary Figures

S6. Supplementary Tables

References

#### S1. EXTENDED DATA

Extended Data Figs. 1–6 repeat Figs. 1–6 of the main text in the independent NNDb-3T+ cohort ($n = 33$; see Methods). Captions follow those of the corresponding main-text figures, and departures from them are noted where they occur. The quantitative agreement between the two cohorts is reported in Supplementary Note 2 and Supplementary Figs. S16–S25.

† These authors jointly supervised this work.

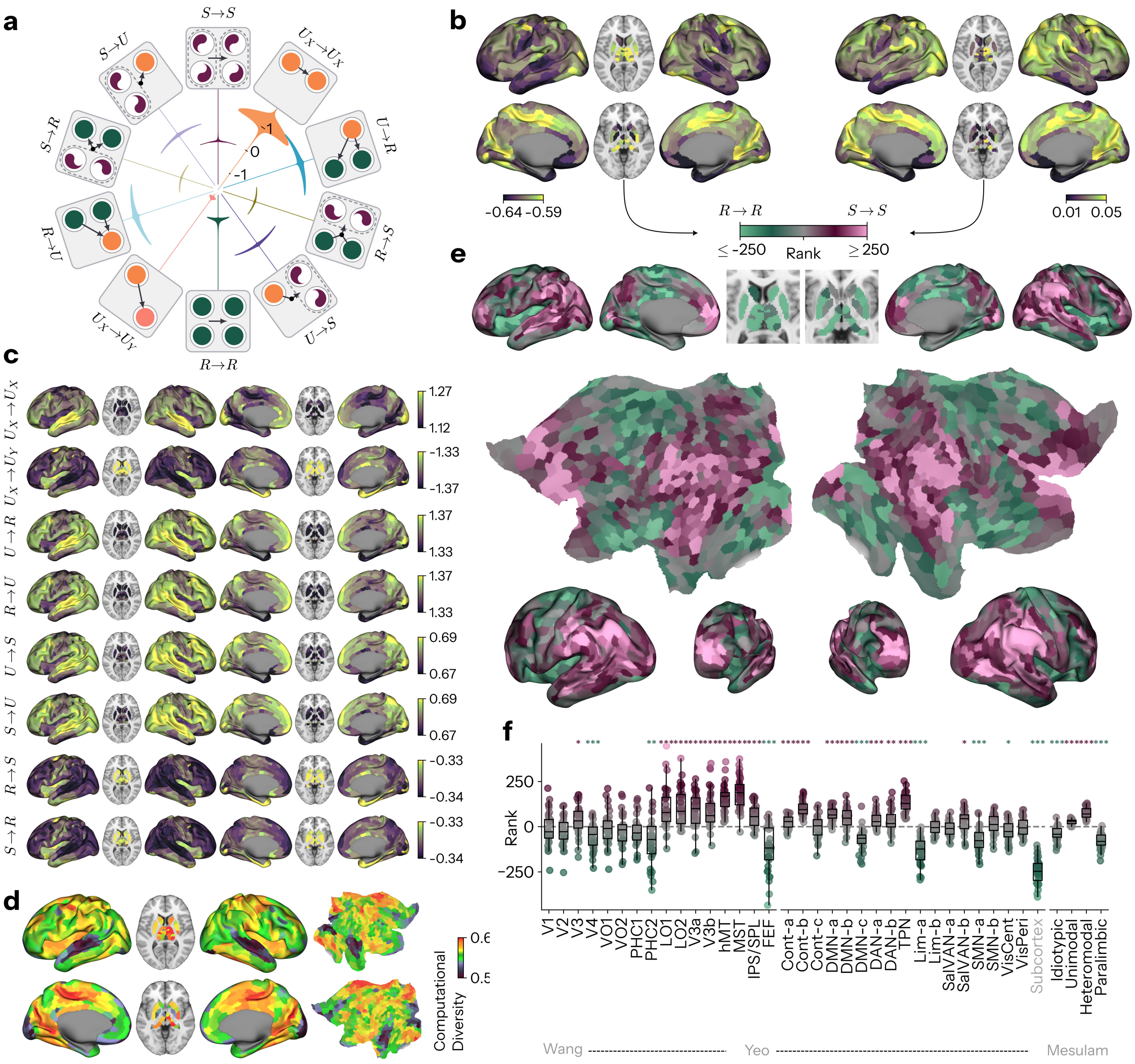


Extended Data Fig. 1. **Replication of the information architecture of the naturalistic human brain.** As Fig. 1, in the NNDb-3T+ cohort ($n = 33$). **a** Gross levels of different information dynamics in naturalistic human brain activity (bits) collapsed across individuals. The ordering of the modes by level replicates ($r = 0.964$, $p_{\text{perm}} = 2.00 \times 10^{-4}$), but $U_X \to U_X$ is third rather than first here, behind $R \to U$ and $U \to R$. **b,c** Group-average topographic patterns of global (mean across interactions for each region) information dynamics. **d** Group-average topographic patterns of global (mean across interactions for each region) 'computational diversity' ($D$) indexing the amount of distinct modes information dynamics engaged in an interaction (see Methods). $D$ is shown here with the atoms expressed in the original cohort's units, since the sampling interval sets the balance of magnitudes; the rescaling and its consequences are given in Supplementary Note 2 and Supplementary Fig. S16. **e** Gradient of ranked group-average $S \to S - R \to R$ maps, the redundancy-to-synergy gradient of Luppi *et al.*[1]. **f** Redundancy-to-synergy gradient values for each subject subset by visual processing areas[2], canonical resting-state networks (Yeo-Krienen-17)[3], and canonical cytoarchitectonic classes (Mesulam)[4] (points are participants). (*,**,*** indicates p<.05,.01,.001 FDR-corrected Benjamini-Hochberg procedure. VO: ventral occipital areas; LO: lateral occipital; V3A, V3B: dorsal extrastriate visual areas; hMT: human middle temporal area (V5); MST: medial superior temporal area; HC: parahippocampal; IPS/SPL: pooled intraparietal and superior parietal fields; FEF: frontal eye fields; Cont: control; DMN: default mode network; DAN: dorsal attention network; TPN: temporoparietal network; Lim: limbic; Sal: salience; VAN: ventral attention network; SMN: somatomotor; Vis: visual.)

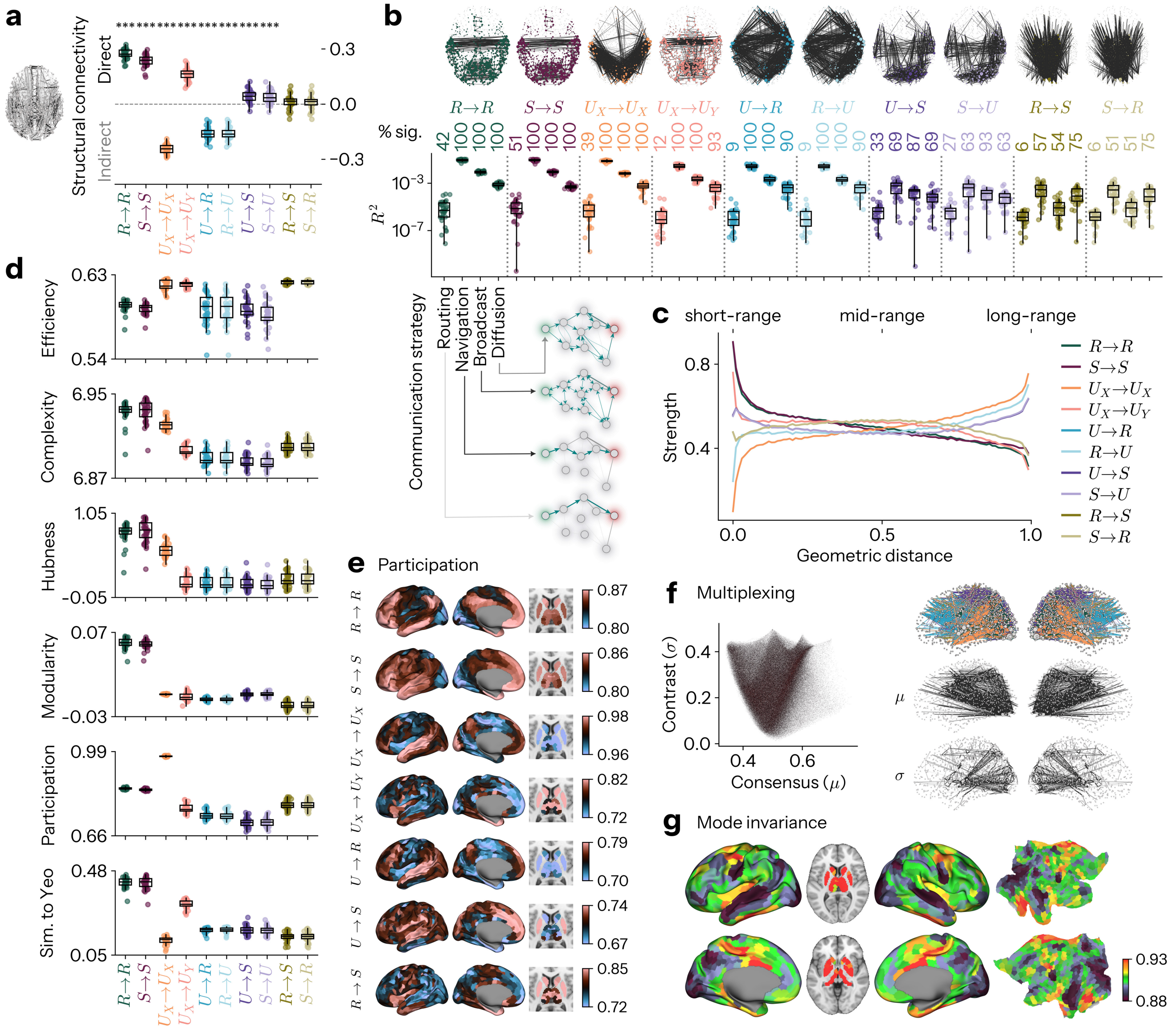


Extended Data Fig. 2. **Replication of the network properties of information dynamics.** As Fig. 2, in the NNDb-3T+ cohort ($n = 33$). **a** Relationship between information dynamics and types of structural connections. Rank-biserial correlation contrasts the atom's edge weights on directly connected region pairs against those on structurally unconnected (polysynaptic) pairs. Asterisks denote a one-sample $t$-test of the effect against zero. This is the panel that replicates least well: the sign of the effect agrees with Fig. 2a for only five of the ten modes, with $U_X \to U_Y$, $U \to R$ and $R \to U$ reversed and $R \to S$ and $S \to R$ no longer distinguishable from zero (Supplementary Note 2). **b** Top: the 1000 strongest edges of each atom's network. Prediction of networks of information dynamics from connectome communication models (see Methods). Box plots show the variance in each atom's edge weights explained ($R^2$) by a per-participant OLS regression on the corresponding communication matrix, points denoting participants. *% sig.* is the percentage of participants whose $R^2$ exceeds that of strength-preserving randomised surrogates of the communication matrix ($p_{\text{perm}} < 0.05$) **c** Relationship between information dynamics and scaled geometric (Euclidean) distance (mean ± SEM). **d** Global topological properties of networks of information dynamics, from top to bottom: global efficiency (Efficiency), entropy of eigenmode contributions (Complexity), hierarchical heavy-tailed connectivity via the $s$-metric of scale-freeness (Hubness), separability of communities $Q$ (Modularity), prevalence of mixed intermodular connectivity from the participation coefficient (Participation), and similarity to the Yeo-Krienen-17 networks via adjusted mutual information (Sim. to Yeo) (see Methods) (points are participants). **e** Topographic plots of nodal participation coefficient for modes of information dynamics. The $U \to R$ and $R \to U$ maps are the only two of the thirteen nodal topographies here that do not significantly replicate (Supplementary Fig. S17). **f** Conceiving each network of information dynamics as a layer in a multilayer graph, we summarise each edge's profile across layers by its mean $\mu$ (consensus) and standard deviation $\sigma$ (contrast). Left: joint scatter density plot. Right, top to bottom: the top 500 edges per atom, the top 1000 consensus edges ($\mu$), and the top 1000 contrast edges ($\sigma$). **g** Topographic plot of mode invariance: the global similarity of each region's connectivity profile across all modes of information dynamics (see Methods). (*** indicates p<.001 FDR-corrected Benjamini-Hochberg procedure.)

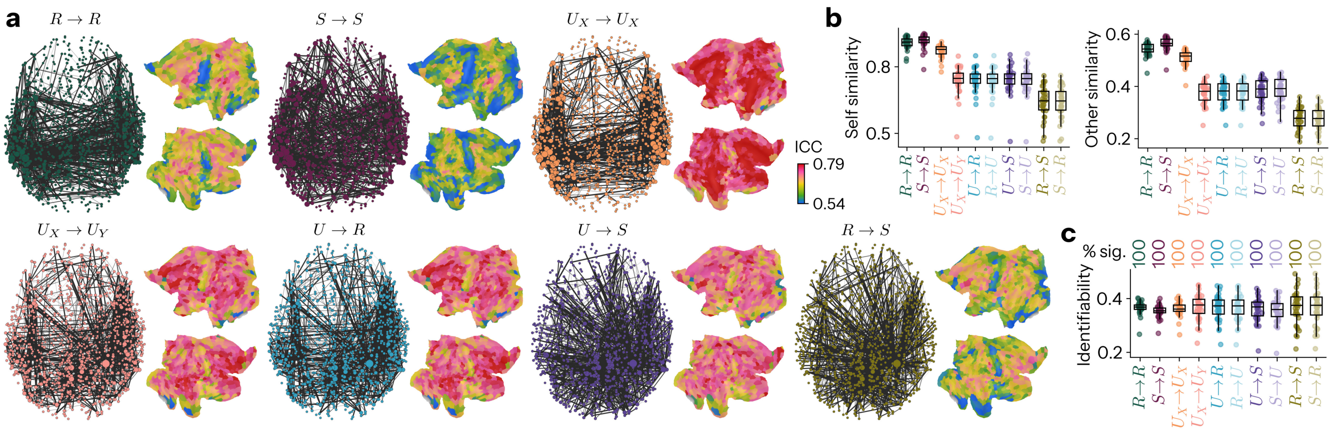


Extended Data Fig. 3. **Replication of individual differences in information dynamics.** As Fig. 3, in the NNDb-3T+ cohort ($n = 33$). **a** Split-half reliability of each connection, via intra-class correlation (ICC) of its weight between the two halves of each individual's scan. Brain networks show the 1000 most reliable edges; cortical flat maps show node-averaged ICC. **b** Fingerprinting of individuals from each mode of information dynamics. Left: correlation between a mode's connectivity in the first and second half of the same individual's scan (Self similarity). Right: correlation between one individual's scan half and the matching half of every other individual (Other similarity). **c** Self minus Other similarity (Identifiability), tested against a permutation null of shuffled subject labels (see Methods). *% sig.* is the percentage of participants exceeding that null. Every participant is identifiable from every mode, as in the original, but identifiability shows no difference across modes ($F(9, 288) = 1.346$, $p = 0.213$, Supplementary Note 2).

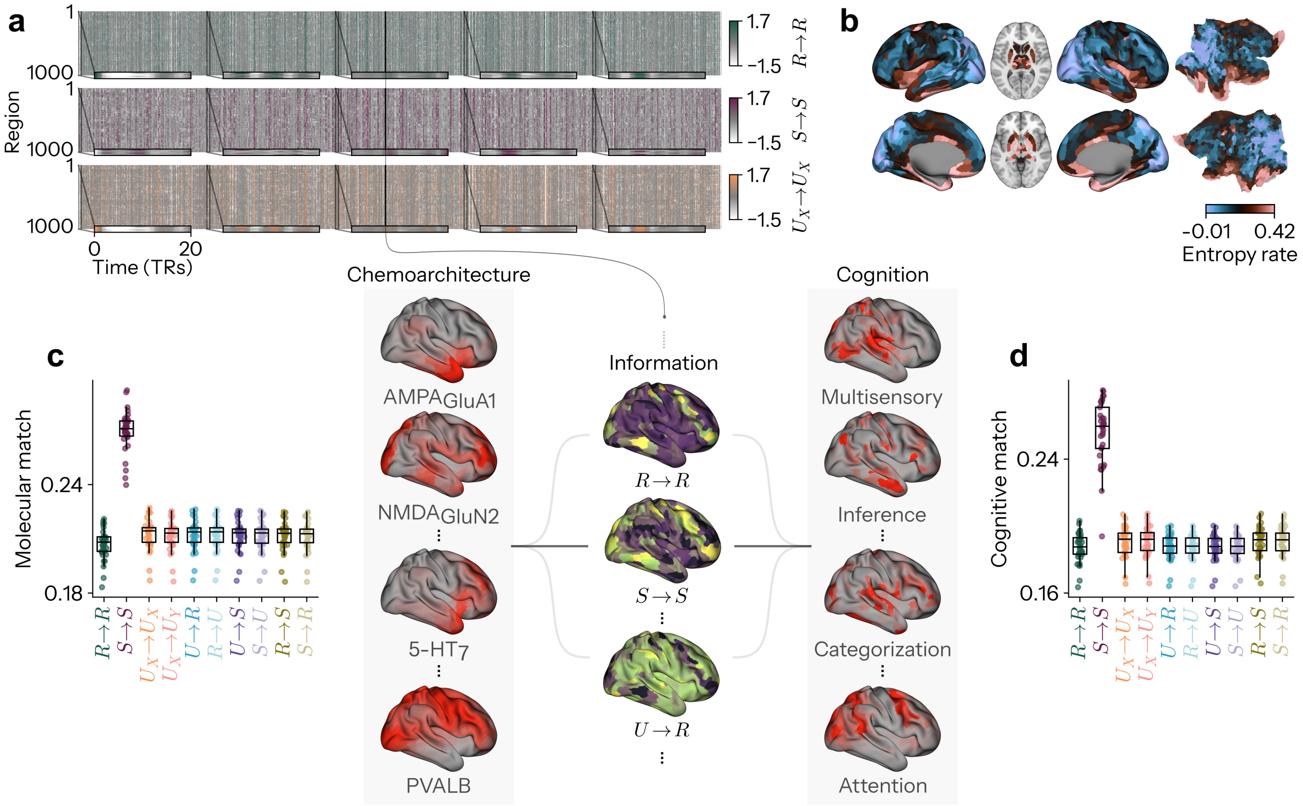


Extended Data Fig. 4. **Replication of reconfigurations of information dynamics in step with chemoarchitectonic and cognitive maps.** As Fig. 4, in the NNDb-3T+ cohort ($n = 33$). **a** Time-resolved fluctuations in information storage (Top: $R \to R$, Middle: $S \to S$, Bottom: $U_X \to U_X$), averaged over interactions for each region for example participant (Sub-04, 4,449 volumes, 6,674 total seconds). **b** Replication of topography of temporal prediction error, estimated via mean state-space entropy rate of regional information-dynamics time courses (nats). The entropy rate is a per-sample quantity, so its absolute level is set by the sampling interval and is not comparable between cohorts (group means −1.149 nats/sample at TR = 1 s against +0.173 nats/sample at TR = 1.5 s). Lower panel: At each timepoint, the whole-brain map of each mode of information dynamics is compared against reference maps of brain chemoarchitecture (left; AHBA mRNA expression maps for >1,900 brain-specific genes) and meta-analytic cognitive function (right; NeuroSynth meta-analytic term maps from over >14,000 neuroimaging experiments) and averaged across time, measuring the extent to which each mode of information dynamics mirror molecular or cognitive patterns respectively (see Methods). **c** Replication of molecular match: correspondence between each mode's time-resolved maps and the chemoarchitectural maps (points are participants). **d** Replication of cognitive match: correspondence between each mode's time-resolved maps and the cognitive maps (points are participants).

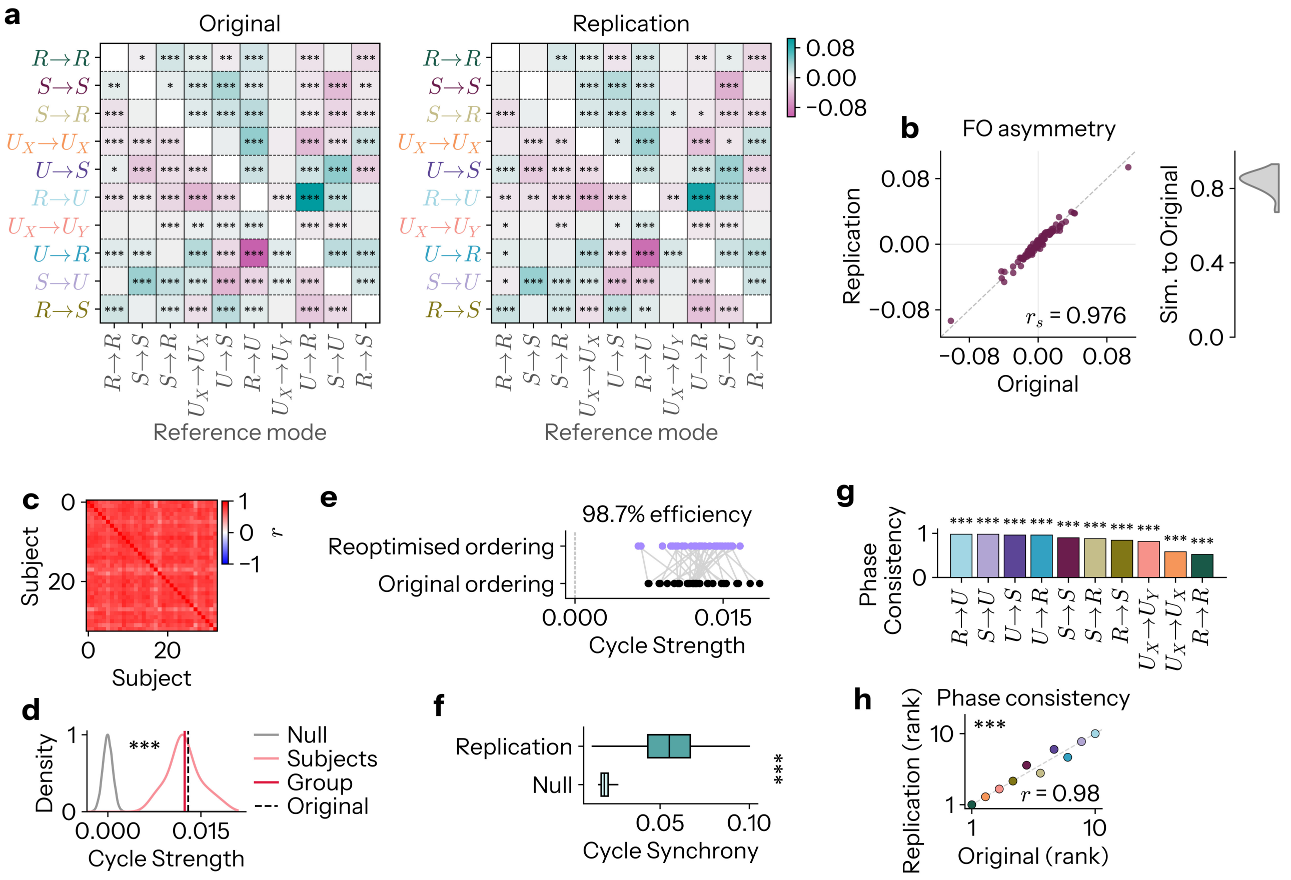


Extended Data Fig. 5. **Replication of the cycle of global information dynamics.** Throughout, *Original* is the NNDb cohort of Fig. 5 ($n = 86$) and *Replication* the NNDb-3T+ cohort ($n = 33$). **a** Group-level fractional occupancy (FO) asymmetry matrices of the two cohorts, as in Fig. 5c, each showing FDR-corrected one-sample $t$-test against zero across its own participants. Of the 90 directed edges, 74 are significant in the original and 70 here, with 65 significant in both ($p_{\mathrm{FDR}} < 0.05$) **b** *Left*: the 90 directed edges of **a**, one cohort against the other. *Right*: distribution over participants of the correlation between each participant's own asymmetry matrix and the original group matrix. **c** Correlation between the FO asymmetry matrices of every pair of participants in this cohort, as in Supplementary Fig. S12c. Every participant here watched the same film, so unlike in the original cohort there is no film contrast to draw. **d** Cycle strength $S$, the net rotational momentum of the FO asymmetry matrix around the ordering, measured on the *original* cohort's circle: at group level (crimson) and per participant (pink), against a null (grey) of 10,000 permutations that independently relabel the modes within each participant and recompute $S$ on the same circle. The original cohort's group value on this circle is drawn dashed. **e** Each participant's cycle strength on the original cohort's circle (black) joined to their strength on the circle optimised within this cohort (purple). *Efficiency* is the ratio of the group strength on the original circle to that on the re-optimised one, that is, how much of the rotational flow available in these data the original ordering already captures. **f** Cycle synchrony between pairs of participants, the time-averaged phase-locking value (PLV) between their cycle-phase time series, against a null of 1,000 independent circular shifts of the same phase series. **g** Phase consistency of each mode across participants in this cohort, the mean resultant length $R$ of the Rayleigh test applied to the 33 preferred phases of that mode, as in Supplementary Fig. S12b. Modes are ordered by $R$. **h** The ten values of **g** against the corresponding values of the original cohort, in ranks, coloured by mode. (*,**,*** indicates p<.05,.01,.001 FDR-corrected Benjamini-Hochberg procedure).

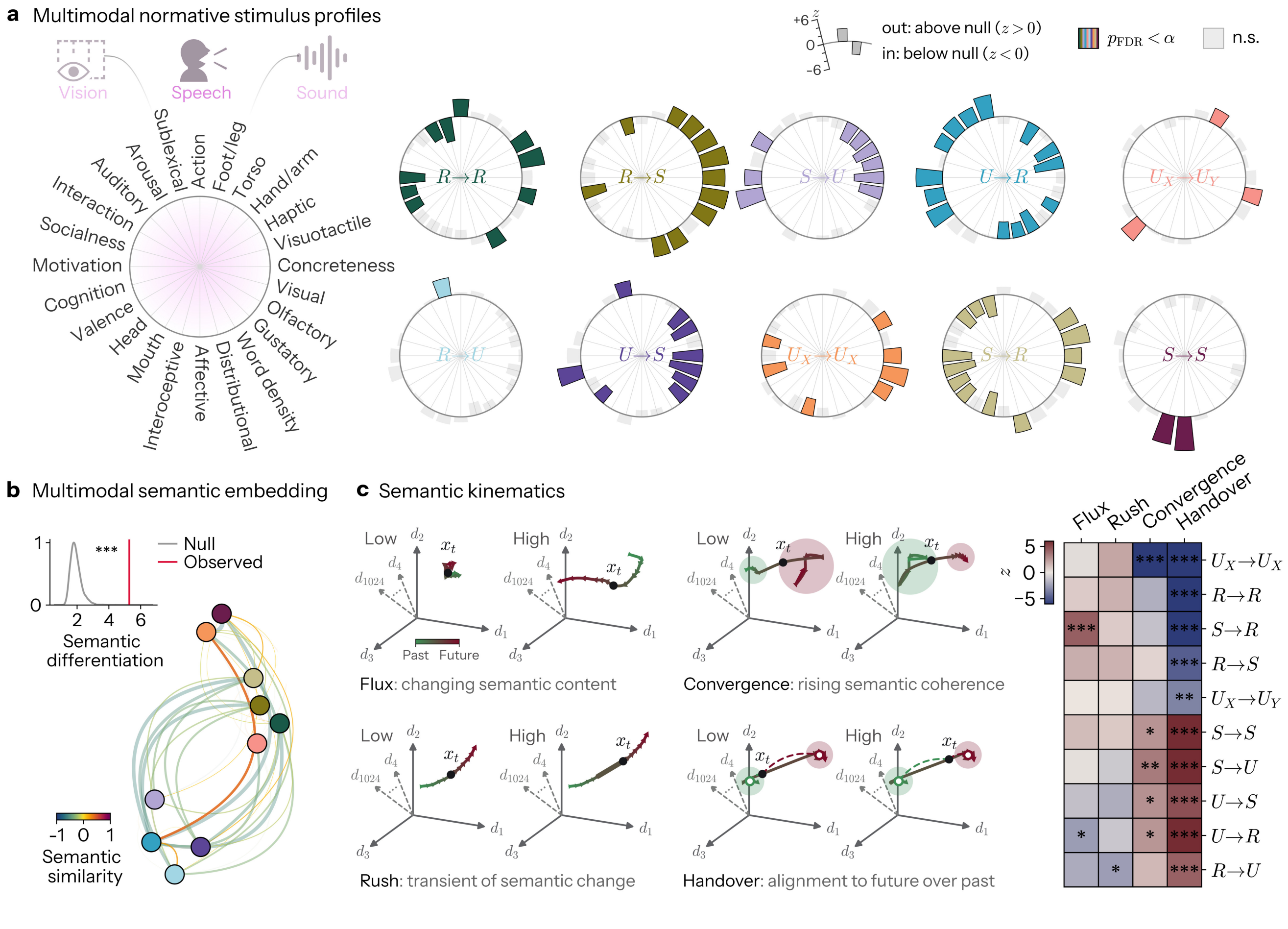


Extended Data Fig. 6. **Replication of the cognitive profile of global information dynamics.** As Fig. 6, in the NNDb-3T+ cohort ($n = 33$). Every participant in this cohort watched *Back to the Future*, one of the ten films annotated for the original analysis, so the annotation, the 24 lexical norms and the 45 trajectory measures are the same in both, and the film-residualisation of Fig. 6 reduces to centering. Mode windows span 1.5s here against 1s in the original and are resampled onto the annotation's 1s timeline of the film; the 5s haemodynamic lag is unchanged. **a** Normative profiles of each mode, as Fig. 6a. Spokes are held in the order seriated on the original cohort rather than re-seriated here, so that the dials can be read directly against those of Fig. 6a. **b** Semantic differentiation of the modes and their semantic similarity network, as Fig. 6b. The null here comprises 5,000 circular shifts. **c** Semantic kinematics, as Fig. 6c. Modes are ordered by their loading on flux *in the original cohort* rather than in this one, again so that the two panels may be compared row by row. (*,**,*** indicates p<.05,.01,.001 FDR-corrected Benjamini-Hochberg procedure).

#### S2. SUPPLEMENTARY NOTE 1: COMPLEXITY AND DIVERSITY

We note that 'computational diversity' ($D$) here is distinct from the measure of 'representational complexity' ($C$) introduced by Ehrlich & Schneider et al.[5]. Representational complexity quantifies the difficulty of accessing the information a system carries by weighting each atom by the number of variables that must be observed jointly to retrieve it. Formally, for a partial information decomposition (PID) of the mutual information $I(T:\mathbf{S})$ that $n$ sources $\mathbf{S} = \{S_1, \ldots, S_n\}$ carry about a target $T$,

$$C = \frac{1}{I(T:\mathbf{S})} \sum_{\alpha} \Pi(T:\mathbf{S}_\alpha)\, m(\alpha), \qquad m(\alpha) = \min_{\mathbf{a}\in\alpha} |\mathbf{a}|, \tag{S1}$$

where $\Pi(T:\mathbf{S}_\alpha)$ is the atom indexed by antichain $\alpha$, and its *degree of synergy* $m(\alpha)$ is the size of the smallest collection of sources from which that atom's information can be obtained. $C$ is therefore the mean degree of synergy, weighted by each atom's fractional information contribution, ranging from 1 when all information is available from single sources to $n$ when it is held purely synergistically across all of them.

Instead, we sought a measure of how *diverse* the modes of information dynamics engaged in an interaction are, without privileging any form of information over another. The contrast is structural: $C$ maps each atom to an external property, its degree of synergy, and averages over it, whereas $D$ is a functional of the distribution of information across the atoms itself. Invariant to atom identity, $D$ is maximal when information is spread evenly across all modes and minimal when it is concentrated in one.

Both measures are relative, and therefore invariant to the total mutual information. The difference in their construction and normalisation is nonetheless consequential for decompositions for which local positivity does not hold. In the case of $C$, each atom enters weighted by its signed fractional contribution $\Pi(T:\mathbf{S}_\alpha)/I(T:\mathbf{S})$. When the decomposition admits negative atoms these weights are no longer all non-negative, and $C$ ceases to be a convex combination of the degrees of synergy: it loses both its guarantee of lying in $[1, n]$ and its interpretation as a mean number of jointly observed variables, and it diverges as cancellation drives the total information in its denominator toward zero. In the case of $D$, normalisation is instead by the sum of squared atom values $\sum_k a_k^2$, which is strictly positive unless every atom vanishes. Squaring discards the sign of each atom, so the $p_k$ remain a valid probability distribution and $D$ a well-defined quantity on $[1/n, 1]$, whatever the signs of the underlying atoms and however much they cancel.

Importantly, $C$ is defined within PID, which designates a single target $T$ that fixes the degree of synergy $m$, and hence $C$. In $\Phi$ID, by contrast, there is no single target, and one would need to choose whether $m$ is taken with respect to the past, the future, or both. We note, however, that $D$ applies just as readily to PID as to $\Phi$ID.

#### S3. SUPPLEMENTARY NOTE 2: REPLICATION

##### Design and scope

We collected an additional multimodal naturalistic neuroimaging dataset[6], and repeated every analysis of Figs. 1–6 end to end in this new, independent cohort (NNDb-3T+, $n = 33$; see Methods). The two datasets differ on many axes to ensure cross-validity of the information architecture: field strength (1.5T against 3T), repetition time (1 s against 1.5 s), spatial resolution (3.2 mm against 2 mm isotropic), preprocessing pipeline, cohort size (86 against 33), and stimulus design. The results are shown in Extended Data Figs. 1–6, and their agreement with the original quantified in Supplementary Figs. S16–S25.

##### How agreement was assessed

Each replicated quantity is compared against its original by Spearman correlation, tested against a null tailored to the object being compared, with $p$ two-tailed and computed as the fraction of nulls whose $|r|$ meets or exceeds the observed $|r|$, with the customary $+1$ in numerator and denominator. Three families of null were used.

Whole-brain maps (1054 regions) are tested against 1,000 surrogates of the original map generated by BrainSMASH variogram matching[7], which preserves the empirical distance-dependence of the map and resamples its values, so that the null is a map as spatially autocorrelated as this one rather than an arbitrary reshuffling. Matrices are tested against node-relabelling permutations, preserving their degree sequence and edge-weight distribution. The FO asymmetry matrix is read as directed, keeping $A_{ij}$ and $A_{ji}$ as separate observations, since its entire content is its antisymmetry. Profiles over the ten modes are tested against 10,000 mode-relabelling permutations. A permutation $p$ cannot fall below $1/(n+1)$, such a $p$ is therefore reported as an upper bound, $p_{\text{perm}} < 10^{-3}$ for 1,000 nulls and $p_{\text{perm}} < 10^{-4}$ for 10,000, and is noted as the permutation floor where it is reported; a permutation $p$ given with an equality lies strictly above the floor.

##### Topographic organisation

The regional topographies of Extended Data Fig. 1b,c reproduce those of Fig. 1 closely: 15 of 16 regional maps replicate at $p < 0.05$, with $r$ ranging from $-0.022$ to $+0.894$ (median $+0.846$; Supplementary Fig. S16). Per mode, $R\to R$ $r = 0.776$, $S\to S$ $r = 0.846$, $U_X \to U_X$ $r = 0.894$, $U_X \to U_Y$ $r = 0.834$, $U\to R$ $r = 0.833$, $R\to U$ $r = 0.833$, $U\to S$ $r = 0.765$, $S\to U$ $r = 0.768$, $R\to S$ $r = 0.848$ and $S\to R$ $r = 0.846$, all at $p_{\text{perm}} < 10^{-3}$ (permutation floor, 1,000 SA preserving surrogates). The full connectomes behave likewise, with all ten modes replicating (median $r = +0.741$): $R\to R$ $r = 0.899$, $S\to S$ $r = 0.913$, $U_X\to U_X$ $r = 0.908$, $U_X\to U_Y$, $U\to R$ and $R\to U$ $r = 0.600$ to 0.601, $U\to S$ and $S\to U$ $r = 0.745$, and $R\to S$ and $S\to R$ $r = 0.737$, all at $p_{\text{perm}} < 10^{-3}$ (permutation floor, 1,000 node-relabelling permutations). Computational diversity $D$ does not, for reasons taken up below. The ordering of the modes by gross level (Fig. 1a) is recovered at $r = 0.964$, $p_{\text{perm}} = 2.00 \times 10^{-4}$, and the levels themselves separate as strongly as in the original (rm-ANOVA $p = 2.33 \times 10^{-57}$, $\eta^2_G = 0.998$; pairwise comparisons $p_{\text{FDR}} = 7.35 \times 10^{-55}$ to $1.52 \times 10^{-5}$).

The redundancy-to-synergy gradient replicates at $r = 0.854$, $p_{\text{perm}} < 10^{-3}$ (permutation floor), and continues to be expressed by each participant individually ($r = 0.626 \pm 0.064$ against the group gradient here, against $0.539 \pm 0.083$ in the original). Its profile across the reference schemes of Fig. 1f is likewise recovered, across visual areas ($r = 0.976$, $p_{\text{perm}} < 10^{-4}$, at the permutation floor of 10,000 nulls, sign agreement 12/16; the four disagreements are V1, V2, VO1 and VO2, none of which reaches significance here), Yeo-Krienen networks ($r = 0.806$, $p_{\text{perm}} = 3.00 \times 10^{-4}$, 13/18) and Mesulam classes ($r = 1.000$, $p_{\text{perm}} = 0.083$, 4/4; with only four classes the permutation distribution has 24 members, two of which attain $|r| = 1$, so a two-tailed $p$ cannot fall below $2/24 = 0.083$ however perfect the agreement, which is exactly the value obtained). The same regions carry the synergistic end of the gradient: hMT ($T = 11.401$, $p_{\text{FDR}} = 6.68 \times 10^{-12}$), MST ($T = 11.901$, $p_{\text{FDR}} = 4.37 \times 10^{-12}$), the temporoparietal network ($T = 13.709$, $p_{\text{FDR}} = 5.47 \times 10^{-14}$) and heteromodal cortex ($T = 14.402$, $p_{\text{FDR}} = 6.21 \times 10^{-15}$), against subcortex ($T = -19.783$, $p_{\text{FDR}} = 2.96 \times 10^{-18}$) and paralimbic cortex ($T = -11.183$, $p_{\text{FDR}} = 1.83 \times 10^{-12}$) at the redundant end. The stream-level structure of the gradient recurs in full: the ventral areas hV4 and PHC2 remain redundancy-dominant ($p_{\text{FDR}} = 6.11 \times 10^{-4}$ and $2.78 \times 10^{-3}$), the pooled intraparietal and superior parietal fields again fall short of hMT and MST ($p = 9.95 \times 10^{-8}$) while carrying no consistent internal ordering, and the frontal eye fields again revert to redundancy dominance ($-126.4$ rank units, $p_{\text{FDR}} = 8.83 \times 10^{-6}$), more strongly here than in the original ($-58.4$).

##### Normalisation of computational diversity

Computational diversity is the one quantity in Fig. 1 that does not transfer as computed. Taken on the raw atoms, its topography is uncorrelated with the original across regions ($r = -0.022$, $p_{\text{perm}} = 0.811$) and weakly anticorrelated across edges ($r = -0.200$, $p_{\text{perm}} < 10^{-3}$, at the permutation floor), and its dynamic range collapses: the group regional map spans $[0.709, 0.732]$ here against $[0.580, 0.656]$ in the original, a range narrower by a factor of three and displaced upwards.

The cause is not the topography but the units, since $D$ is a functional of the distribution of magnitudes, which moves whenever the sampling interval rescales them differentially. Taking the per-atom root-mean-square over every subject and every edge in each cohort, the ratio of this run's atom magnitudes to the original's is close to uniform for thirteen of the sixteen atoms (0.883 to 0.890), but is 0.826 for $S \to S$ and only 0.564 and 0.561 for the two atoms that compose unique storage $U_X \to U_X$. The longer TR therefore flattens the two unique-storage atoms relative to the rest by roughly a further 36%. We therefore express both cohorts' atoms in common units before computing $D$, dividing each of this run's atoms by its own RMS ratio so that the sixteen atoms carry the same relative scale in both. The correction is a single vector, applied identically at every edge and every region: it carries no spatial information whatever, and so cannot manufacture topographic agreement, and the original run is untouched, its ratio being one by construction. In common units $D$ recovers both its range (regional $[0.548, 0.657]$ against the original's $[0.580, 0.656]$) and its topography, at $r = 0.859$, $p_{\text{perm}} < 10^{-3}$ across regions and $r = 0.880$, $p_{\text{perm}} < 10^{-3}$ across edges (both at the permutation floor) (Supplementary Fig. S16). Extended Data Fig. 1d shows the common-units map.

##### Network organisation

Of the 13 nodal maps in Extended Data Fig. 2, 11 replicate at $p < 0.05$, with a median $r$ of 0.661 (Supplementary Fig. S17): nodal participation for $U_X \to U_X$ at $r = 0.939$, $R \to S$ at 0.798, $S \to R$ at 0.793, $R \to R$ at 0.691, $S \to S$ at 0.657, $U_X \to U_Y$ at 0.648, $S \to U$ at 0.297 and $U \to S$ at 0.289, together with mode invariance ($r = 0.887$), the nodal consensus $\mu$ ($r = 0.758$) and the nodal contrast $\sigma$ ($r = 0.661$), all at $p_{\text{perm}} < 10^{-3}$ (at the permutation floor of 1,000 nulls). At the edge level the multiplex summaries hold over all 554,931 edges (consensus $r = 0.809$, contrast $r = 0.609$).

The topological ordering of the modes (Fig. 2d) is reproduced for five of the six measures (Supplementary Fig. S18): Efficiency $r = 0.891$ ($p_{\text{perm}} = 0.001$), Complexity $r = 0.830$ (0.005), Hubness $r = 0.770$ (0.014), Modularity $r = 0.733$ (0.022) and similarity to Yeo-Krienen $r = 0.794$ (0.009). The pairings of Fig. 2d are preserved: $R \to R$ and $S \to S$ are again the two most complex, hierarchical, modular and Yeo-Krienen-like networks, $R \to S$ and $S \to R$ again the most efficient and least modular pair, and $U_X \to U_X$ again the most participatory mode of all. Each remains significantly separated from the mean of the other nine (for $S \to S$, $p_{\text{FDR}} = 9.78 \times 10^{-18}$, $1.74 \times 10^{-19}$, $2.49 \times 10^{-33}$ and $1.78 \times 10^{-28}$ on complexity, hubness, modularity and Yeo-Krienen similarity; for $S \to R$, $p_{\text{FDR}} = 8.25 \times 10^{-16}$ and $3.16 \times 10^{-25}$ on efficiency and modularity; for $U_X \to U_X$, $p_{\text{FDR}} = 4.82 \times 10^{-39}$ on participation), and all 60 mode-by-measure contrasts survive $p_{\text{FDR}} < 0.05$, against 59 of 60 in the original. Rank order *within* those pairs is less stable, however, and $R \to R$ takes first place from $S \to S$ on complexity, modularity and Yeo similarity, on differences of the fourth decimal place. The communication-model results carry over both in ordering (mean $\rho = 0.960$ across modes, $p_{\text{perm}} < 10^{-4}$, at the permutation floor of 10,000 nulls, against a null of $-0.001 \pm 0.183$) and in magnitude, with mean $\bar{R}^2$ of $8.74 \times 10^{-6}$ for Routing, 0.034 for Navigation, 0.003 for Broadcast and $3.96 \times 10^{-4}$ for Diffusion, against $5.44 \times 10^{-6}$, 0.035, 0.003 and $2.50 \times 10^{-4}$ originally. The pooled ordering Navigation > Broadcast > Diffusion > Routing is identical in the two cohorts, though it now holds within 8 of the 10 modes individually rather than all 10, with $R \to S$ and $S \to R$ better predicted by diffusion than by broadcast. The strength-by-distance curves of Fig. 2c replicate for all ten modes, but unevenly: near-perfectly for the storage modes ($R \to R$ $r = 0.997$, $S \to S$ $r = 0.997$, $U_X \to U_X$ $r = 0.998$), strongly for the unification and interconversion modes ($r = 0.584$ to 0.681), and only just for $U_X \to U_Y$, $U \to R$ and $R \to U$ ($r = 0.223$, 0.213 and 0.213; $p = 0.028$, 0.036 and 0.036).

##### Individual differences

The reliability maps of Extended Data Fig. 3 replicate for every mode, both nodally ($r = 0.267$ to 0.549, all $p_{\text{perm}} < 10^{-3}$, at the permutation floor; Supplementary Fig. S19) and edge-wise ($r = 0.183$ to 0.468, all $p_{\text{perm}} < 10^{-3}$, likewise at the floor), over a comparable range of values (ICC from $-0.4321$ to 0.9919 here against $-0.1087$ to 0.9906 originally, differing across modes at $F = 1.72 \times 10^5$, $p < 10^{-300}$, against $F = 3.19 \times 10^5$). Fingerprinting is unchanged in kind and in degree: 100% of participants exceed the identity-shuffling null for all ten modes here, against 100% for eight modes and 99% for $R \to S$ and $S \to R$ in the original, and the profiles of self and other similarity across modes are recovered almost exactly (self $r = 0.939$, $p_{\text{perm}} = 4.00 \times 10^{-4}$; other $r = 0.976$, $p_{\text{perm}} = 2.00 \times 10^{-4}$;

Supplementary Fig. S20), with the same modes at the extremes (self $F(9, 288) = 378.139$, $p = 7.40 \times 10^{-154}$; other $F(9, 288) = 1.18 \times 10^3$, $p = 1.41 \times 10^{-221}$). Individual networks remain typical of the group in the same order as before ($S \to S$ $0.774 \pm 0.031$, $R \to R$ $0.762 \pm 0.030$, $U_X \to U_X$ $0.749 \pm 0.040$, against $R \to S$ $0.597 \pm 0.072$ and $S \to R$ $0.596 \pm 0.073$, although the gradient, the most variable quantity of all in the original, is no longer so here at $0.635 \pm 0.060$; compare Supplementary Fig. S6), and partitioning each participant's networks individually reproduces the group partition to the same degree (AMI $0.396 \pm 0.088$, range 0.139 to 0.551, against $0.385 \pm 0.088$, range 0.138 to 0.574; 27 of 30 mode-by-measure contrasts surviving $p_{\mathrm{FDR}} < 0.05$, against 29 of 30; compare Supplementary Fig. S5).

#### Matching dynamics

The matching results of Extended Data Fig. 4 are faithfully reproduced. All five profiles replicate, and in every one the strongest mode is the same as in the original (Supplementary Fig. S21): brain genes $r = 0.964$ ($p_{\mathrm{perm}} < 10^{-4}$, at the permutation floor of 10,000 nulls), synapse genes $r = 0.964$ ($p_{\mathrm{perm}} < 10^{-4}$, likewise at the floor), neuron genes $r = 0.855$ ($p_{\mathrm{perm}} = 0.004$), cognitive terms $r = 0.842$ ($p_{\mathrm{perm}} = 0.004$) and the functional-to-structural ratio $r = 0.939$ ($p_{\mathrm{perm}} = 5.00 \times 10^{-4}$). $S \to S$ again stands apart from all nine other modes in every corpus (brain genes $F(9, 288) = 1.28 \times 10^3$, $p = 1.16 \times 10^{-226}$, weakest $S \to S$ contrast $t = 33.552$, $p_{\mathrm{FDR}} = 4.63 \times 10^{-26}$, $d = 5.620$; neuron genes $F(9, 288) = 1.21 \times 10^3$, $p = 3.36 \times 10^{-223}$; synaptome $F(9, 288) = 1.29 \times 10^3$, $p = 8.02 \times 10^{-227}$; cognitive $F(9, 288) = 470.623$, $p = 1.50 \times 10^{-166}$, weakest $S \to S$ contrast $t = 21.167$, $p_{\mathrm{FDR}} = 8.24 \times 10^{-20}$, $d = 4.308$), and the functional-to-structural ratio again separates the modes ($F(9, 288) = 64.457$, $p = 7.98 \times 10^{-64}$, 40 of 45 pairwise contrasts at $p_{\mathrm{FDR}} < 0.05$, against 41 of 45 originally). We also note that the topography of mean temporal prediction error (Extended Data Fig. 4b) replicates for every mode, at $r = 0.909$. The time-resolved reconfigurations of the gradient behave the same way: instantaneous gradients correlate weakly with the participant's own static gradient ($r = 0.060 \pm 0.086$ here, $0.054 \pm 0.096$ originally) and with the static group map ($0.046 \pm 0.079$ against $0.043 \pm 0.089$).

#### Global state dynamics and the cycle

The winner-take-all dynamics statistics of Fig. 5b are recovered essentially exactly (Supplementary Fig. S22): appearance rate $r = 1.000$, dwell time $r = 0.988$ and fractional occupancy $r = 0.988$, all $p_{\mathrm{perm}} < 10^{-4}$ (permutation floor, 10,000 mode relabellings), with $U_X \to U_X$ again the longest-dwelling and most occupied state and $U_X \to U_Y$ again the rarest and most fleeting. The modes differ on all three measures as strongly as before (appearance rate $F = 592.269$, $p = 4.37 \times 10^{-180}$, $\eta_G^2 = 0.941$; dwell time $F = 289.428$, $p = 1.19 \times 10^{-138}$, $\eta_G^2 = 0.865$; occupancy $F = 534.211$, $p = 5.46 \times 10^{-174}$, $\eta_G^2 = 0.943$).

The cycle replicates on every measure that does not depend on the sampling interval (Extended Data Fig. 5). The fractional occupancy asymmetry matrix, which is the object the cycle summarises, agrees at $r = 0.976$ over its 90 directed edges ($p_{\mathrm{perm}} < 10^{-4}$, at the permutation floor of 10,000 node-relabelling permutations; Pearson $r = 0.986$), with 70 of 90 edges significant here against 74 of 90 originally, 65 significant in both, and sign concordance of 71/74 (96%, $p = 7.16 \times 10^{-18}$) on the edges the original called significant. The agreement is not carried by a subset of participants: every one of the 33 correlates positively with the original group matrix ($r = 0.837 \pm 0.054$, minimum 0.673, $T = 87.391$, $p = 1.18 \times 10^{-39}$), and the asymmetry structure is as conserved within this cohort as within the original (mean pairwise $r = 0.714 \pm 0.079$, 528/528 pairs positive, range 0.398 to 0.867, against $0.714 \pm 0.090$ originally).

Measured on the discovered circle, the rotational flow is clearly present ($S = 0.0124$ against a null of $-0.0000 \pm 0.0008$, $p_{\mathrm{perm}} < 10^{-4}$ at the permutation floor of 10,000 nulls), holds in every participant ($S = 0.0124 \pm 0.0027$, 33/33 positive, $T = 26.384$, $p = 2.77 \times 10^{-23}$), and is strong enough that the weakest participant ($S = 0.0074$) exceeds the strongest of the 10,000 null permutations ($S = 0.0027$). Its magnitude matches the original's on that same circle ($S = 0.0130$, 86/86 positive). Re-optimising the ordering for these 33 participants yields a circle of only marginally greater strength ($S = 0.0125$), with the original ordering already capturing 98.7% of the rotational flow. Phase consistency is likewise recovered: all ten modes are Rayleigh-significant here as in the original, at a comparable concentration (mean $R = 0.841$ against 0.850; weakest mode $R = 0.519$, $p = 7.99 \times 10^{-5}$), and the profile of $R$ across modes agrees at $r = 0.976$, $p_{\mathrm{perm}} = 2.00 \times 10^{-4}$. Cycle synchrony across participants decisively exceeded a circular-shift null (observed PLV $0.055 \pm 0.008$ against $0.020 \pm 0.002$; paired $T(32) = 21.997$, $p = 6.93 \times 10^{-21}$, $d = 5.694$, $BF_{10} = 5.60 \times 10^{17}$, 33/33 participants, $p_{\mathrm{perm}} < 10^{-3}$, at the permutation floor).

#### Stimulus semantics and kinematics

Given the use of 'Back to the Future' again as the stimulus, which was annotated on a virtual timeline of 1s core unit and a 3s context window, we absorb the change of TR by mapping each brain sample onto the annotation row on which it opens, $\lfloor 1.5i \rfloor$. A 1.5s window thus always straddles two of the 1s annotation bins, and incorporating the second of them changes nothing about the replication ($r = 0.981$ between the two mode by norm tables; Supplementary Figs. S23 and S24).

The semantic geometry replicates. The omnibus partition of the embedding by mode identity again exceeds its circular-shift null (pseudo-$F = 5.283$, $p_{\text{perm}} = 4.00 \times 10^{-4}$, one of the 5,000 nulls reaching the observed value), and 29/45 pairwise separations survive correction. The pairwise structure itself, the object the similarity network of Fig. 6b summarises, agrees at $r = 0.738$, $p_{\text{perm}} < 10^{-4}$, at the permutation floor of 10,000 nulls. In fact, the fraction of semantic variance attributed to mode identity is higher here than in the original ($R^2 = 3.24 \times 10^{-4}$ against $1.23 \times 10^{-4}$).

The kinematic axes transfer as a set at $r = 0.737$ ($p_{\text{perm}} = 9.00 \times 10^{-4}$) and individually for three of the four: handover $r = 0.867$, convergence $r = 0.842$ and rush $r = 0.685$ ($p_{\text{FDR}} = 0.006$, 0.008 and 0.0499; Supplementary Fig. S25), with flux falling just short ($r = 0.636$, $p_{\text{FDR}} = 0.057$). All four again vary reliably across modes, and their ordering by strength is preserved, with handover again by far the strongest ($SS = 362.3$ here against 243.0, $p_{\text{perm}} = 2.00 \times 10^{-4}$) and rush again the weakest (16.4 against 95.2). Flux and rush are the axes materially weakened, flux falling from $SS = 138.0$ to 42.5 and rush from 95.2 to 16.4, both off the permutation floor ($p_{\text{perm}} = 6.40 \times 10^{-3}$ and 0.018). Of the 40 cells, 18 survive correction here against 29 originally.

The three checks that the axes track their constructs rather than the measures chosen behave as they did in the original. Clustering of the 45 measures recovers the four assigned families for 42 of them (`coherence_asym_3` and `coherence_change_7` clustering with handover, and `lag_asym_6` with convergence as in the original); the axes are again close to independent (largest absolute correlation $r = 0.376$ against 0.398, two correlation-matrix eigenvalues above one in both); and the measures again reproduce the profile of their axis when tested on their own against the leave-one-out composite of their family, though less uniformly than before (mean $r = +0.71$ for flux, +0.98 for handover, +0.61 for convergence and +0.61 for rush, against +0.71, +0.98, +0.82 and +0.85). Two of the 45 invert against their family here, `coherence_asym_24` and `speed_vs_next3` at $r = -0.271$ and $-0.181$, as do two in the original, `lag12_distance` and `speed_sd_past12` at $-0.538$ and $-0.263$. The relative typicality of measures within their families is itself preserved ($r = 0.986$ over the 45 measures).

#### What does not replicate

*Direct against indirect structural connectome support.* The rank-biserial profile of Fig. 2a correlates across modes at $r = 0.733$ ($p_{\text{perm}} = 0.023$), but its *signs* agree for only five of the ten modes. The two storage modes keep their preference for monosynaptic connections ($R \to R$ $T = 57.907$, $p_{\text{FDR}} = 5.71 \times 10^{-33}$; $S \to S$ $T = 41.996$, $p_{\text{FDR}} = 4.87 \times 10^{-29}$), $U_X \to U_X$ keeps its preference for polysynaptic ones ($T = -56.050$, $p_{\text{FDR}} = 8.03 \times 10^{-33}$), and $U \to S$ and $S \to U$ remain positive though much attenuated ($T = 7.344$ and 6.278, against 46.730 and 45.489 originally). But, $U_X \to U_Y$ reverses from negative to positive ($T = 27.588$, $p_{\text{FDR}} = 1.74 \times 10^{-23}$), $U \to R$ and $R \to U$ reverse from positive to negative ($T = -27.310$ and $-27.236$), and $R \to S$ and $S \to R$, strongly negative in the original, are not distinguishable from zero here ($T = 1.682$, $p_{\text{FDR}} = 0.114$, $BF_{10} = 0.660$; $T = 1.416$, $p_{\text{FDR}} = 0.167$, $BF_{10} = 0.462$).

*Nodal participation for copy and erasure.* Eleven of the thirteen nodal maps of Fig. 2 replicate; the two that do not are the participation coefficient maps of $U \to R$ ($r = -0.121$, $p_{\text{perm}} = 0.394$) and $R \to U$ ($r = -0.126$, $p_{\text{perm}} = 0.373$). Consistently, the ordering of modes by mean participation is the one topological profile of the six not to replicate ($r = 0.394$, $p_{\text{perm}} = 0.268$), although $U_X \to U_X$ remains the most participatory mode in both cohorts by a wide margin.

*The ordering of modes by edge reliability and identifiability.* Nodal ICC replicates map by map, but the *ranking* of modes by mean ICC does not ($r = 0.212$, $p_{\text{perm}} = 0.547$), nor does the ranking by identifiability ($r = 0.539$, $p_{\text{perm}} = 0.114$). This is less a failure than an absence: identifiability does not differ across modes at all in this cohort ($F(9, 288) = 1.346$, $p = 0.213$), and only 8 of the 45 pairwise contrasts survive correction, against 33 of 45 originally, so there is little ordering left to reproduce. Identifiability itself is unambiguous, at 100% of participants above the identity-shuffling null for all ten modes in both cohorts.

*Replicating profiles with shifting rank orders.* Several profile correlations are high while the identity of the rank-1 mode moves. Three results are notable in this respect. The ordering of modes by gross level (Fig. 1a) replicates at $r = 0.964$, but $U_X \to U_X$ falls from first to third, behind $R \to U$ and $U \to R$, so this cohort is not dominated by unique storage in the way the original is. The networks that are least complex and least efficient are $S \to U$ here rather than $U \to R$, although $U \to R$ is still significantly below the mean of the other nine on both ($p_{\text{FDR}} = 7.38 \times 10^{-19}$ and $1.19 \times 10^{-6}$). And within the storage pair, $R \to R$ takes first place over $S \to S$ on complexity, modularity and Yeo-Krienen similarity.

*Normative stimulus profiles of information dynamics not involving redundancy.* The normative results are the least uniform. In absolute terms, the normative results are, if anything, stronger: 79 of the 240 mode-by-norm cells survive correction here against 62 originally, over all ten modes in both, with larger effects (largest 0.071 SD and median 0.038 SD, against 0.046 and 0.023). Crucially, however, the correspondence with the original is nonetheless only partial. Taken as a table the two agree ($r = 0.449$, $p_{\mathrm{perm}} = 0.010$; Supplementary Fig. S23), but resolved mode by mode five of the ten profiles are recovered and five are not: $U \to R$ $r = 0.741$, $R \to R$ $r = 0.727$, $R \to U$ $r = 0.705$, $R \to S$ $r = 0.647$ and $S \to R$ $r = 0.616$, all at $p_{\mathrm{FDR}} \leq 0.004$, against $U_X \to U_Y$ $r = 0.263$, $S \to U$ $r = 0.208$, $U \to S$ $r = 0.202$, $U_X \to U_X$ $r = -0.018$ and $S \to S$ $r = -0.204$, none of which survive correction. Interestingly, these two sets divide exactly along the involvement of redundancy: the five recovered profiles belong to precisely those modes in which redundancy is either the source or the target of the transfer.

#### S4. SUPPLEMENTARY METHODS

##### Integrated information decomposition via information geometry

The decomposition introduced in the main text (see Methods) is realised as a system of sixteen linear equations for the sixteen ΦID atoms $I_\partial^{\alpha\to\beta}$[8]. Nine constraints are provided by mutual information terms between subsets of the past and future, while the remaining seven are supplied by geometric measures of synergy[9]. Together, these quantities uniquely determine the complete ΦID of a bivariate process. In this section, we describe how each quantity is obtained from the covariance structure of the lagged system and how the resulting equations are solved to obtain the ten symmetric information modes used in the main text.

For each pair of brain regions, we consider the bivariate process

$$\mathbf{Z} = (Z^1, Z^2), \tag{S2}$$

with past and future states

$$\mathbf{Z}_t = (Z_t^1, Z_t^2), \qquad \mathbf{Z}_{t'} = (Z_{t'}^1, Z_{t'}^2), \tag{S3}$$

where $t' = t + \tau$. The joint distribution

$$P = P(\mathbf{Z}_t, \mathbf{Z}_{t'}) \tag{S4}$$

is estimated from the observed time series under the Gaussian assumption. Consequently, the complete statistical structure of the four-dimensional system is specified by its covariance matrix, from which all mutual information terms and the probability distributions required for the geometric construction can be obtained.

###### *Mutual information constraints*

The first nine constraints are given by the mutual information between all relevant groupings of the two components of the past and future. Four correspond to individual source-target pairs

$$\begin{array}{ll} I(Z_t^1; Z_{t'}^1), & I(Z_t^1; Z_{t'}^2), \\ I(Z_t^2; Z_{t'}^1), & I(Z_t^2; Z_{t'}^2), \end{array} \tag{S5}$$

two describe the information between the full past and each future component

$$I(\mathbf{Z}_t; Z_{t'}^1), \qquad I(\mathbf{Z}_t; Z_{t'}^2), \tag{S6}$$

and two describe the converse quantities

$$I(Z_t^1; \mathbf{Z}_{t'}), \qquad I(Z_t^2; \mathbf{Z}_{t'}). \tag{S7}$$

The final constraint is the time-delayed mutual information (TDMI) between the complete past and future

$$I(\mathbf{Z}_t; \mathbf{Z}_{t'}). \tag{S8}$$

For jointly Gaussian variables, these quantities are obtained directly from the relevant covariance matrices. In particular, for two jointly Gaussian variable sets $\mathbf{A}$ and $\mathbf{B}$,

$$I(\mathbf{A}; \mathbf{B}) = \frac{1}{2} \ln \frac{\det \Sigma_{\mathbf{A}} \det \Sigma_{\mathbf{B}}}{\det \Sigma_{\mathbf{A},\mathbf{B}}}, \tag{S9}$$

where $\Sigma_{\mathbf{A}}$, $\Sigma_{\mathbf{B}}$, and $\Sigma_{\mathbf{A},\mathbf{B}}$ denote the covariance matrices of the corresponding marginal and joint distributions. Thus, once the covariance matrix of $(\mathbf{Z}_t, \mathbf{Z}_{t'})$ has been estimated, all nine mutual information constraints are available analytically.

Each of these mutual informations is a sum of the ΦID atoms whose dependency modes are compatible with the corresponding source and target subsets. For example,

$$\begin{aligned} I(Z_t^1; Z_{t'}^1) &= I_\partial^{R\to R} + I_\partial^{R\to U_1} + I_\partial^{U_1\to R} + I_\partial^{U_1\to U_1}, \\ I(Z_t^1; \mathbf{Z}_{t'}) &= I_\partial^{R\to R} + I_\partial^{R\to U_1} + I_\partial^{R\to U_2} + I_\partial^{R\to S} \\ &\quad + I_\partial^{U_1\to R} + I_\partial^{U_1\to U_1} + I_\partial^{U_1\to U_2} + I_\partial^{U_1\to S}. \end{aligned} \tag{S10}$$

The remaining source-target mutual informations follow analogously. Finally, the TDMI decomposes into all sixteen atoms,

$$I(\mathbf{Z}_t; \mathbf{Z}_{t'}) = \sum_{\alpha,\beta \in \mathcal{A}} I_\partial^{\alpha \to \beta}. \tag{S11}$$

*Information-geometric synergy*

The nine mutual information equations do not uniquely determine the sixteen atoms. The seven remaining constraints are provided by our geometric measures of synergy introduced in Ref.[9]. These quantities are obtained by identifying the component of statistical dependence that cannot be accounted for by lower-order source-target pathways.

Consider, for example, three jointly distributed variable sets $\mathbf{A}_1$, $\mathbf{A}_2$, and $\mathbf{B}$. To isolate their synergistic dependence, we construct distributions in which the dependency between one source and the target is explained entirely through the other source. For Gaussian distributions, these constrained distributions can be obtained directly from the covariance structure by conditioning over specified components. The resulting distributions form the information-geometric submanifolds associated with the corresponding conditional-independence constraints. Within this approach, synergy is identified as the part of a joint dependency that cannot be explained by lower-order source-target pathways.

Here, we first describe the construction for three jointly distributed variable sets, which will then be applied to the relevant marginals of the dynamical system.

Consider a joint distribution

$$P = P(\mathbf{A}_1, \mathbf{A}_2, \mathbf{B}), \tag{S12}$$

where $\mathbf{A}_1$ and $\mathbf{A}_2$ are two source groups and $\mathbf{B}$ is a target group. We construct two constrained distributions by projecting $P$ onto submanifolds in which the target is conditionally independent of one source given the other. Specifically, let $\bar{P}^{(1)}$ denote the projection onto the family satisfying

$$\mathbf{A}_2 \perp \mathbf{B} \mid \mathbf{A}_1, \tag{S13}$$

and let $\bar{P}^{(2)}$ denote the corresponding projection satisfying

$$\mathbf{A}_1 \perp \mathbf{B} \mid \mathbf{A}_2. \tag{S14}$$

These projections represent distributions in which the dependence between the target and one source can be completely accounted for by the other source. The KL divergence from the original distribution to either projection therefore quantifies the amount of statistical structure removed by imposing the corresponding conditional-independence constraint.

Neither projection alone captures all dependencies that can be explained without a genuinely joint contribution from $\mathbf{A}_1$ and $\mathbf{A}_2$. To account for both possibilities, we consider the $e$-flat interpolation between the two projections. In the exponential-family representation, this interpolation takes the form

$$P_{\text{interp}}(r) = \frac{1}{\mathcal{Z}(r)} \bar{P^{(1)}}^{1-r} \bar{P^{(2)}}^{r}, \tag{S15}$$

where $r \in \mathbb{R}$ and $\mathcal{Z}(r)$ is a normalising constant. The resulting family contains the distributions obtained by continuously interpolating between the two alternative lower-order explanations of the observed dependency.

The synergistic information is then defined as the residual divergence between the original distribution and the closest member of this family:

$$S(\mathbf{A}_1, \mathbf{A}_2; \mathbf{B}) = \min_{r \in \mathbb{R}} D_{\text{KL}} \left( P \,\|\, P_{\text{interp}}(r) \right). \tag{S16}$$

Thus, synergy corresponds to the statistical structure that cannot be reproduced by any interpolation between models in which the target-source dependency is mediated through one source or the other.

For the bivariate dynamical system, this construction can be applied to the appropriate three-variable marginals to obtain the order-three synergies:

$$\begin{aligned} S_3(Z_t^1; \mathbf{Z}_{t'}) &= I_\partial^{R \to S} + I_\partial^{U_1 \to S}, \\ S_3(Z_t^2; \mathbf{Z}_{t'}) &= I_\partial^{R \to S} + I_\partial^{U_2 \to S}, \\ S_3(\mathbf{Z}_t; Z_{t'}^1) &= I_\partial^{S \to R} + I_\partial^{S \to U_1}, \\ S_3(\mathbf{Z}_t; Z_{t'}^2) &= I_\partial^{S \to R} + I_\partial^{S \to U_2}. \end{aligned} \tag{S17}$$

Two additional order-three synergies are obtained by treating the complete past or future as a single multivariate group:

$$\begin{aligned} S_3(\mathbf{Z}_t; Z^1_{t'}, Z^2_{t'}) &= I_\partial^{S\to R} + I_\partial^{S\to U_1} + I_\partial^{S\to U_2} + I_\partial^{S\to S}, \\ S_3(Z^1_t, Z^2_t; \mathbf{Z}_{t'}) &= I_\partial^{R\to S} + I_\partial^{U_1\to S} + I_\partial^{U_2\to S} + I_\partial^{S\to S}. \end{aligned} \tag{S18}$$

The final constraint must isolate the contribution that is synergistic in both the past and the future. To construct this, we consider the four source-target pairs $(Z^1_t, Z^1_{t'})$, $(Z^1_t, Z^2_{t'})$, $(Z^2_t, Z^1_{t'})$, and $(Z^2_t, Z^2_{t'})$. For each pair, we construct a projected distribution in which the dependency between the complementary source and target components is removed, leaving the corresponding source-target pathway as the only direct pathway. Denote these four projections by

$$\bar{P}^{11}, \qquad \bar{P}^{12}, \qquad \bar{P}^{21}, \qquad \bar{P}^{22}. \tag{S19}$$

As in the three-variable construction, no single projection captures all dependencies that can be explained without a genuinely four-way contribution. We therefore consider their joint $e$-flat interpolation,

$$P_{\text{interp}} = \frac{1}{\mathcal{Z}(\mathbf{w})} \left(\bar{P}^{11}\right)^{w_{11}} \left(\bar{P}^{12}\right)^{w_{12}} \left(\bar{P}^{21}\right)^{w_{21}} \left(\bar{P}^{22}\right)^{w_{22}}, \qquad \sum_{i,j=1}^{2} w_{ij} = 1, \tag{S20}$$

where $\mathcal{Z}(\mathbf{w})$ ensures normalisation. This family contains the distributions that can be explained by combinations of pairwise and three-way source-target pathways, while excluding the component that is simultaneously synergistic across the two source components and the two target components.

The highest-order synergy is therefore given by the minimum KL divergence from the original distribution to this family:

$$S_4(\mathbf{Z}_t; \mathbf{Z}_{t'}) = \min_{\mathbf{w}} D_{\text{KL}}\left(P \,\|\, P_{\text{interp}}(\mathbf{w})\right). \tag{S21}$$

For the bivariate ΦID, this quantity corresponds exactly to the atom that is synergistic in both the past and the future,

$$S_4(\mathbf{Z}_t; \mathbf{Z}_{t'}) = I_\partial^{S\to S}. \tag{S22}$$

This establishes the link between the information-geometric definition of synergy and the ΦID lattice, with the geometric construction supplying precisely the additional constraints needed to resolve the atoms that cannot be determined from mutual information alone.

*Gaussian implementation*

For the Gaussian processes considered throughout this work, all quantities required by the information-geometric construction can be obtained directly from the covariance matrix of the lagged system. Let

$$\Sigma = \text{Cov}\left(Z^1_t, Z^2_t, Z^1_{t'}, Z^2_{t'}\right) \tag{S23}$$

denote the covariance matrix of the joint past–future distribution

$$P = \mathcal{N}(\mathbf{0}, \Sigma). \tag{S24}$$

The covariance matrices of all marginal distributions appearing in the decomposition are obtained by selecting the corresponding rows and columns of $\Sigma$.

For two $d$-dimensional Gaussian distributions

$$P = \mathcal{N}(\boldsymbol{\mu}_P, \Sigma_P), \qquad Q = \mathcal{N}(\boldsymbol{\mu}_Q, \Sigma_Q), \tag{S25}$$

the KL divergence reads

$$D_{\text{KL}}(P\|Q) = \frac{1}{2}\left[\text{tr}\left(\Sigma_Q^{-1}\Sigma_P\right) + (\boldsymbol{\mu}_Q - \boldsymbol{\mu}_P)^\top \Sigma_Q^{-1}(\boldsymbol{\mu}_Q - \boldsymbol{\mu}_P) - d + \ln\frac{\det\Sigma_Q}{\det\Sigma_P}\right]. \tag{S26}$$

All time series are standardised before estimating the joint distribution, so that the relevant distributions have zero mean. The KL divergence therefore reduces to

$$D_{\text{KL}}(P\|Q) = \frac{1}{2}\left[\text{tr}\left(\Sigma_Q^{-1}\Sigma_P\right) - d + \ln\frac{\det\Sigma_Q}{\det\Sigma_P}\right]. \tag{S27}$$

The order-three geometric synergies used to constrain the ΦID can then be evaluated using the Gaussian information-geometric construction of Ref.[10]. Here, we briefly outline the approach.

*a. Order-three geometric synergies.* Consider a jointly Gaussian triple

$$(\boldsymbol{A}_1, \boldsymbol{A}_2; \boldsymbol{B}) \tag{S28}$$

with zero mean and covariance matrix $\Sigma^{(3)}$. Here $\boldsymbol{A}_1$ and $\boldsymbol{A}_2$ are the two source groups and $\boldsymbol{B}$ is the target group.

Following the reasoning above, for each source we construct a projection in which the target is conditionally independent of the other source given that source. Specifically, for $k \in \{1, 2\}$, let $\bar{P}^{(k)}$ denote the projection satisfying

$$\boldsymbol{A}_{-k} \perp \boldsymbol{B} \mid \boldsymbol{A}_k, \tag{S29}$$

where $-k$ denotes the source complementary to $k$. In the Gaussian case, this constraint is imposed by replacing the cross-covariance between $\boldsymbol{A}_{-k}$ and $\boldsymbol{B}$ by the covariance implied by conditioning on $\boldsymbol{A}_k$:

$$\bar{\Sigma}^{(k)}_{\boldsymbol{A}_{-k}\boldsymbol{B}} = \Sigma_{\boldsymbol{A}_{-k}\boldsymbol{A}_k} \Sigma^{-1}_{\boldsymbol{A}_k\boldsymbol{A}_k} \Sigma_{\boldsymbol{A}_k\boldsymbol{B}}, \tag{S30}$$

with the symmetric transpose block set accordingly and all other covariance blocks left unchanged. Equation (S30) follows directly from the Gaussian conditional-independence relation: the residual covariance between $\boldsymbol{A}_{-k}$ and $\boldsymbol{B}$ after conditioning on $\boldsymbol{A}_k$ is set to zero, and thus the resulting covariance matrix $\bar{\Sigma}^{(k)}$ defines the unique Gaussian projection onto the corresponding conditional-independence submanifold.

The two projections represent the two alternative lower-order explanations of the source–target dependency: one in which $\boldsymbol{A}_2$ contributes only through $\boldsymbol{A}_1$, and one in which $\boldsymbol{A}_1$ contributes only through $\boldsymbol{A}_2$. To include both possibilities, we consider the $e$-geodesic connecting the two projections. For a zero-mean Gaussian distribution, the natural parameter is the precision matrix, so this geodesic is linear in precision space. Defining

$$K_k = \left(\bar{\Sigma}^{(k)}\right)^{-1}, \tag{S31}$$

the interpolating family is

$$K(u) = (1-u)K_1 + uK_2, \qquad P_u = \mathcal{N}\left(\mathbf{0}, K(u)^{-1}\right), \tag{S32}$$

where $u$ is restricted to values for which $K(u)$ remains positive definite, which we denote as $\mathcal{U}$. The order-three synergy is then the minimum KL divergence between the original distribution and this family,

$$S(\boldsymbol{A}_1, \boldsymbol{A}_2; \boldsymbol{B}) = \min_{u \in \mathcal{U}} D_{\mathrm{KL}}\left(P \| P_u\right). \tag{S33}$$

For the Gaussian case, this divergence simplifies substantially. Substituting $P_u = \mathcal{N}(0, K(u)^{-1})$ into the Gaussian KL divergence gives

$$D_{\mathrm{KL}}(P \| P_u) = \frac{1}{2}\left[\mathrm{tr}\left(K(u)\Sigma^{(3)}\right) - d + \ln \frac{\det K(u)^{-1}}{\det \Sigma^{(3)}}\right], \tag{S34}$$

where $d$ is the total dimensionality of the three groups. Because each projection preserves all covariance blocks except the one constrained by the corresponding conditional independence, its precision matrix satisfies

$$\mathrm{tr}\left(K_k \Sigma^{(3)}\right) = d. \tag{S35}$$

Linearity of the trace therefore gives

$$\mathrm{tr}\left(K(u)\Sigma^{(3)}\right) = d \tag{S36}$$

throughout the admissible interpolation. Hence the trace and dimensional terms cancel exactly, and the synergy can be evaluated as

$$S(\boldsymbol{A}_1, \boldsymbol{A}_2; \boldsymbol{B}) = \min_{u \in \mathcal{U}} \frac{1}{2} \ln \frac{\det K(u)^{-1}}{\det \Sigma^{(3)}}. \tag{S37}$$

In the present bivariate dynamical system, this construction is applied to four three-variable marginals, with either the past or future consisting of a single component, yielding Eq. (S17). The same construction can also be applied when one side consists of the complete bivariate group, giving in turn Eq. (S18). These six geometric synergies provide the six order-three constraints required by the ΦID system. The remaining constraint is supplied by the order-four construction below.

*b. Order-four synergy.* The four-variable construction is obtained by applying the same geometric principle to the complete lagged distribution. Consider one of the four source–target pairs,

$$(Z_t^i, Z_{t'}^j), \qquad i, j \in \{1, 2\}, \tag{S38}$$

and denote the complementary variables by

$$A = Z_t^{-i}, \qquad B = Z_{t'}^{-j}, \tag{S39}$$

while retaining

$$\mathbf{C} = \left(Z_t^i, Z_{t'}^j\right). \tag{S40}$$

We construct a projected distribution in which the dependence between the complementary variables $A$ and $B$ is entirely mediated by the retained pair $\mathbf{C}$. Equivalently, the projection satisfies

$$A \perp B \mid \mathbf{C}. \tag{S41}$$

This removes the direct statistical interaction between the two complementary variables while preserving all lower-order interactions involving the retained source–target pair.

For a Gaussian distribution, this projection has a particularly simple expression. If $\Sigma$ is partitioned according to $(A, \mathbf{C}, B)$, the covariance matrix of the projected distribution, denoted by $\bar{\Sigma}^{(ij)}$, is identical to $\Sigma$ in all blocks except the $A$–$B$ cross-covariance, which is replaced by the value implied by the conditional-independence constraint:

$$\bar{\Sigma}_{AB}^{(ij)} = \Sigma_{A\mathbf{C}}\Sigma_{\mathbf{CC}}^{-1}\Sigma_{\mathbf{C}B}, \qquad \bar{\Sigma}_{BA}^{(ij)} = \left(\bar{\Sigma}_{AB}^{(ij)}\right)^{\top}. \tag{S42}$$

Thus, the four choices $(i, j) \in \{1, 2\}^2$ give four projected covariance matrices,

$$\bar{\Sigma}^{(11)}, \quad \bar{\Sigma}^{(12)}, \quad \bar{\Sigma}^{(21)}, \quad \bar{\Sigma}^{(22)}, \tag{S43}$$

each corresponding to one of the four possible conditionings.

As discussed above, the four projections provide alternative lower-order explanations of the statistical dependencies in the system. Importantly, none of them individually isolates the genuinely four-way contribution: each projection retains a different source–target pathway and therefore remains capable of accounting for pairwise and three-way dependencies. We therefore consider the $e$-flat family generated by these four projections. For Gaussian distributions, $e$-geodesics are linear in the natural parameter, which for a zero-mean Gaussian is the precision matrix. Defining

$$K_{ij} = \left(\bar{\Sigma}^{(ij)}\right)^{-1}, \tag{S44}$$

the interpolating family is therefore

$$K(\boldsymbol{\lambda}) = \lambda_{11}K_{11} + \lambda_{12}K_{12} + \lambda_{21}K_{21} + \lambda_{22}K_{22}, \qquad \sum_{i,j=1}^{2} \lambda_{ij} = 1, \tag{S45}$$

with

$$Q_{\boldsymbol{\lambda}} = \mathcal{N}\left(\mathbf{0}, K(\boldsymbol{\lambda})^{-1}\right). \tag{S46}$$

The admissible parameter set consists of those $\boldsymbol{\lambda}$ for which $K(\boldsymbol{\lambda})$ remains positive definite. We denote this set by $\Lambda$.

The order-four synergy is the residual information that cannot be represented by any distribution in this family. It is therefore obtained as the minimum KL divergence between the original distribution and the interpolating family,

$$S_4 = \min_{\boldsymbol{\lambda}\in\Lambda} D_{\mathrm{KL}}\left(P \,\|\, Q_{\boldsymbol{\lambda}}\right). \tag{S47}$$

For the Gaussian case, this optimisation has a useful simplification. Substituting $Q_{\boldsymbol{\lambda}} = \mathcal{N}(\mathbf{0}, K(\boldsymbol{\lambda})^{-1})$ into Eq. (S27) gives

$$D_{\mathrm{KL}}\left(P \,\|\, Q_{\boldsymbol{\lambda}}\right) = \frac{1}{2}\left[\mathrm{tr}\left(K(\boldsymbol{\lambda})\Sigma\right) - 4 + \ln\frac{\det K(\boldsymbol{\lambda})^{-1}}{\det\Sigma}\right]. \tag{S48}$$

Each projection preserves all covariance blocks except the single complementary $A$–$B$ block, and its precision matrix satisfies

$$\operatorname{tr}(K_{ij}\Sigma) = 4. \tag{S49}$$

Because the trace is linear,

$$\operatorname{tr}(K(\boldsymbol{\lambda})\Sigma) = \sum_{i,j} \lambda_{ij} \operatorname{tr}(K_{ij}\Sigma) = 4, \tag{S50}$$

for every admissible $\boldsymbol{\lambda}$. Consequently, the trace and dimensional terms cancel exactly, leaving the particularly simple expression

$$S_4 = \min_{\boldsymbol{\lambda}\in\Lambda} \frac{1}{2} \ln \frac{\det K(\boldsymbol{\lambda})^{-1}}{\det \Sigma}. \tag{S51}$$

Thus, once the covariance matrix $\Sigma$ has been estimated, computing $S_4$ requires only the construction of the four projected covariance matrices, their inverses, and a low-dimensional optimisation over the admissible interpolation weights. As explained above, this provides the final constraint over $I_\partial^{S\to S}$, which closes the ΦID system of sixteen equations.

*Solving the decomposition*

The nine mutual information constraints and seven geometric synergy constraints together form a closed linear system for the sixteen ΦID atoms. We solve this system for each pair of brain regions and subsequently combine atoms related by exchange of the two components. This gives the ten symmetric information modes introduced in the main text:

$$\begin{aligned}
R\to R &= I_\partial^{R\to R},\\
R\to U &= I_\partial^{R\to U_1} + I_\partial^{R\to U_2},\\
R\to S &= I_\partial^{R\to S},\\
U\to R &= I_\partial^{U_1\to R} + I_\partial^{U_2\to R},\\
U_X\to U_X &= I_\partial^{U_1\to U_1} + I_\partial^{U_2\to U_2},\\
U_X\to U_Y &= I_\partial^{U_1\to U_2} + I_\partial^{U_2\to U_1},\\
U\to S &= I_\partial^{U_1\to S} + I_\partial^{U_2\to S},\\
S\to R &= I_\partial^{S\to R},\\
S\to U &= I_\partial^{S\to U_1} + I_\partial^{S\to U_2},\\
S\to S &= I_\partial^{S\to S}.
\end{aligned} \tag{S52}$$

Here, the two unique components are grouped whenever their distinction is irrelevant to the quantity of interest. In particular, $U_X\to U_X$ denotes unique information that remains associated with the same component, whereas $U_X\to U_Y$ denotes unique information that is transferred between components.

*Pointwise decomposition*

The decomposition described above yields information quantities defined as expectations over the joint distribution of the past and future states. These global quantities therefore characterise the average information dynamics of the system. To obtain a time-resolved description, we evaluate the same information-geometric construction pointwise, assigning an information value to each observed transition $(\mathbf{z}_t, \mathbf{z}_{t'})$.

The key observation is that every quantity entering the linear system for the global ΦID can be written as an expectation of a corresponding pointwise quantity. Thus, once the joint probability distribution and the geometric projections have been estimated, the global decomposition can be converted into a local decomposition simply by replacing each expectation by its pointwise log-density ratio. Importantly, the underlying probability models are estimated once from the complete time series and are not re-estimated separately at each time point. The temporal variation of the resulting information atoms therefore arises from the particular states visited by the system, rather than from changes in the estimated probability model.

For example, the pointwise TDMI reads

$$i(\mathbf{z}_t; \mathbf{z}_{t'}) = \ln \frac{p(\mathbf{z}_t, \mathbf{z}_{t'})}{p(\mathbf{z}_t)p(\mathbf{z}_{t'})}, \tag{S53}$$

whose temporal average recovers the usual TDMI,

$$I(\mathbf{Z}_t; \mathbf{Z}_{t'}) = \mathbb{E}_P[i(\mathbf{z}_t; \mathbf{z}_{t'})]. \tag{S54}$$

The same principle applies to the mutual information terms involving marginal source and target groups, as well as to the KL divergences used to define the geometric synergies.

For the Gaussian distributions considered here, the pointwise KL divergence between two distributions

$$P = \mathcal{N}(\boldsymbol{\mu}_P, \Sigma_P), \qquad Q = \mathcal{N}(\boldsymbol{\mu}_Q, \Sigma_Q), \tag{S55}$$

is given by the log-density ratio

$$d_{\mathrm{KL}}(\mathbf{z}; P|Q) = \ln \frac{p(\mathbf{z})}{q(\mathbf{z})}. \tag{S56}$$

Writing the Gaussian log densities explicitly gives

$$d_{\mathrm{KL}}(\mathbf{z}; P|Q) = \frac{1}{2}\Big[\ln \frac{\det \Sigma_Q}{\det \Sigma_P} - (\mathbf{z} - \boldsymbol{\mu}_P)^\top \Sigma_P^{-1}(\mathbf{z} - \boldsymbol{\mu}_P) + (\mathbf{z} - \boldsymbol{\mu}_Q)^\top \Sigma_Q^{-1}(\mathbf{z} - \boldsymbol{\mu}_Q)\Big]. \tag{S57}$$

For the standardised, zero-mean variables used here this reduces to

$$d_{\mathrm{KL}}(\mathbf{z}; P|Q) = \frac{1}{2}\left[\ln \frac{\det \Sigma_Q}{\det \Sigma_P} + \mathbf{z}^\top \left(\Sigma_Q^{-1} - \Sigma_P^{-1}\right)\mathbf{z}\right]. \tag{S58}$$

Taking the expectation of Eq. (S58) under $P$ recovers the usual Gaussian KL divergence in Eq. (S27). Unlike the global KL divergence, however, its pointwise counterpart can take either positive or negative values. It therefore represents the signed information contribution of a particular observation to the average divergence. This expression allows the geometric synergy measures to be evaluated locally without modifying the underlying optimisation procedure.

Using the quantities introduced above, for the order-three construction, let $P$ denote the original Gaussian distribution and let

$$P_u = \mathcal{N}\left(\mathbf{0}, K(u)^{-1}\right) \tag{S59}$$

denote the interpolating distribution obtained from the two projected models, with $u^*$ chosen by the global optimisation

$$u^* = \underset{u\in\mathcal{U}}{\mathrm{argmin}}, D_{\mathrm{KL}}(P|P_u). \tag{S60}$$

The corresponding pointwise order-three synergy is then obtained by evaluating the log-density ratio between the original distribution and the optimally interpolated distribution,

$$s_3(\mathbf{z}) = d_{\mathrm{KL}}\left(\mathbf{z}; P|P_{u^*}\right). \tag{S61}$$

Using Eq. (S58), this becomes

$$s_3(\mathbf{z}) = \frac{1}{2}\left[\ln \frac{\det K(u^*)^{-1}}{\det \Sigma^{(3)}} + \mathbf{z}^\top \left(K(u^*) - (\Sigma^{(3)})^{-1}\right)\mathbf{z}\right]. \tag{S62}$$

The global order-three synergy is recovered by averaging this quantity over observations,

$$S_3(\mathbf{z}) = \mathbb{E}_P[s_3(\mathbf{z})]. \tag{S63}$$

Thus, the same interpolating distribution that defines the global geometric synergy also defines its local counterpart; only the evaluation of the KL divergence changes from an expectation over the distribution to a log-density ratio evaluated at the observed state.

The order-four construction follows identically. Let

$$K^* = K(\boldsymbol{\lambda}^*), \qquad \boldsymbol{\lambda}^* = \underset{\boldsymbol{\lambda}\in\Lambda}{\mathrm{argmin}} D_{\mathrm{KL}}\left(P|Q_{\boldsymbol{\lambda}}\right), \tag{S64}$$

where

$$Q_{\boldsymbol{\lambda}} = \mathcal{N}\left(\mathbf{0}, K(\boldsymbol{\lambda})^{-1}\right). \tag{S65}$$

The local order-four synergy is then

$$s_4(\mathbf{z}_t, \mathbf{z}_{t'}) = d_{\mathrm{KL}}\left(\mathbf{z}_t, \mathbf{z}_{t'}; P|Q_{\boldsymbol{\lambda}^*}\right), \tag{S66}$$

or, explicitly,

$$s_4(\mathbf{z}_t, \mathbf{z}_{t'}) = \frac{1}{2}\left[\ln\frac{\det(K^*)^{-1}}{\det\Sigma} + (\mathbf{z}_t\ \mathbf{z}_{t'})^\top\left(K^* - \Sigma^{-1}\right)(\mathbf{z}_t\ \mathbf{z}_{t'})\right]. \tag{S67}$$

As expected, its temporal average satisfies

$$S_4(\mathbf{Z_t}; \mathbf{Z_{t'}}) = \mathbb{E}_P[s_4(\mathbf{z}_t, \mathbf{z}_{t'})]. \tag{S68}$$

Having obtained the pointwise versions of the nine mutual-information constraints and seven geometric synergy constraints, we solve the same $16\times 16$ linear system independently at each time point. Specifically, if $\mathbf{c}$ denotes the vector of the sixteen global constraints and $\mathbf{x}$ the vector of the sixteen ΦID atoms, the global decomposition can be written as

$$\mathbf{c} = M\mathbf{x}, \tag{S69}$$

where $M$ is the fixed coefficient matrix determined solely by the ΦID lattice. At each time point, the corresponding local quantities define

$$\mathbf{c}_t = M\mathbf{x}_t, \tag{S70}$$

and hence

$$\mathbf{x}_t = M^{-1}\mathbf{c}_t. \tag{S71}$$

The coefficient matrix is identical at every time point; only the vector of observed information quantities changes. The resulting $\mathbf{x}_t$ therefore contains the local value of every ΦID atom for the particular transition from $\mathbf{z}_t$ to $\mathbf{z}_{t'}$.

Because the linear system is the same as that used for the global decomposition, temporal expectation commutes with the solution of the system:

$$\mathbb{E}_P[\mathbf{x}_t] = M^{-1}\mathbb{E}_P[\mathbf{c}_t]. \tag{S72}$$

Consequently, the temporal average of each local ΦID atom is equal to its corresponding global value, up to finite-sample estimation effects.

#### S5. SUPPLEMENTARY FIGURES

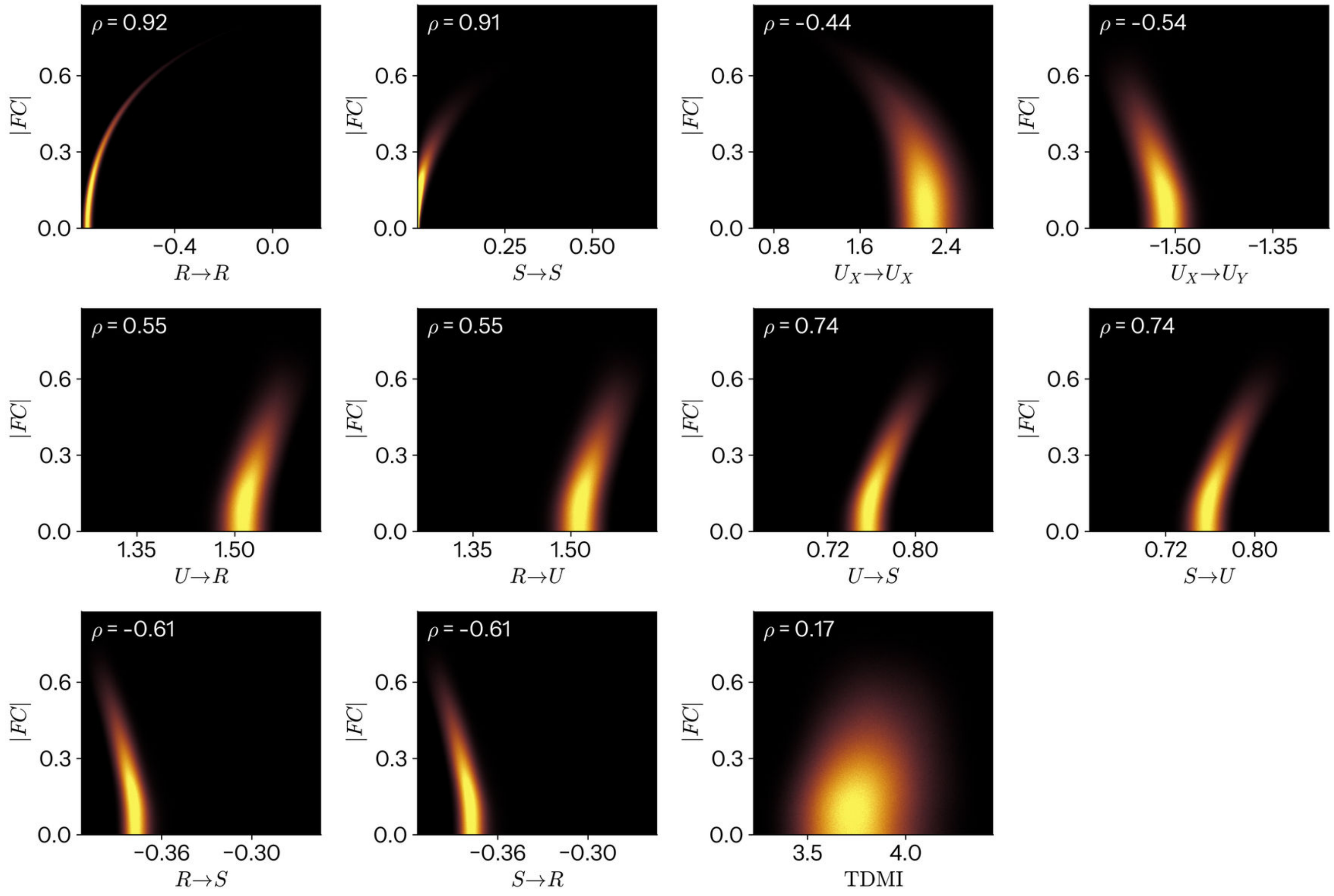


FIG. S1. **Relationship between modes of information dynamics and traditional functional connectivity.** Joint distribution of each mode's edge weight against the absolute Pearson functional connectivity $|FC|$ of the same edge, pooled over all 47,717,752 edge observations of the cohort. Panels are two-dimensional histograms. The final panel gives the time-delayed mutual information (TDMI) that the atoms sum to.

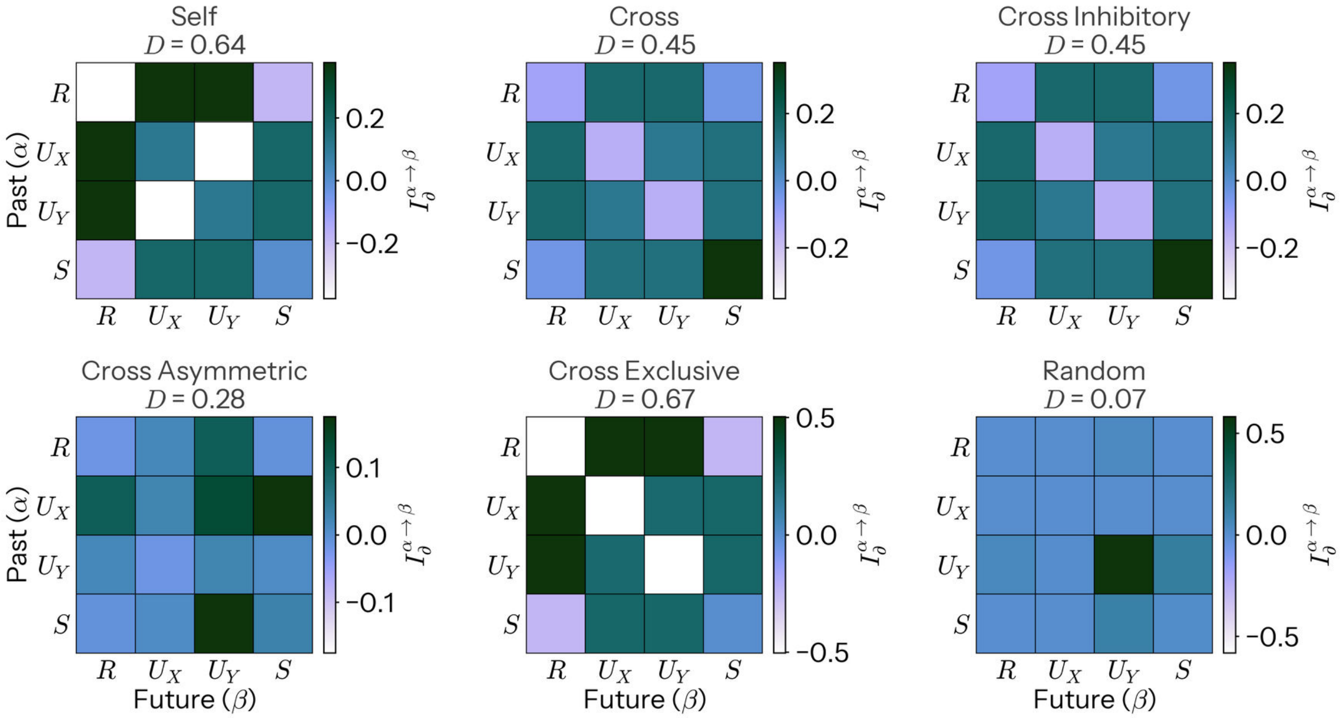


FIG. S2. **Demonstration of computational diversity in toy systems.** Two-node vector-autoregressive (VAR(1)) system, whose dynamics are set by a transition matrix $A$ with self-coupling $a$ (diagonal) and cross-coupling $b$ (off-diagonal). Each panel gives one system's atoms as a matrix, rows the mode the information occupied in the past ($\alpha$) and columns the mode it occupies in the future ($\beta$), so that the diagonal is persistence and the off-diagonal conversion, with $D$ stated above each panel. The systems are *Self* ($a = 0.7$, $b = 0$), *Cross* ($a = 0.3$, $b = 0.6$), *Cross Inhibitory* ($a = 0.3$, $b = -0.6$), *Cross Asymmetric* ($a = 0.4$, with $b = 0.6$ on $X \to Y$ only), *Cross Exclusive* ($a = 0$, $b = 0.8$) and *Random* (entries drawn from a standard normal and rescaled to a maximum singular value of 0.85). As expected, systems that concentrate their information in a few atoms have low diversity (e.g. the random system), whereas those that spread it broadly have high diversity (e.g. the heavily cross-coupled system). Because $D$ depends only on the magnitudes of the atoms, the excitatory and inhibitory cross-coupled systems yield identical diversity and TDMI despite their opposite coupling sign.

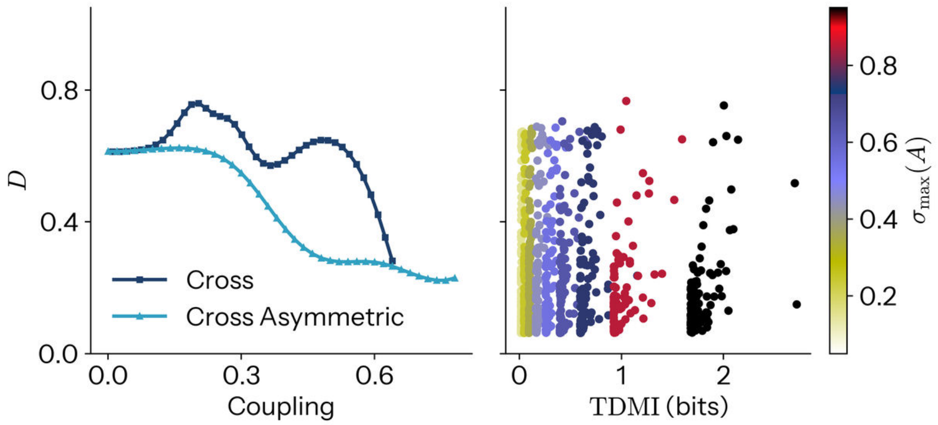


FIG. S3. **Computational diversity diminishes at strong coupling, and is unrelated to TDMI.** *Left*: computational diversity $D$ of two families of two-node VAR(1) systems as a function of coupling strength. Strengthening the cross-coupling between the two variables, whether symmetric (*Cross*; $a = 0.3$) or asymmetric ($X \to Y$ only, *Cross Asymmetric*; $a = 0.4$), drives $D$ down as information becomes concentrated in progressively fewer atoms. *Right*: $D$ against total time-delayed mutual information (TDMI) for 1,000 random transition matrices spanning a range of largest singular values $\sigma_{\max}(A)$ (colour). Across the full range of TDMI, $D$ shows no systematic dependence on the total information carried (Pearson $r = -0.1$), confirming that computational diversity is not a proxy for the amount of information in a system.

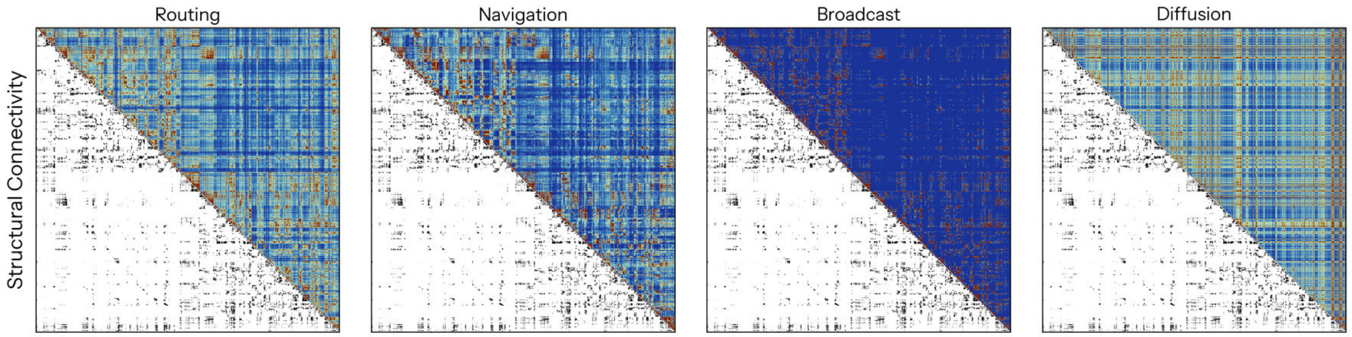


FIG. S4. **Communication strategies on the structural connectome.** The four models of polysynaptic signalling regressed onto each network of information dynamics in Fig. 2b, ordered from centralised to decentralised: shortest-path efficiency (Routing), navigation efficiency (Navigation), communicability (Broadcast) and diffusion efficiency (Diffusion) (see Methods). Each panel shows the model in its upper triangle against the structural connectome it is computed from in its lower triangle, so that the two can be compared edge for edge.

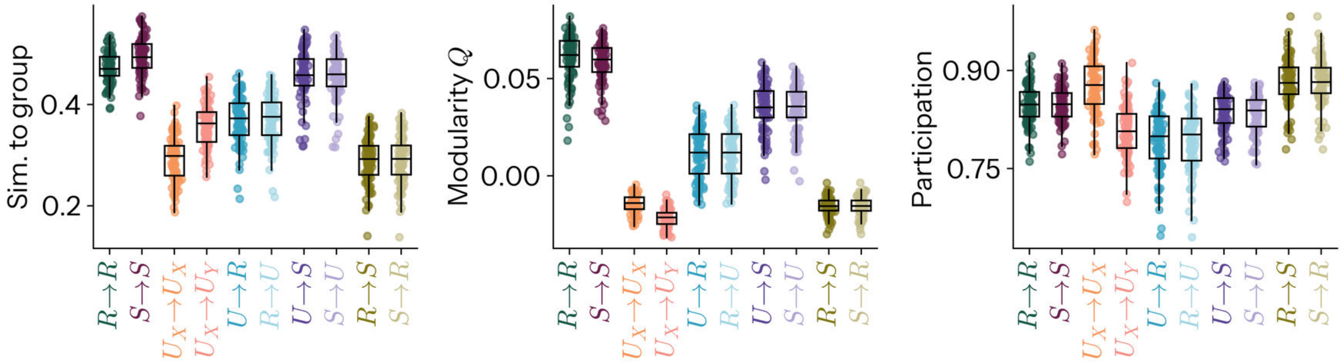


FIG. S5. **Replication of topology with communities defined as individual rather than group level.** The modularity and participation reported in Fig. 2d are computed on a group partition applied to every participant. Here each participant's network is instead partitioned on its own, and the same quantities recomputed from those individual communities (see Methods). *Left*: adjusted mutual information between a participant's own partition and the group partition of the same mode (mean over all modes and participants $0.385 \pm 0.088$, range 0.138 to 0.574). *Middle*: modularity $Q$ of each participant's own partition. *Right*: participation coefficient under that partition, averaged over regions. Points are participants and boxes their distribution. The ordering recovered is the same one the group partitions give: $R \to R$ and $S \to S$ are the most modular and the most reproducible against the group, $R \to S$ and $S \to R$ the least, so the topological division of labour reported in the main text is not an artefact of imposing a single partition on everyone. Contrasting each mode against the mean of the other nine, 29 of the 30 mode by measure cells survive $p_{\mathrm{FDR}} < 0.05$.

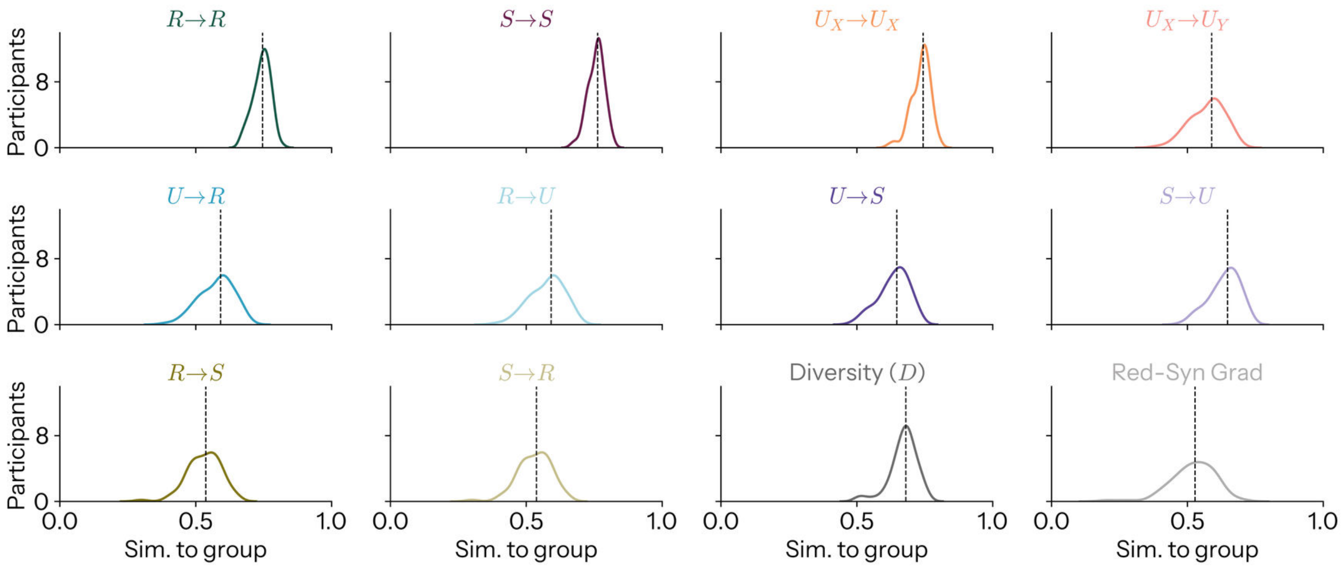


FIG. S6. **Similarity of individual networks to group.** Every participant is positively and substantially similar to the group in every quantity (minimum over all participants and quantities $r = 0.21$, maximum $r = 0.82$), so the group maps of Fig. 1 are representative rather than an average over dissimilar individuals. The modes nonetheless differ in how typical they are, and they do so in the same direction as their identifiability (Fig. 3b): the storage of information is the most group-typical ($S \to S$, $r = 0.757 \pm 0.030$; $R \to R$, $0.743 \pm 0.032$; $U_X \to U_X$, $0.736 \pm 0.035$) and the interconversion of redundancy and synergy the least ($R \to S$, $S \to R$, $0.531 \pm 0.061$), with the redundancy-to-synergy gradient the most variable quantity of all ($0.522 \pm 0.082$).

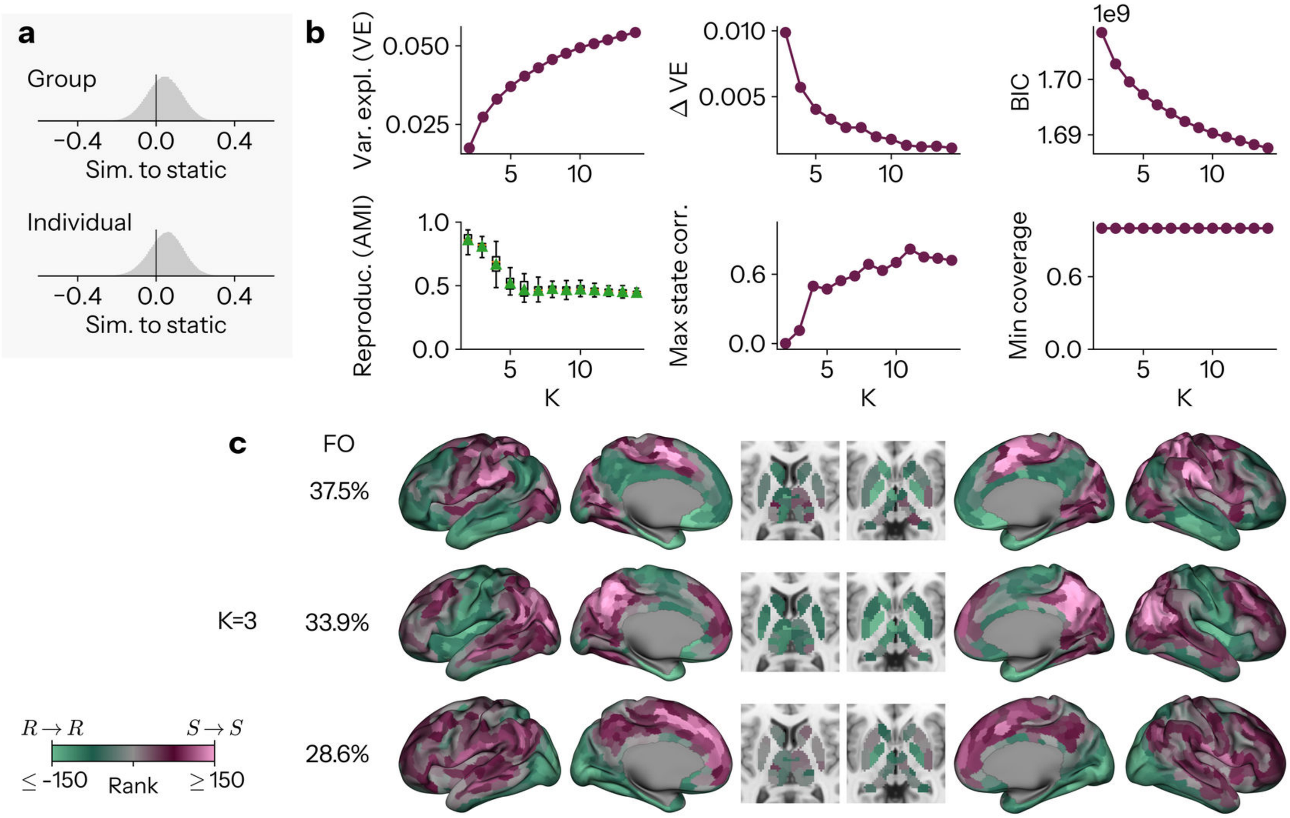


FIG. S7. **Dynamic reconfigurations of the redundancy-to-synergy gradient a** Distribution, pooled across all timepoints and participants, of the spatial correlation between each time-resolved redundancy-to-synergy gradient and the static group-level gradient of Fig. 1e (top) and the participant's own static gradient (bottom). **b** To show examples of time-resolved redundancy-to-synergy gradients that recurred[11], gradients across subjects were concatenated in time and clustered with K-means using Pearson correlation as the distance metric (so that clusters reflect spatial pattern rather than magnitude). For each candidate $K$, clustering was repeated from 50 random initialisations and the solution with the best separation (lowest within-cluster sum of squares) retained; this whole procedure was repeated 10 times independently, the adjusted mutual information (AMI) computed between every pair of the 10 resulting partitions, and the partition sharing the greatest total AMI with all others taken forward. Panels show, across $K = 2$–$14$: variance explained by clustering (VE) and its increment ($\Delta$VE); the Bayesian information criterion (BIC); reproducibility, quantified as the AMI between the 10 partitions (boxplots; green triangles, means); the maximum absolute correlation between state maps; and the minimum across-participant coverage. K=3 is shown for clarity and due to the fact and higher number of states leads to degeneracy among states (correlations between states exceeding r=0.5) **c** The three group-level states at $K = 3$, each shown as the mean time-resolved gradient over all time points assigned to that state (rank units; green, redundancy-dominant $R \to R$; pink, synergy-dominant $S \to S$; grey, balanced), on cortical surfaces (lateral and medial views, both hemispheres) and subcortical volumes. States are ordered top-to-bottom by fractional occupancy (FO, the mean proportion of time points spent in each state; shown at left).

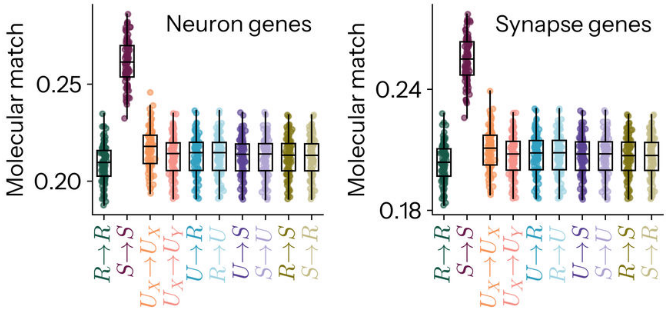


FIG. S8. **Replication of molecular matching with alternatively defined gene sets.** *Left*: Molecular matching with neuron-specific genes from Cahoy et al.[12,13] ($F = 1.79 \times 10^3$, pairwise max $S \rightarrow S$ $p_{\text{FDR}} = 1.26 \times 10^{-52}$) *Right*: Molecular matching with synaptome-specific genes from SynaptomeDB[14] ($F = 1.99 \times 10^3$, pairwise max $S \rightarrow S$ $p_{\text{FDR}} = 4.84 \times 10^{-56}$)

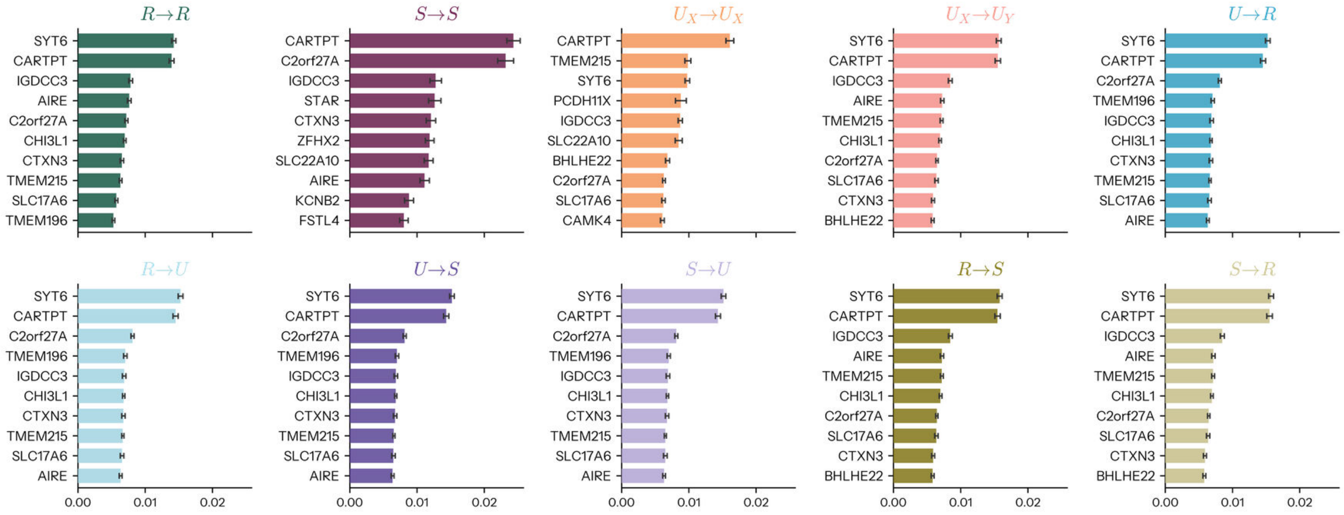


FIG. S9. **Top matching terms for molecular matching** Molecular matching is driven by genes encoding proteins for synaptic vesicle trafficking and neurotransmitter exocytosis like SYT6 (synaptotagmin 6) and SLC17A6 (vesicular glutamate transporter 2), but was interestingly strongest for CARTPT (cocaine- and amphetamine-regulated transcript protein).

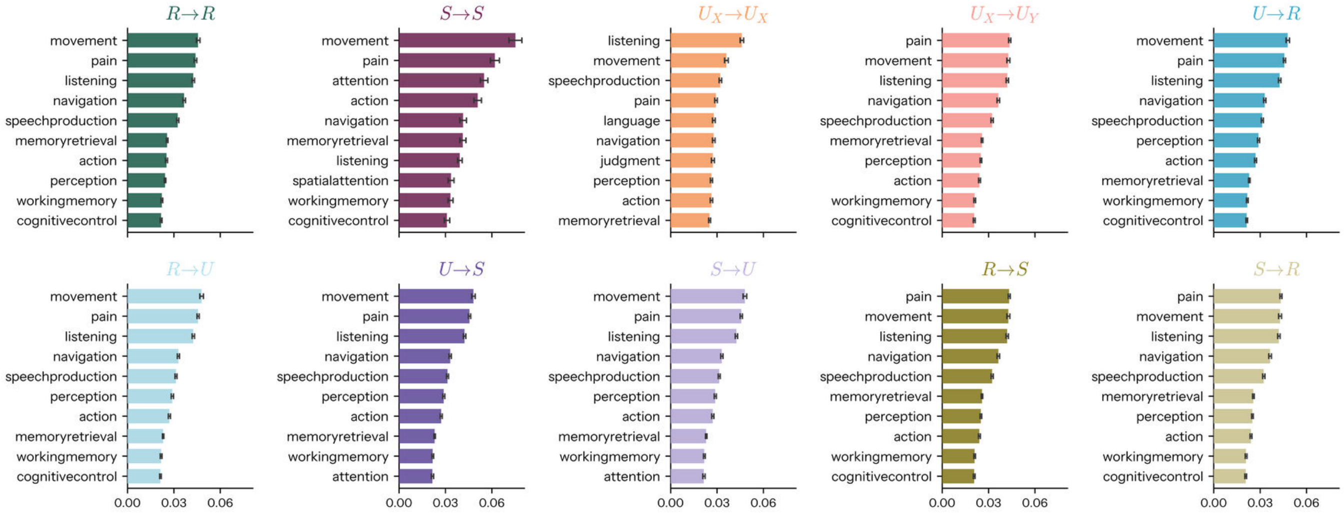


FIG. S10. **Top matching terms for cognitive matching** Cognitive matching was driven by active (e.g movement), affective (e.g. pain) and linguistic (e.g listening) maps.

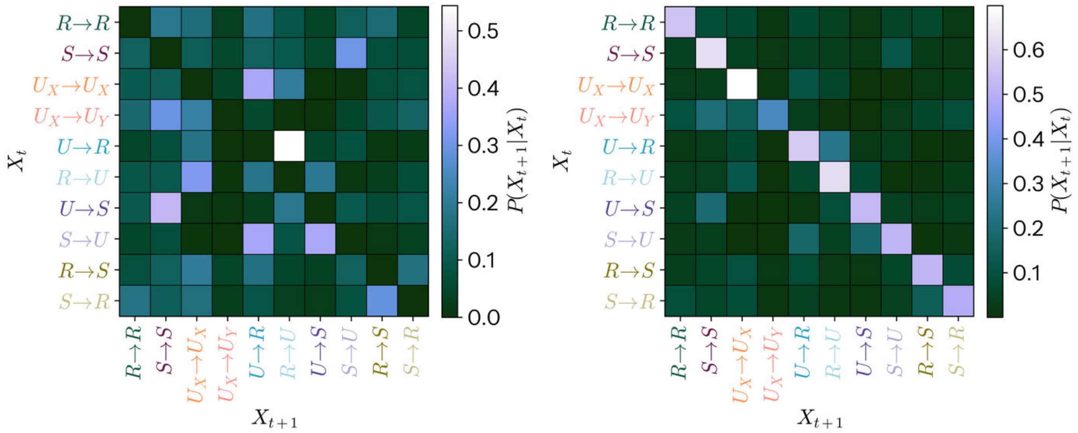


FIG. S11. **Immediate transition structure of global information states.** Group-mean transition probability matrix of the winner-take-all state sequence, $P(X_{t+1}|X_t)$ at one repetition time (1 s), averaged over the 86 participants. *Left*: excluding self-transitions, *Right*: including them.

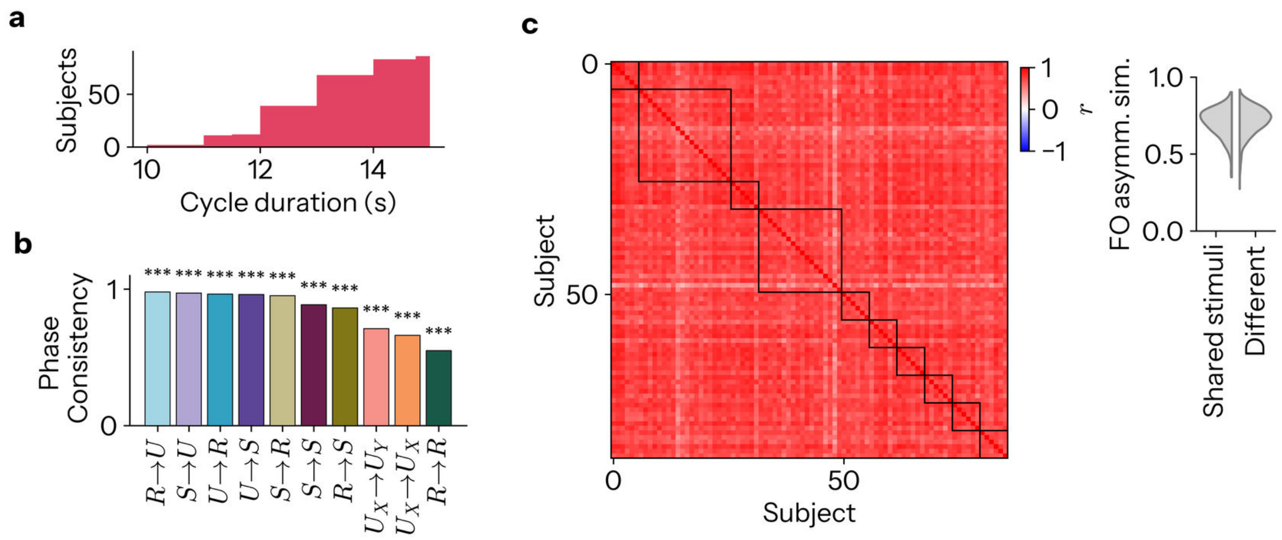


FIG. S12. **Cycle speed and universality a** Cumulative histogram of the duration elapsed in completing a full cycle (median 13.0 s; mean $12.6 \pm 1.1$ s, equivalently $0.080 \pm 0.007$ Hz). Due to the absence of the natural meta-states of van Es *et al.*[15], whose second-level Poisson hidden Markov model times the cycle, we define its duration directly from the intervals that TINDA already partitions. For each participant we take every interval separating successive visits to the same state, pool these revisit intervals across all ten states, and take their median. Since a single traversal of the cycle visits each state once, the typical revisit interval serves as a proxy for the period of one revolution, and its reciprocal for the cycle rate. **b** Phase consistency of each mode across participants. For a given participant, the preferred phase of a mode is obtained by weighting the cycle position of every other mode by the FO asymmetry between the two, and taking the argument of the resulting complex sum, so that a mode is drawn towards the modes it follows and away from those it precedes. The Rayleigh test of circular uniformity is then applied to the 86 phases of each mode, with the mean resultant length $R \in [0, 1]$ (plotted) quantifying their concentration: $R = 0$ when phases are scattered uniformly around the circle, and $R = 1$ when every participant places the mode at the same phase. All ten modes are strongly concentrated (mean $R = 0.85$, all $p < 10^{-12}$). **c** Correlation between the fractional occupancy asymmetry matrices of every pair of participants shows no difference between those observing the same stimuli versus different ones. Left: the pairwise correlation matrix, ordered by film, with black squares delineating the ten films. Right: the same correlations, split by whether the two participants watched the same film or different films. The asymmetry structure is strongly conserved across the cohort (mean pairwise $r = 0.71 \pm 0.09$, positive for all 3655 pairs) and is unrelated to the stimulus. This was assessed with a distance-based PERMANOVA, the subject-by-subject distances $(1 - r)^2$ are partitioned into within- versus between-film dispersion to give a pseudo-$F$ ratio, whose null distribution is obtained by permuting the film labels (10,000 permutations), such that the only assumption is exchangeability under the null. No effect of film was found (pseudo-$F = 1.184$, $p_{\text{perm}} = 0.18$), and a subject-level paired contrast of each participant's mean similarity to same-film versus different-film participants returned a Bayes factor favouring the absence of an effect ($BF_{01} = 5.99$). (Bayes factors throughout are JZS Bayes factors computed from the paired $T$ statistic under a Cauchy prior of scale $r = 0.707$.)

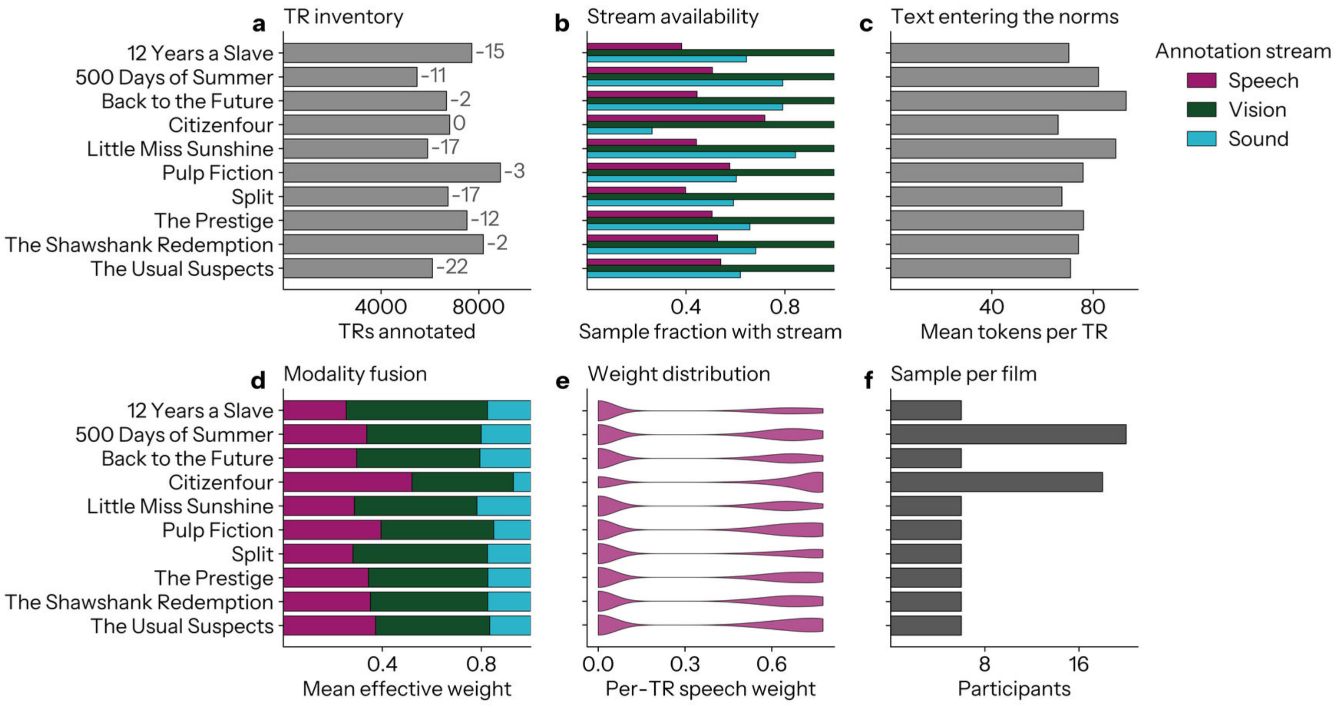


FIG. S13. **Quality control of the multimodal stimulus annotation.** Every TR of all ten films was described by an annotation of its speech, vision and sound, and the three streams fused into a single per-TR description (see Methods). Films are ordered alphabetically throughout. **a** Number of TRs annotated per film. The count appended to each bar is the number subsequently dropped as invalid, being those carrying an empty or degenerate caption, which removes title cards, black frames and other content-free intervals. Losses are negligible (101 of 69,982 TRs, 0.14%), and are written on rather than drawn because they are far too small to see. **b** Fraction of TRs at which each stream contributes any text. Vision is available at every TR of every film, whereas speech and sound are intermittent, so the visual stream carries the annotation wherever a film is quiet. **c** Mean number of tokens per TR entering the lexical norm scoring, pooled over the three streams. **d** Mean effective weight of each stream in the fused embedding, after the salience gate and the dialogue floor (see Methods). **e** Distribution of the per-TR speech weight within each film. The mass at zero is the set of TRs with no speech, at which the weight is reassigned to the remaining two streams. **f** Number of participants who watched each film.

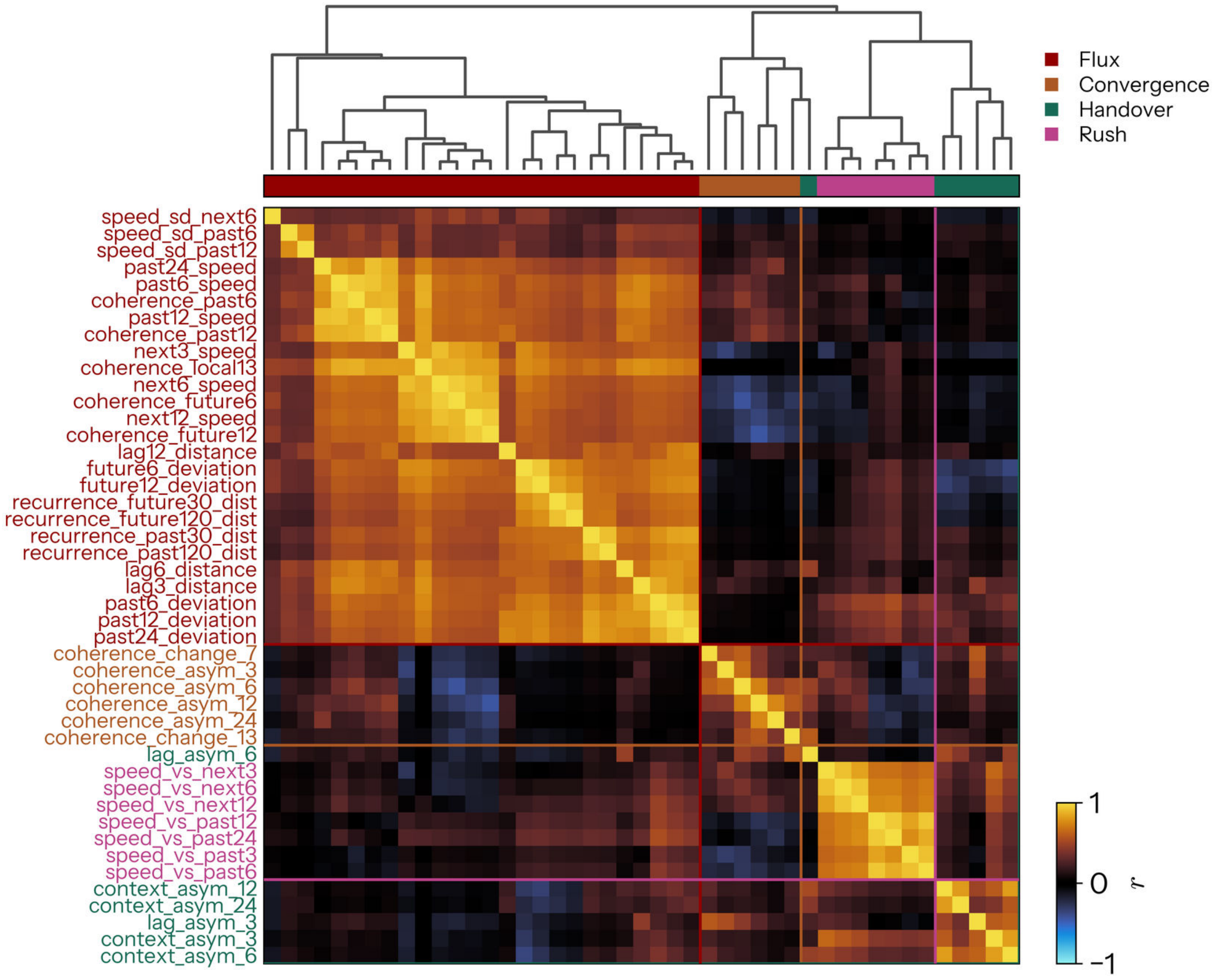


FIG. S14. **Kinematic measures recover their assigned constructs without supervision.** Correlation between every pair of the 45 trajectory measures, pooled across all films and computed after each measure has been oriented so that positive means more of its construct (Supplementary Table S1). Measures are ordered by average-linkage hierarchical clustering on $1 - r$, shown above, which is blind to the assignment. The colour strip beneath the dendrogram gives the family each measure was assigned to *a priori*, and coloured rules delimit the four clusters. Cutting the tree at four clusters recovers the hand assignment for 44 of the 45 measures, the single exception being `lag_asym_6`, which clusters with convergence rather than handover. The four blocks are internally coherent and largely independent of one another, so the axes are not simply four names for one quantity.

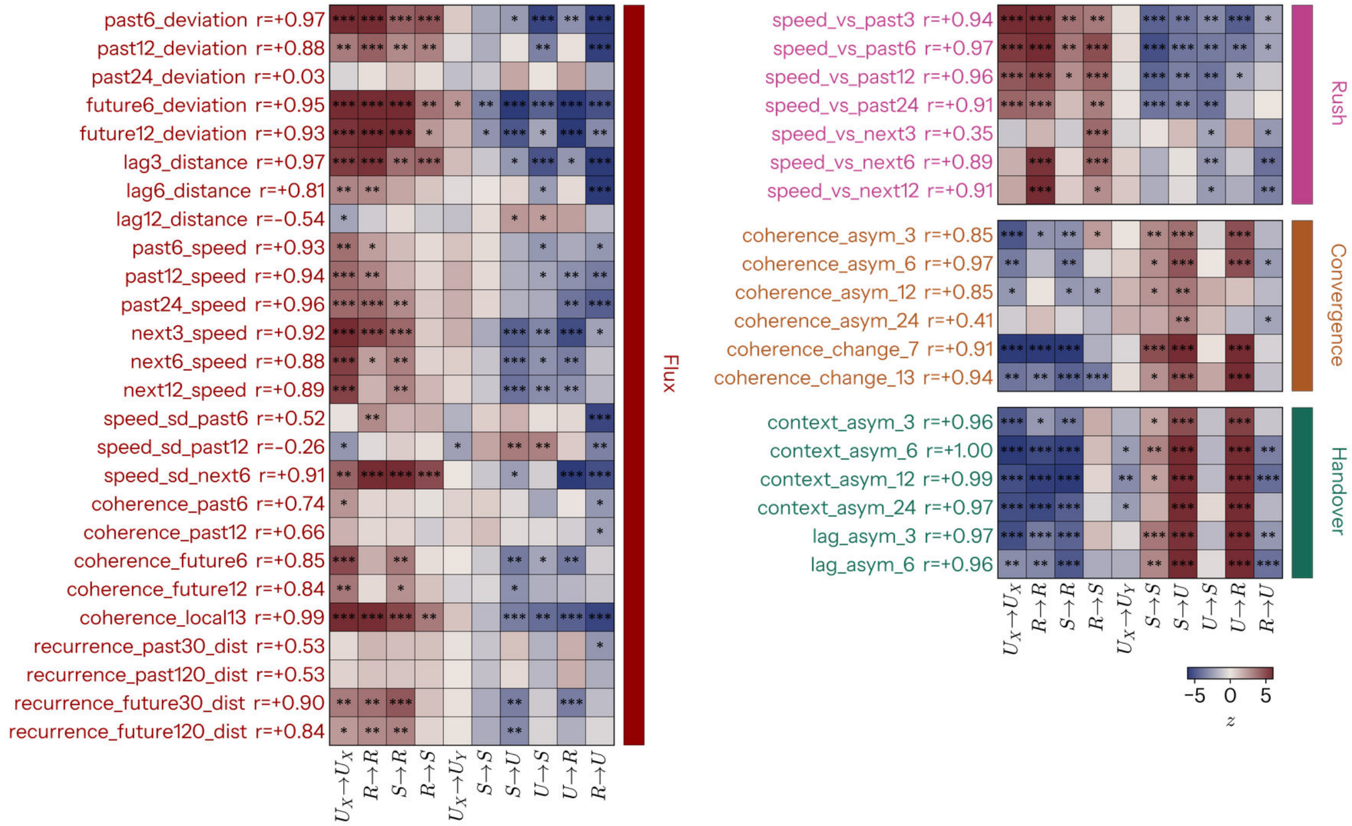


FIG. S15. **Replication with every kinematic measure individually** Because each axis in Fig. 6c is a mean over its family, its profile across modes could in principle be an artefact of averaging rather than a property of the construct. Each of the 45 measures was therefore tested individually against the same participant-wise circular-shift null, giving the $z$ shown here for every measure by mode cell. A measure cannot be assessed against a composite of which it is itself an ingredient, so the $r$ beside each measure name is its agreement with the *leave-one-out* composite of its family, that is the mean of the other members only, correlated across the ten modes (see Methods). Agreement is high throughout (mean $r = +0.71$ for flux, $+0.82$ for convergence, $+0.98$ for handover and $+0.85$ for rush), so the axes reflect their constructs rather than the particular measures chosen. Individual measures reproduce the profile of their axis, most visibly for handover, where every member separates $U \to R$ and $S \to U$ from $R \to R$, $S \to R$ and $U_X \to U_X$. Rows are grouped and coloured by family, columns are the ten modes ordered as in Fig. 6c. (*,**,*** indicates p<.05,.01,.001 FDR-corrected Benjamini-Hochberg procedure.)

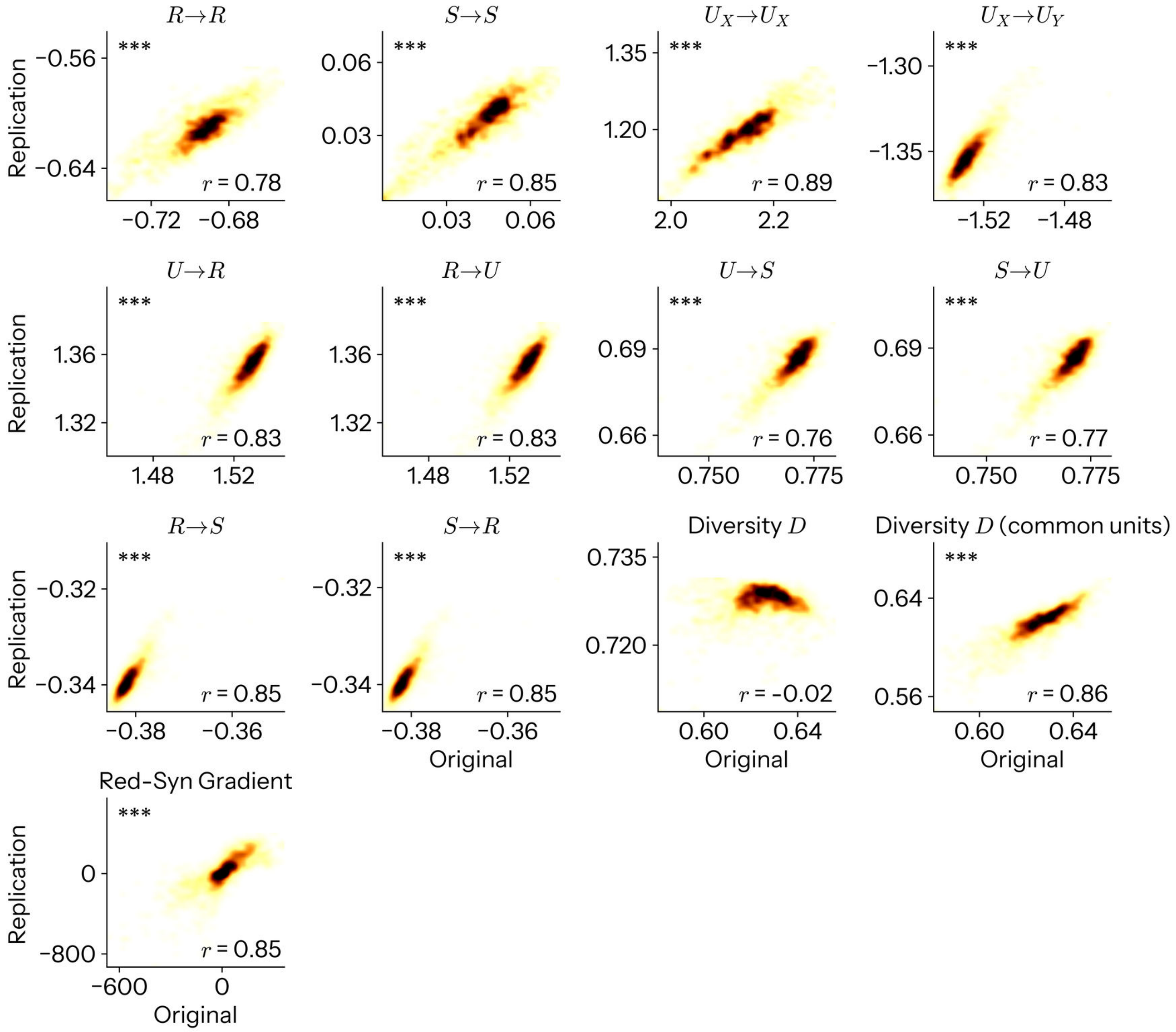


FIG. S16. **Replication of regional topographies.** Each panel is a two-dimensional histogram of one group-level regional map in the original cohort ($x$) against the same map in the replication cohort ($y$), over the 1054 regions of the atlas, smoothed for display. Panels give the ten modes of information dynamics, computational diversity $D$ on the raw atoms and on atoms placed in the original's units (*common units*), and the redundancy-to-synergy gradient. Sixteen maps were tested in all; the three representational complexity maps, which are near-identical to one another, are omitted from the figure and reported in Supplementary Note 2. The $r$ in each panel is the Spearman correlation between the two maps and the asterisks its significance against 1,000 SA-matched surrogates of the original map (uncorrected). Fifteen of the sixteen replicate at $p < 0.05$, with $r$ from $-0.022$ to $+0.894$ (median $+0.846$). The single exception is $D$ computed on the raw atoms, which is uncorrelated between cohorts ($r = -0.022$, $p_{\text{perm}} = 0.811$) and compressed into a narrow band, because the sampling interval rescales the sixteen atoms unequally and $D$ is a functional of their relative magnitudes; expressing both cohorts' atoms in common units restores it to $r = 0.859$, $p_{\text{perm}} < 10^{-3}$, at the permutation floor (Supplementary Note 2).

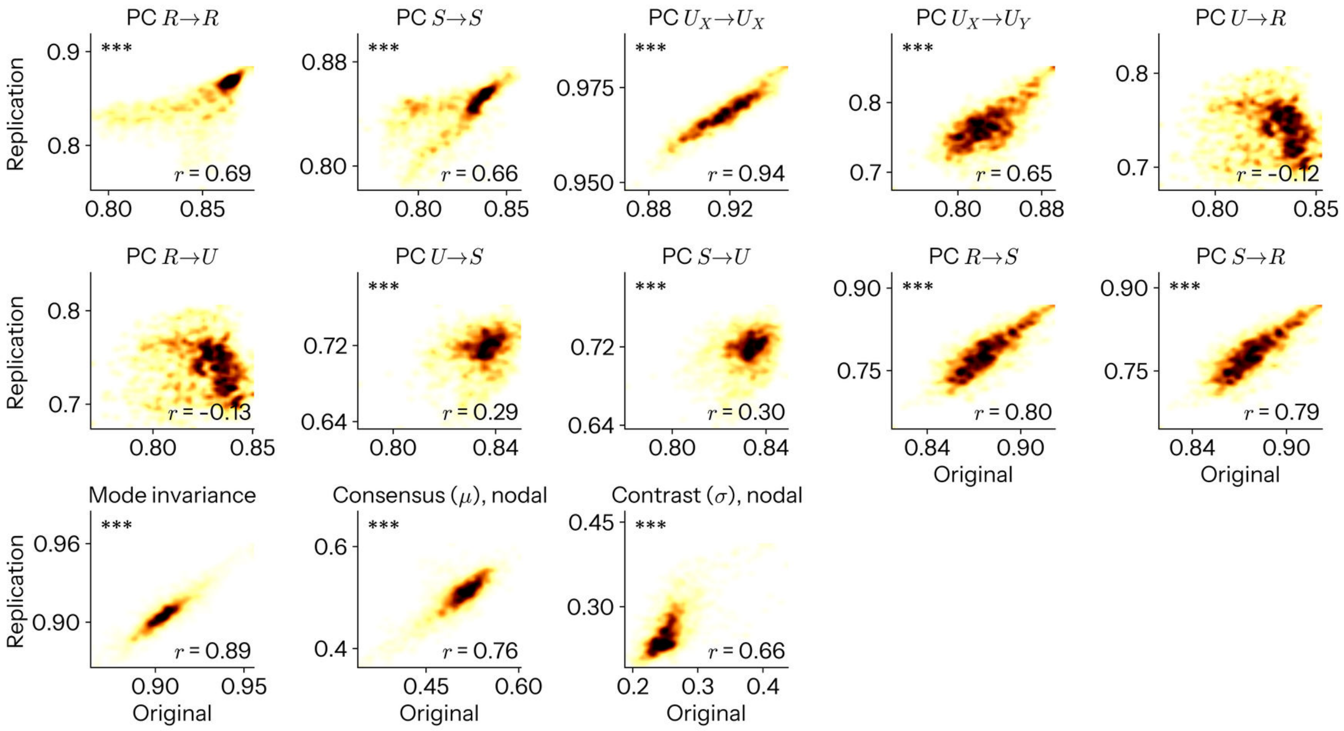


FIG. S17. **Replication of nodal network topographies.** As Supplementary Fig. S16, for the nodal quantities of Fig. 2: the participation coefficient (PC) of each mode's network, mode invariance, and the nodal means of the multiplex consensus $\mu$ and contrast $\sigma$. Eleven of the thirteen maps replicate at $p < 0.05$ SA-matched surrogates(median $r = +0.661$). The two that do not are the participation maps of the copy and erasure modes ($U \to R$ $r = -0.121$, $p_{\text{perm}} = 0.394$; $R \to U$ $r = -0.126$, $p_{\text{perm}} = 0.373$).

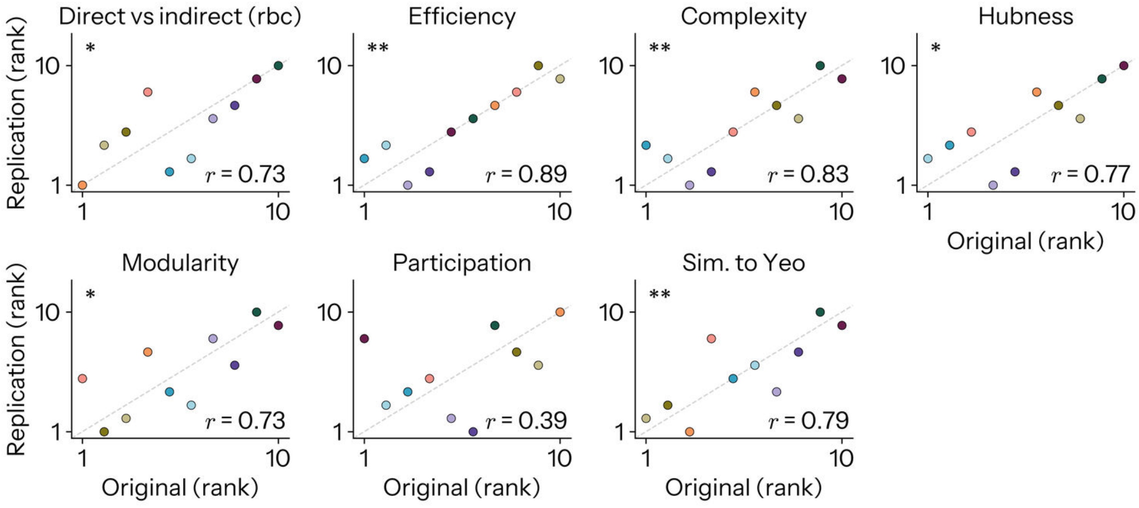


FIG. S18. **Replication of network profiles across modes.** Each panel plots one profile over the ten modes, original against replication, in ranks; points are modes, coloured as throughout, and the dashed line is the identity. Ten points are too few for a density, so these remain scatters. The $r$ is the Spearman correlation across modes and the asterisks its significance against 10,000 mode-relabelling permutations (uncorrected). Five of the six topological measures of Fig. 2d replicate (Efficiency $r = 0.891$, $p_{\text{perm}} = 0.001$; Complexity $r = 0.830$, 0.005; Hubness $r = 0.770$, 0.014; Modularity $r = 0.733$, 0.022; Sim. to Yeo $r = 0.794$, 0.009), as does the direct-against-indirect contrast of Fig. 2a as an ordering ($r = 0.733$, $p_{\text{perm}} = 0.023$), though only five of its ten signs agree. Participation does not replicate as an ordering ($r = 0.394$, $p_{\text{perm}} = 0.268$), notwithstanding that $U_X \to U_X$ is the most participatory mode in both cohorts.

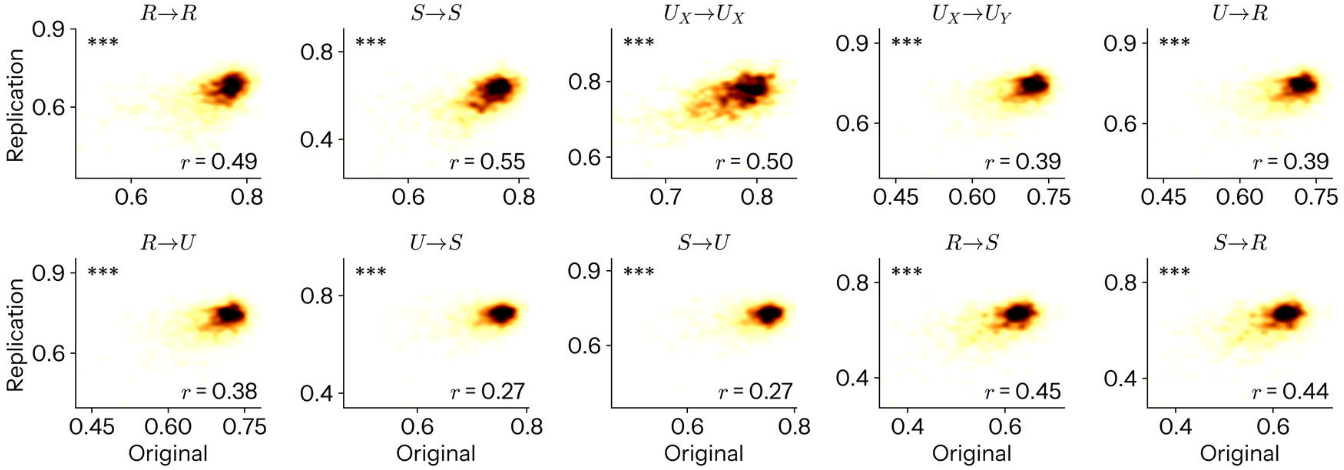


FIG. S19. **Replication of split-half reliability.** As Supplementary Fig. S16, for the node-averaged intra-class correlation of each mode (Fig. 3a). All ten maps replicate ($r = 0.267$ to $0.549$, all $p_{\text{perm}} < 10^{-3}$, the permutation floor of 1,000 SA-matched surrogates), as do the corresponding edge-wise ICC matrices when tested against the node-relabelling null ($r = 0.183$ to $0.468$, all $p_{\text{perm}} < 10^{-3}$, likewise at the floor; not shown).

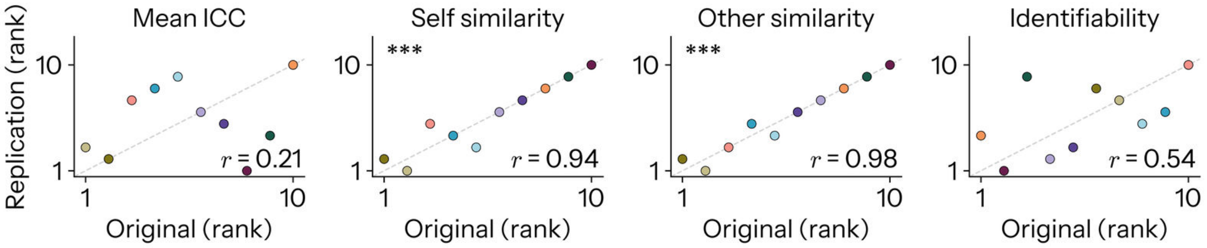


FIG. S20. **Replication of reliability and fingerprinting profiles across modes.** As Supplementary Fig. S18, for the mean ICC of each mode and the three fingerprinting quantities of Fig. 3b. Self and other similarity reproduce their orderings across modes almost exactly (self $r = 0.939$, $p_{\text{perm}} = 4.00 \times 10^{-4}$; other $r = 0.976$, $p_{\text{perm}} = 2.00 \times 10^{-4}$), whereas the ordering of modes by mean ICC does not ($r = 0.212$, $p_{\text{perm}} = 0.547$), nor by identifiability ($r = 0.539$, $p_{\text{perm}} = 0.114$). Identifiability does not differ across modes at all in the replication cohort ($F(9, 288) = 1.346$, $p = 0.213$), while remaining significant for 100% of participants in all ten modes, as in the original.

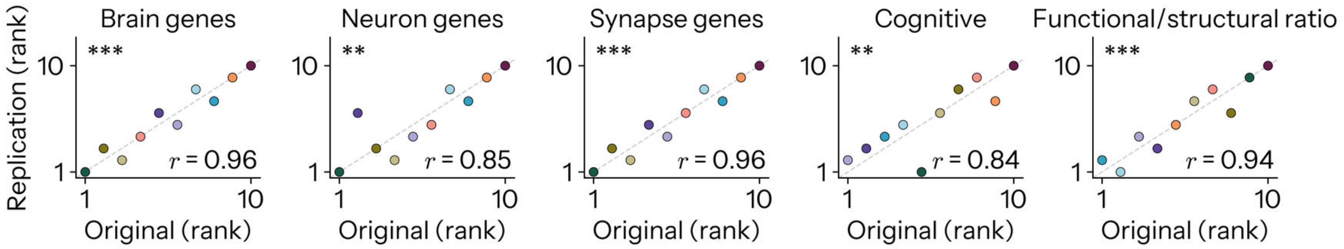


FIG. S21. **Replication of molecular and cognitive matching profiles across modes.** As Supplementary Fig. S18, for the matching profiles of Fig. 4c,d, the two alternative gene sets of Supplementary Fig. S8, and the functional-to-structural ratio. All five replicate (brain genes $r = 0.964$, $p_{\text{perm}} < 10^{-4}$, at the permutation floor of 10,000 nulls; neuron genes $r = 0.855$, 0.004; synapse genes $r = 0.964$, $< 10^{-4}$; cognitive $r = 0.842$, 0.004; functional/structural ratio $r = 0.939$, $5.00 \times 10^{-4}$), and in each the strongest mode is the same one, $S \to S$.

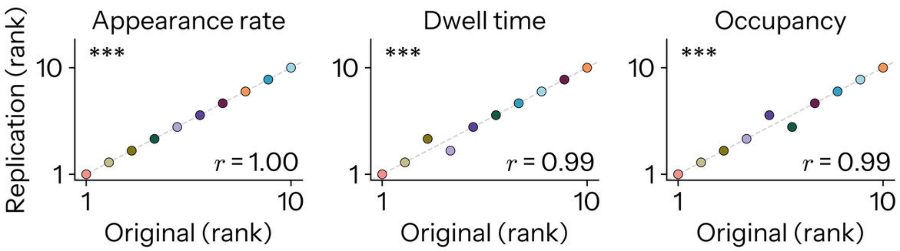


FIG. S22. **Replication of global state dynamics across modes.** As Supplementary Fig. S18, for the three temporal properties of the winner-take-all state sequence in Fig. 5b. All three orderings are recovered (appearance rate $r = 1.000$; dwell time $r = 0.988$; occupancy $r = 0.988$; all $p_{\mathrm{perm}} < 10^{-4}$ at the permutation floor of 10,000 mode relabellings), with $U_X \to U_X$ the longest-dwelling and most occupied mode and $U_X \to U_Y$ the rarest and most fleeting in both cohorts.

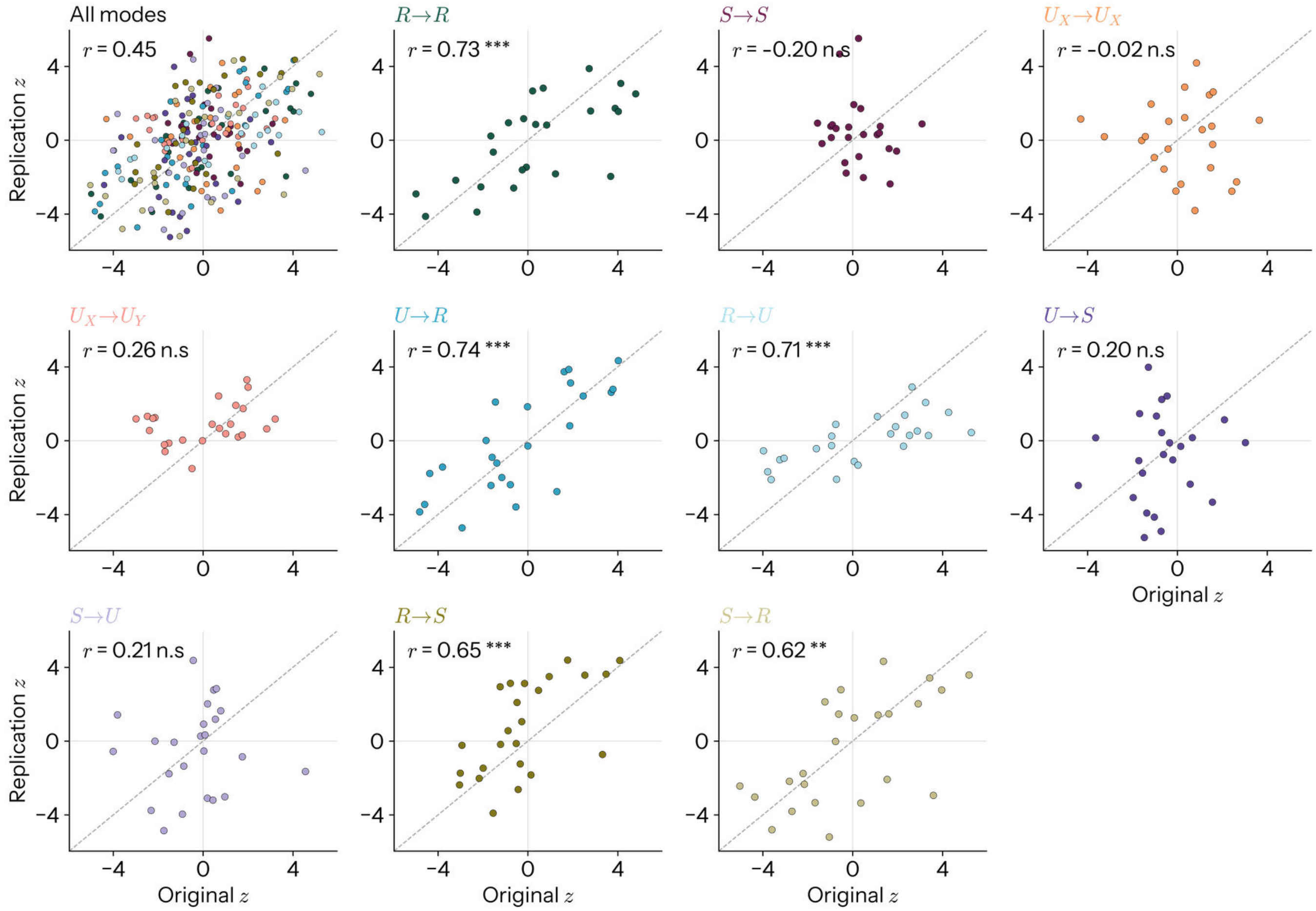


FIG. S23. **Replication of the normative profiles, mode by mode.** Each point is one mode by feature cell, plotted as its $z$ (against the participant-wise circular-shift null) in the original cohort against the same quantity in the replication NNDb-3T+ cohort. The dashed line is identity. Each 1.5s mode window is scored on the words of both 1s caption bins it straddles (Supplementary Note 2). **a** All mode by norm cells of Fig. 6a, coloured by mode. The $r$ is the Spearman correlation over all cells, tested against 10,000 permutations that relabel the modes of the original table, moving whole profiles at a time (uncorrected). **b–k** The same cells one mode at a time, 24 norms in each, with $r$ that mode's own profile agreement between the cohorts, tested against 10,000 permutations of the norms within that mode (uncorrected; the FDR-corrected values are given in Supplementary Note 2).

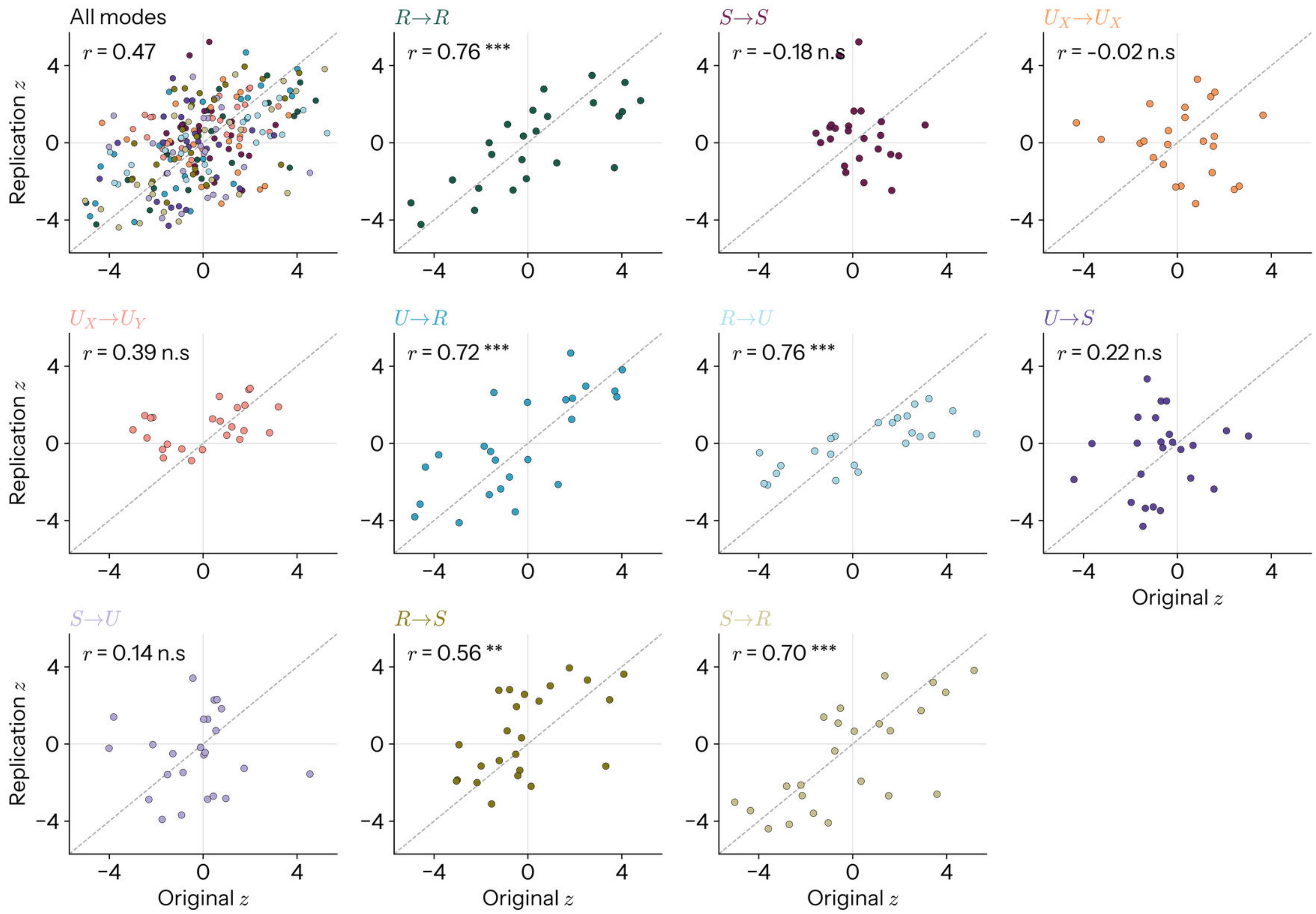


FIG. S24. **Replication of the normative profiles under the single-bin caption convention.** As Supplementary Fig. S23, with each 1.5s mode window scored only on the 1s caption bin it opens on, which is the convention of the original run, and here discards roughly a third of the film's words. The two conventions agree with one another at $r = 0.981$ over the 240 cells, and the same five profiles, and only those five, are recovered here as in Supplementary Fig. S23, so the division among the modes is not an artefact of how the caption bins are pooled.

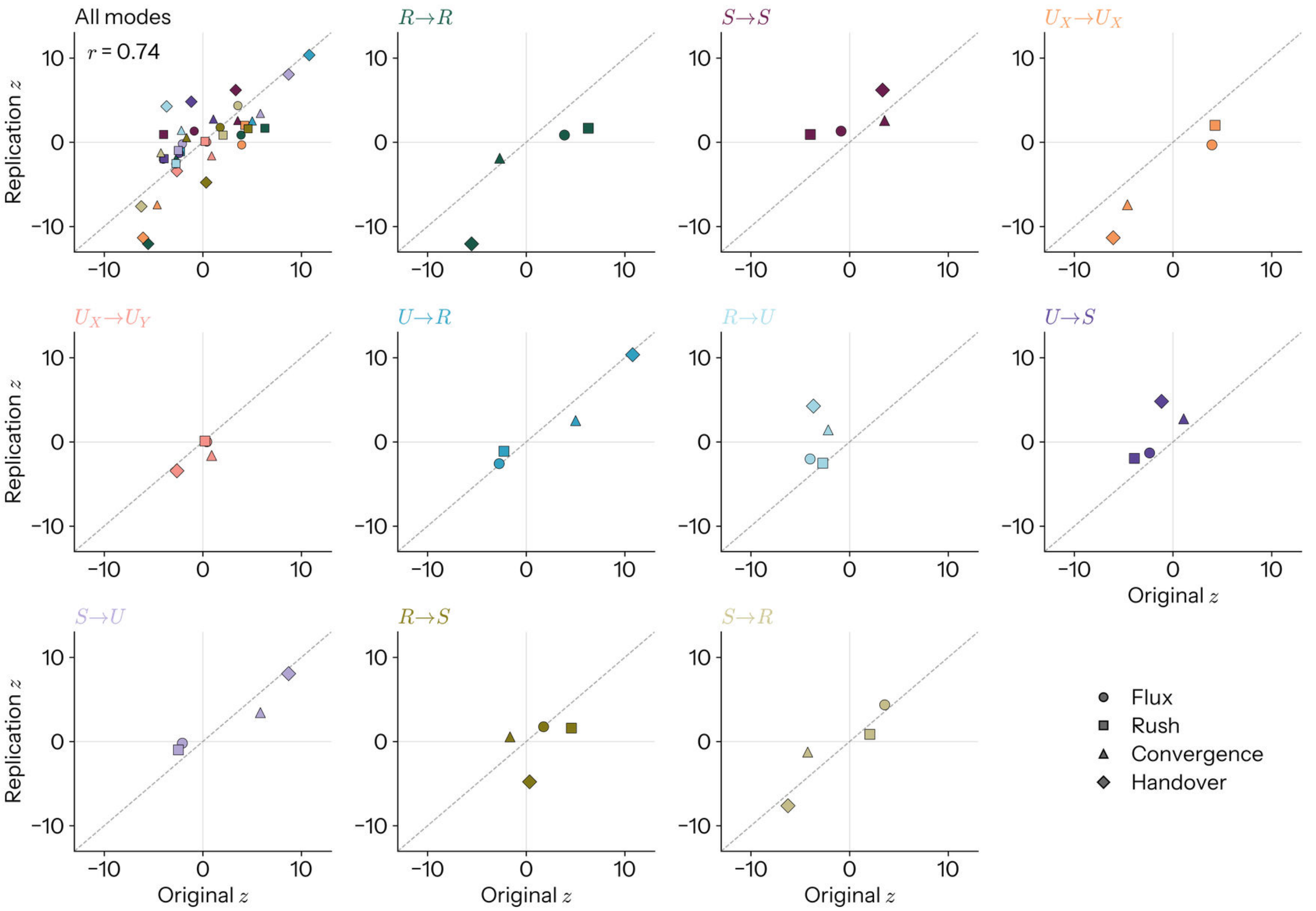


FIG. S25. **Replication of the semantic kinematics, mode by mode. a** All mode by axis cells of Fig. 6c, coloured by mode and marked by axis ($r = 0.737$, $p_{\text{perm}} = 9.00 \times 10^{-4}$, against the same mode-relabelling null as Supplementary Fig. S23). **b–k** The same cells one mode at a time. With only four points to a panel no per-mode correlation is reported; the agreement of each axis across the ten modes is given instead in Supplementary Note 2 (handover $r = 0.867$, flux $r = 0.636$, convergence $r = 0.842$, rush $r = 0.685$).

#### S6. SUPPLEMENTARY TABLES

| Measure | Sign | $r$ | Description |
|---|---|---|---|
| *Flux* (26 measures) | | | |
| past6_deviation | +1 | 0.65 | Distance from the mean of the previous 6 TRs. |
| past12_deviation | +1 | 0.65 | Distance from the mean of the previous 12 TRs. |
| past24_deviation | +1 | 0.63 | Distance from the mean of the previous 24 TRs. |
| future6_deviation | +1 | 0.63 | Distance from the mean of the next 6 TRs. |
| future12_deviation | +1 | 0.63 | Distance from the mean of the next 12 TRs. |
| lag3_distance | +1 | 0.63 | Distance to the state 3 TRs ago. |
| lag6_distance | +1 | 0.62 | Distance to the state 6 TRs ago. |
| lag12_distance | +1 | 0.55 | Distance to the state 12 TRs ago. |
| past6_speed | +1 | 0.64 | Mean step size over the previous 6 TRs. |
| past12_speed | +1 | 0.63 | Mean step size over the previous 12 TRs. |
| past24_speed | +1 | 0.57 | Mean step size over the previous 24 TRs. |
| next3_speed | +1 | 0.62 | Mean step size over the next 3 TRs. |
| next6_speed | +1 | 0.63 | Mean step size over the next 6 TRs. |
| next12_speed | +1 | 0.61 | Mean step size over the next 12 TRs. |
| speed_sd_past6 | +1 | 0.38 | SD of step size over the previous 6 TRs. |
| speed_sd_past12 | +1 | 0.37 | SD of step size over the previous 12 TRs. |
| speed_sd_next6 | +1 | 0.34 | SD of step size over the next 6 TRs. |
| coherence_past6 | −1 | 0.64 | Coherence of the previous 6 states. |
| coherence_past12 | −1 | 0.63 | Coherence of the previous 12 states. |
| coherence_future6 | −1 | 0.62 | Coherence of the next 6 states. |
| coherence_future12 | −1 | 0.59 | Coherence of the next 12 states. |
| coherence_local13 | −1 | 0.72 | Coherence of the 13 states centred on this TR. |
| recurrence_past30_dist | +1 | 0.60 | Distance to the nearest match 4-30 TRs back. |
| recurrence_past120_dist | +1 | 0.56 | Distance to the nearest match 4-120 TRs back. |
| recurrence_future30_dist | +1 | 0.58 | Distance to the nearest match 4-30 TRs ahead. |
| recurrence_future120_dist | +1 | 0.55 | Distance to the nearest match 4-120 TRs ahead. |
| *Rush* (7 measures) | | | |
| speed_vs_past3 | +1 | 0.75 | Current step size minus the previous 3-TR mean. |
| speed_vs_past6 | +1 | 0.80 | Current step size minus the previous 6-TR mean. |
| speed_vs_past12 | +1 | 0.81 | Current step size minus the previous 12-TR mean. |
| speed_vs_past24 | +1 | 0.79 | Current step size minus the previous 24-TR mean. |
| speed_vs_next3 | +1 | 0.73 | Current step size minus the next 3-TR mean. |
| speed_vs_next6 | +1 | 0.78 | Current step size minus the next 6-TR mean. |
| speed_vs_next12 | +1 | 0.78 | Current step size minus the next 12-TR mean. |
| *Convergence* (6 measures) | | | |
| coherence_asym_3 | +1 | 0.41 | Future-3 coherence minus past-3 coherence. |
| coherence_asym_6 | +1 | 0.60 | Future-6 coherence minus past-6 coherence. |
| coherence_asym_12 | +1 | 0.58 | Future-12 coherence minus past-12 coherence. |
| coherence_asym_24 | +1 | 0.42 | Future-24 coherence minus past-24 coherence. |
| coherence_change_7 | +1 | 0.41 | Change in centred 7-TR coherence from the prior TR. |
| coherence_change_13 | +1 | 0.38 | Change in centred 13-TR coherence from the prior TR. |
| *Handover* (6 measures) | | | |
| context_asym_3 | −1 | 0.51 | Future-3 deviation minus past-3 deviation. |
| context_asym_6 | −1 | 0.66 | Future-6 deviation minus past-6 deviation. |
| context_asym_12 | −1 | 0.65 | Future-12 deviation minus past-12 deviation. |
| context_asym_24 | −1 | 0.51 | Future-24 deviation minus past-24 deviation. |
| lag_asym_3 | −1 | 0.46 | Distance to the state 3 TRs ahead minus 3 TRs back. |
| lag_asym_6 | −1 | 0.36 | Distance to the state 6 TRs ahead minus 6 TRs back. |

TABLE S1. **The bank of semantic kinematic measures.** The 45 measures of the stimulus trajectory through semantic space, grouped by the construct each operationalises. *Sign* is the orientation applied before averaging, +1 where the measure already increases with the construct and −1 where it runs opposite to it (coherence, for instance, is high when meaning is *not* moving, and so enters flux negatively). $r$ is each measure's correlation with the mean of the other members of its family, pooled over all films, and indexes how well it stands in for the construct. All measures are computed on the raw per-TR embedding using cosine distance, and $z$-scored within film. TR = 1 s throughout.

[1] Luppi, A. I. *et al.* A synergistic core for human brain evolution and cognition. *Nat. Neurosci.* **25**, 771–782 (2022).
[2] Wang, L., Mruczek, R. E. B., Arcaro, M. J. & Kastner, S. Probabilistic maps of visual topography in human cortex. *Cereb. Cortex* **25**, 3911–3931 (2015).
[3] Yeo, B. T. T. *et al.* The organization of the human cerebral cortex estimated by intrinsic functional connectivity. *J. Neurophysiol.* **106**, 1125–1165 (2011).
[4] Mesulam, M. M. Large-scale neurocognitive networks and distributed processing for attention, language, and memory. *Ann. Neurol.* **28**, 597–613 (1990).
[5] Ehrlich, D. A., Schneider, A. C., Priesemann, V., Wibral, M. & Makkeh, A. A measure of the complexity of neural representations based on partial information decomposition. *arXiv [cs.IT]* (2022). 2209.10438.
[6] Levchenko, E. *et al.* A neuroimaging dataset combining movie-watching, eye-tracking, sensorimotor mapping, and cognitive tasks. *Sci. Data* 1–19 (2026).
[7] Burt, J. B., Helmer, M., Shinn, M., Anticevic, A. & Murray, J. D. Generative modeling of brain maps with spatial autocorrelation. *Neuroimage* **220**, 117038 (2020).
[8] Mediano, P. A. M. *et al.* Toward a unified taxonomy of information dynamics via integrated information decomposition. *Proc. Natl. Acad. Sci. U. S. A.* **122**, e2423297122 (2025).
[9] Liardi, A., Blackburne, G. & Mediano, P. A. M. An information-geometric formulation of distributed computation. *arXiv [cs.IT]* (2026). 2611.03545.
[10] Kay, J. W. A partial information decomposition for multivariate gaussian systems based on information geometry. *Entropy (Basel)* **26**, 542 (2024).
[11] Varley, T. F., Pope, M., Puxeddu, M. G., Faskowitz, J. & Sporns, O. Partial entropy decomposition reveals higher-order information structures in human brain activity. *Proc. Natl. Acad. Sci. U. S. A.* **120**, e2300888120 (2023).
[12] Cahoy, J. D. *et al.* A transcriptome database for astrocytes, neurons, and oligodendrocytes: a new resource for understanding brain development and function. *J. Neurosci.* **28**, 264–278 (2008).
[13] Genovese, G. *et al.* Increased burden of ultra-rare protein-altering variants among 4,877 individuals with schizophrenia. *Nat. Neurosci.* **19**, 1433–1441 (2016).
[14] Pirooznia, M. *et al.* SynaptomeDB: an ontology-based knowledgebase for synaptic genes. *Bioinformatics* **28**, 897–899 (2012).
[15] van Es, M. W. J. *et al.* Large-scale cortical functional networks are organized in structured cycles. *Nat. Neurosci.* 1–11 (2025).
